# Population genetic analysis reveals the role of natural selection and phylogeography on genome-wide diversity in an extremely compact and reduced microsporidian genome

**DOI:** 10.1101/2022.03.29.486185

**Authors:** Pascal Angst, Dieter Ebert, Peter D. Fields

## Abstract

The determinants of variation in a species’ genome-wide nucleotide diversity include historical, environmental, and stochastic aspects. This diversity can inform us about the species’ past and present evolutionary dynamics. In parasites, the mode of transmission and the interactions with the host might supersede the effects of these aspects in shaping parasite genomic diversity. We used genomic samples from ten populations of the microsporidian parasite *Ordospora colligata* to investigate present genomic diversity and how it was shaped by evolutionary processes, specifically, the role of phylogeography, co-phylogeography (with the host), natural selection, and transmission mode. Although very closely related microsporidia cause diseases in humans, *O. colligata* is specific to the freshwater crustacean *Daphnia magna* and has one of the smallest known eukaryotic genomes. We found an overlapping phylogeography between *O. colligata* and its host highlighting the long-term, intimate relationship between them. The observed geographic distribution reflects previous findings that *O. colligata* exhibits adaptations to colder habitats, which differentiates it from other microsporidian gut parasites of *D. magna* predominantly found in warmer areas. The co-phylogeography allowed us to calibrate the *O. colligata* phylogeny and thus estimate its mutation rate. We found patterns of more efficient purifying selection in *O. colligata* relative to other microsporidia sharing the same host, which likely allowed this parasite to maintain its very compact genome. We also identified regions under potential selection related to coevolution including the ribosomal protein L24, a leucyl-tRNA synthetase, and a putative ABC-like lipid transport protein. Our whole-genome study provides insights into the evolution of one of the most reduced eukaryotic genomes and shows how different processes shape genomic diversity of an obligate parasite.

**Author summary:** Microsporidia are intracellular parasites that infect vertebrates, invertebrates, and even unicellular organisms. Due to their high variation in many aspects of life history and genomics, microsporidia have become a model clade for understanding evolutionary processes related to intracellular parasitism. However, the evolution of extreme genomic architectures in microsporidia and the coevolution with their hosts is still under-surveyed, especially given their role in human disease. Here, we study past and present evolutionary dynamics in a microsporidian species with one of the smallest known eukaryotic genomes, *O. colligata*. Close relatives of *O. colligata* cause death and disease in humans and agriculturally important animals. We show that purifying selection helped maintaining its reduced, compact genome and corroborate hypotheses about the evolution of different genome sizes in microsporidia. Importantly, we utilize the highly resolved phylogeny of its host to estimate the parasite’s mutation rate. This methodology allowed us to establish the first mutation rate estimate for a microsporidium, an estimate which is within the range of mutation rates estimated for phylogenetically related, non-parasitic fungi. Our study exemplifies how the combined knowledge about a species’ biology, ecology, and genomic diversity helps to resolve its evolutionary dynamics, in particular when phylogenomic information can be brought to bear for both host and parasite.

## Introduction

Understanding a species’ genome-wide nucleotide diversity requires information about historical conditions (e.g., phylogeography and population structure), environmental conditions (e.g., adaptation to local climate), and stochastic processes (e.g., genetic drift and founder events) [1–3]. The contribution of each factor to the observed genomic diversity is highly variable. For example, in species prone to small population sizes, alleles may reach fixation solely due to genetic drift [4]. In contrast, in larger populations, natural selection is more efficient in preventing deleterious mutations from being fixed and in deterministically increasing the frequency of beneficial alleles. For parasites, hosts are part of the environment, with interactions between host and parasite possibly having profound impact on the parasite’s genomic diversity [5]. A distinct feature of parasite life history is its mode of transmission. A parasite can be transmitted vertically (from parent to offspring), horizontally (no parent-progeny relationship between hosts) or mixed-mode (featuring vertical and horizontal transmission) [6]. The genetic diversity of specific regions under selection, or genome-wide allele frequencies might change due to the parasite’s mode of transmission [7]. Notably, the acquisition of vertical transmission by horizontally transmitted parasites was suggested to increase the potential for stochastic processes to produce random changes in the genome, because with vertical transmission population bottlenecks become more likely and thus effective population sizes, *N*_*e*_, would be reduced [8]. Disentangling the effects of parasite-specific life histories from more general processes on a genomic level is still a challenge in the field of evolutionary genomics.

Microsporidia are a clade of intracellular parasites that is characterized by high variation in many aspects of life history and genomics. They cause diseases commonly referred to as microsporidiosis in agriculturally important animals, honeybees, and immunocompromised humans, among others [9,10] and are reported to infect even unicellular organisms [11]. A growing body of research features them as a model clade for understanding evolutionary processes related to extreme parasitism [10,11]. Microsporidia are phylogenetically associated with the fungi, but show high specialization to intracellular life-style combined with a large variation in life histories [12]. Most microsporidia do not have mitochondria but take up energy from their host using transmembrane transporters, although a few species have maintained mitochondria (e.g., [13]). Their genome size (about two to 50 Megabase pairs) and number of genes vary extremely [10], but microsporidia generally have a small number of genes and small genome size, including one of the smallest known eukaryotic genomes [14]. Recent work suggests that large genomes evolved in microsporidia due to the accumulation of repetitive elements when the strength of purifying selection decreases [15]. For example, large, gene-sparse genomes evolved in the microsporidian genera *Hamiltosporidium* and *Nosema* after a switch from horizontal transmission to mixed-mode transmission, which reduced *N*_*e*_ [8]. However, only ∼18 % of the species are potentially vertically transmitted [11], while the more common exclusive horizontal transmission should allow for large *N*_*e*_ and the maintenance of the typical streamlined, i.e., reduced and compact, genomes.

Hosts of several microsporidia are planktonic freshwater crustacea of the genus *Daphnia*, well-established model systems in ecology, evolution, and in the study of host-parasite interactions [16–18]. Several microsporidia that infect *D. magna* have been studied both ecologically, as well as on the genomic level, which is important when ecological scenarios are used to explain genome evolution. Examples include the microsporidium *Hamiltosporidium tvaerminnensis*, a microsporidium with a large genome size and mixed-mode of transmission [19–22], and several microsporidian gut parasites – e.g. *Glugoides intestinalis, Mitosporidium daphniae*, and *Ordospora colligata* – that have small genomes, look superficially similar under the microscope, infect the same host tissue, and are of relatively low virulence to the host [13,23,24]. However, the three mentioned *D. magna* gut microsporidia diverge substantially on the molecular level [25]. *O. colligata* has received a lot of attention because it has one of the most reduced genomes across all eukaryotes and is closely related to the microsporidian genus *Encephalitozoon* that primarily infects humans and other mammals [14]. This species features in several ecological and evolutionary studies (e.g., [26–29]), some of which suggest the species is adapted to colder habitats [30]. Compared to, for example, *H. tvaerminnensis, O. colligata* is suggested to exhibit large population sizes that may be a key factor for maintaining its streamlined genome [8]. Importantly, *O. colligata* is transmitted entirely horizontally [31], and little is known about the evolution of exclusively horizontally transmitted microsporidian parasites.

In this population genomic study, we characterize genomic diversity across the species range of *O. colligata*. We aimed to understand the relative contribution of phylogeography, selection, and mode of transmission to shaping the variation in nucleotide diversity. The phylogeography of *D. magna*, the only known host of *O. colligata*, shows three main lineages and isolation-by-distance (IBD) [32–35]. Based on neutral assumptions, we expected a shared phylogeography between host and obligate parasite, i.e., co-cladogenesis. Furthermore, because of *O. colligata*’s horizontal mode of transmission and its streamlined genome, we expected to find a constant, large population size, allowing for efficient selection. We wanted to use sequentially Markovian coalescent analyses and McDonald-Kreitman tests to estimate the species’ effective population size history and rate of adaptive evolution, respectively. We expected to find a higher selection efficacy in *O. colligata* than previously published in *Hamiltosporidium*, which would support the hypothesis by Haag et al. [8] that the mode of transmission is a driving factor for the evolution of genome size in microsporidia. Specifically, horizontal transmission is expected to go hand in hand with large *N*_*e*_ and the maintenance of a streamlined genome [8]. At least in part, our population genomic study corroborates earlier speculation about mechanisms driving the evolution of strongly diverged genome sizes in microsporidia by investigating the relative importance and interplay of biologically meaningful processes shaping the variation in genomic diversity.

## Results

### Samples, mapping, and sequence variation

The accuracy of microscopy in identifying *D. magna* microsporidian gut parasites on a species level was not as high as we had assumed. By using the combination of microscopy and PCR, we could reliably identify species, and we found that *D. magna* shows infections with microsporidian gut parasites across its Holarctic species range (Fig 1, S1 Table). *O. colligata* showed the widest geographic distribution but was absent from southern regions. Genomic samples of ten sequenced host clones sufficiently represented the *O. colligata* genome to describe its variation in genomic diversity (Table 1). The percentage of sequencing reads, originating from the combined sequencing of host and parasite, being mapped to the parasite’s genome ranged from 0.22 to 2.97 % for *O. colligata*. The average whole-genome coverage for *O. colligata* was greater than 10× in all samples. *O. colligata* has a SNP density of approximately five SNPs per Kbp and a total number of 12,427 SNPs. The average whole-genome estimate of π based on one Kbp windows was 0.003, approximately half a percent.

**Table 1.** Sample information.

| Sample ID | Country | Latitude | Longitude | Coverage |
| --- | --- | --- | --- | --- |
| CN-WON-2 | China | 46.825017 | 124.378953 | 10× |
| FI-SK-17-1 | Finland | 59.8312191 | 23.2577045 | 19× |
| FI-SKW-2-1 | Finland | 59.8329692 | 23.2562571 | 26× |
| GB-EP-1 | Great Britain | 52.4535830 | -1.9194440 | 71× |
| NO-V-7 | Norway | 67.68696 | 12.67197 | 70× |
| RU-BAYA1-1 | Russia | 53.017833 | 106.886833 | 29× |
| RU-KU1-2 | Russia | 55.303833 | 63.405333 | 21× |
| US-SP131-1 | United States | 44.3332 | -68.0632 | 40× |
| US-SP15-1 | United States | 44.3331 | -68.0629 | 55× |
| US-SP163-1 | United States | 44.3336 | -68.0636 | 37× |

*O. colligata*'s average whole-genome coverage is denoted in the last column.

**Fig 1.**
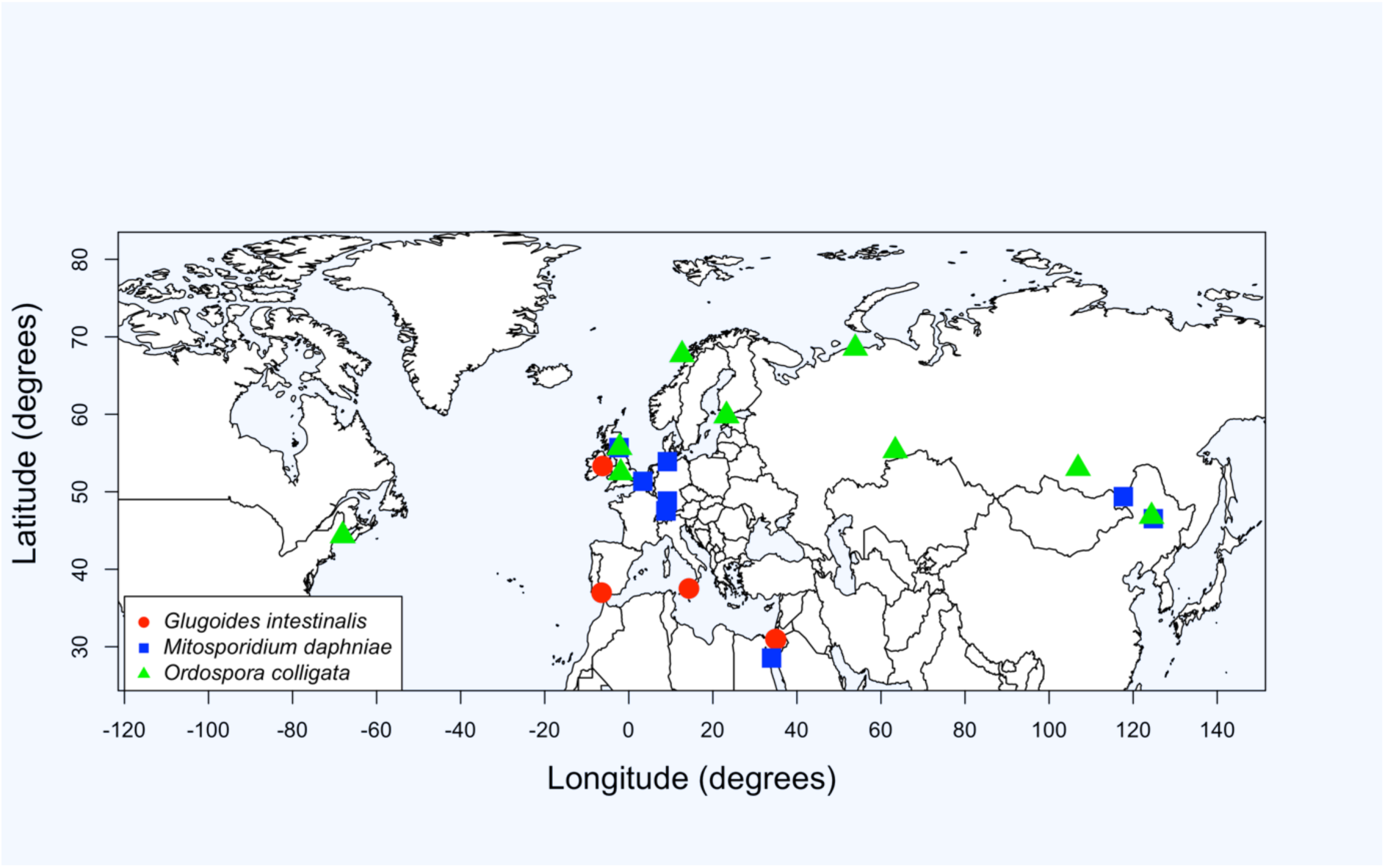
Sampling distribution of three *D. magna* gut microsporidia. Based on sampling within the framework of the *D. magna* diversity panel, we found wide-range distributions of the host species-specific microsporidia *Glugoides intestinalis* (red circles), *Mitosporidium daphniae* (blue squares), and *Ordospora colligata* (green triangles). *O. colligata* has the widest distribution within the Holarctic host-distribution, while the other two microsporidia have not been observed in Northern America.

The *O. colligata* genome has at least three regions that were horizontally transmitted from the host [14]. Windows overlapping with these regions showed increased diversity due to unintentional mapping of *D. magna* reads (Fig 2). Other windows within the 99th percentiles for π overlapped with functionally annotated genes (locus IDs as adopted from Pombert et al. [14]: the ribosomal protein L24 (M896_051540), a leucyl-tRNA synthetase (M896_060290) and a putative ABC-like lipid transport protein (M896_121220)), of which a putative ABC-like lipid transport protein additionally had the fourth highest π_N_/π_S_ ratio, i.e., it might experience strong positive or balancing selection (S2 Table). The remaining windows within the 99th percentiles for π did not overlap with annotated genes.

**Fig 2.**
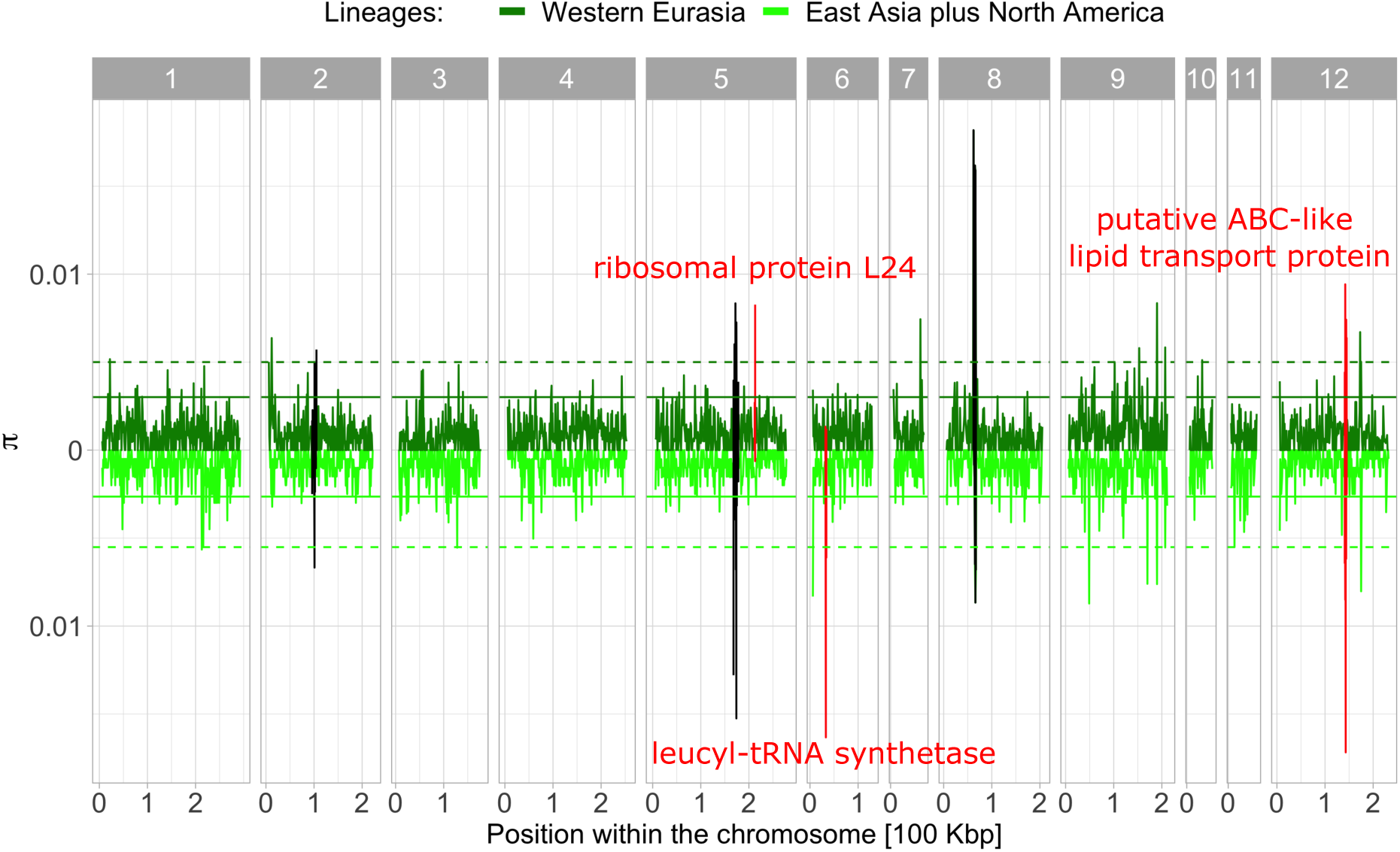
Average per-site nucleotide differences (π) in *O. colligata*. Per-site π is averaged over 1 Kbp windows along the genome. The π values of both main *O. colligata* lineages are presented in 12 facets corresponding to the parasite’s 12 chromosomes. We used the combined sample of East Asian and North American samples to have equal sample sizes for comparing lineages (N = 5). The 95th and 99th percentiles of each lineage are indicated with a solid and a dashed line, respectively. The first and last five Kbp per chromosome were masked for better presentation. Black windows indicate increased diversity due to host-to-parasite horizontal gene transfer. Annotated genes with increased π values mentioned in the discussion are indicated in red with their locus IDs: the ribosomal protein L24 (M896_051540), a leucyl-tRNA synthetase (M896_060290) and a putative ABC-like lipid transport protein (M896_121220).

### Ploidy

Allele frequency signatures of *O. colligata* samples showed haploid characteristics in 7 out of 9 samples (Fig 3, S1 and S2 Figs). The other two were not as clear due to low levels of polymorphisms or potential multiple infections of *O. colligata*. Specifically, no prominent peaks were found in the putative allele frequency histogram calculated using ploidyNGS. One peak can be seen in the k-mer histogram around the average whole-genome coverage. Therefore, the trimmed sequencing reads supported a single variant for most positions along the genome and no signs of heterozygosity were visible, hence *O. colligata* was treated as haploid in all analyses.

**Fig 3.**
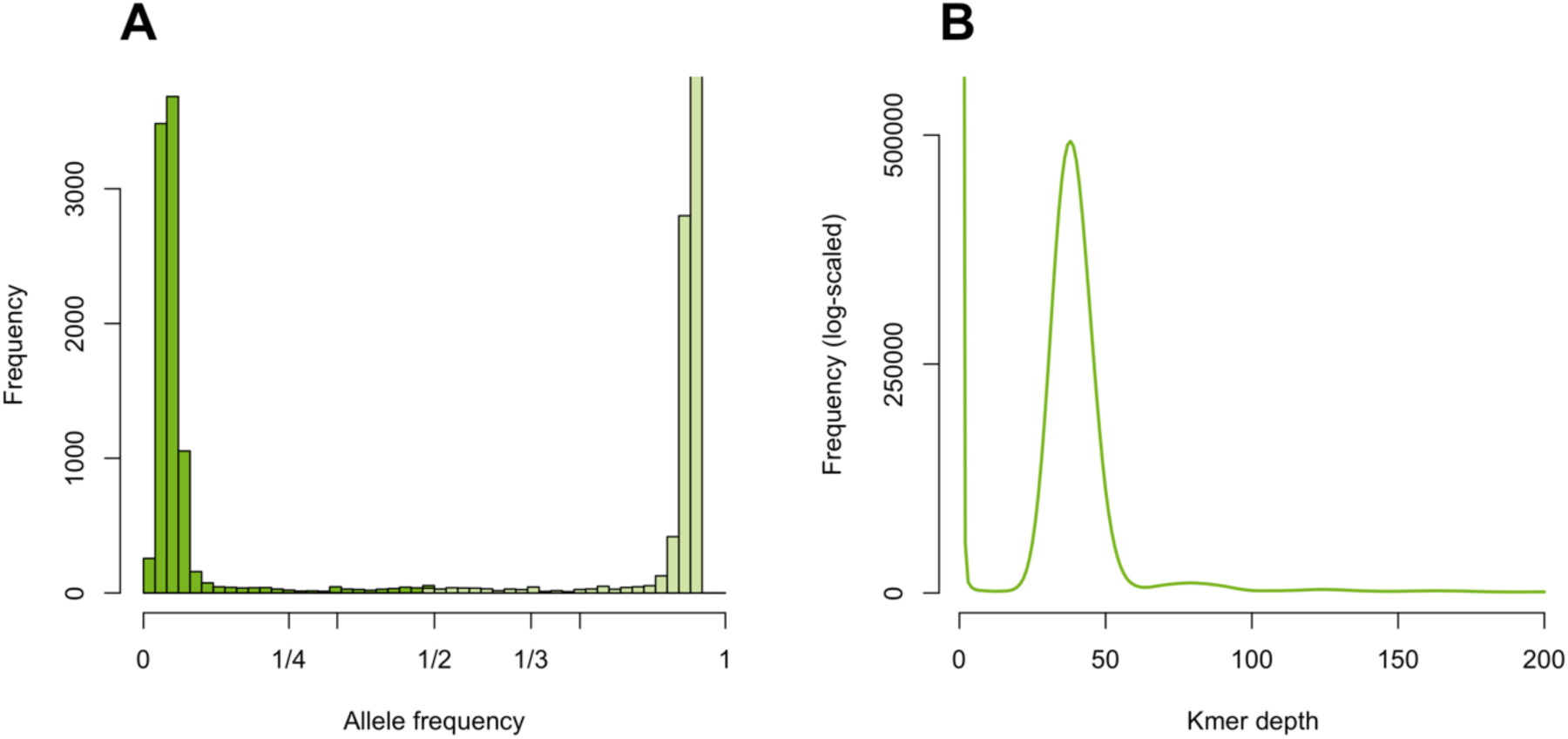
Graphical summaries of the two approaches to determine ploidy levels for *O. colligata* sample US-SP15-1. Histogram of allele frequency of putative alleles in the BAM file found using ploidyNGS (A), where reads supported a single variant for most positions. Few sites and no sharp central peak at intermediate allele frequencies is seen as would be the case in diploids. Dark and light shadings of green indicate reference and alternative allele, respectively. K-mer frequency histogram inferred using kmercountexact.sh (B), with a single peak for homozygous sites and no secondary peaks which would indicate di-/polyploidy.

### Population structure

To determine the shared or differing amounts of genomic variation between *O. colligata* samples, we used PCA on the whole-genome SNP data. PC1 separated samples in three clusters, which became even clearer in combination with PC2 (S3 Fig). The two PCs explained a substantial amount of genomic variation, with 63.60 % and 15.43 %, respectively. The three clusters (Western Eurasian (N = 5), East Asian (N = 2), and North American (N = 3) cluster, named according to the geographic regions the samples in each cluster originate from) were also found in the cluster analysis and the Bayesian phylogenetic tree estimation based on four-fold degenerate sites (Fig 4, S4 and S5 Figs). Pairwise relatedness in *O. colligata* was negatively correlated with geographic distance (Fig 5; *R*^*2*^ = 0.54, *p* = 0.001), suggesting IBD.

**Fig 4.**
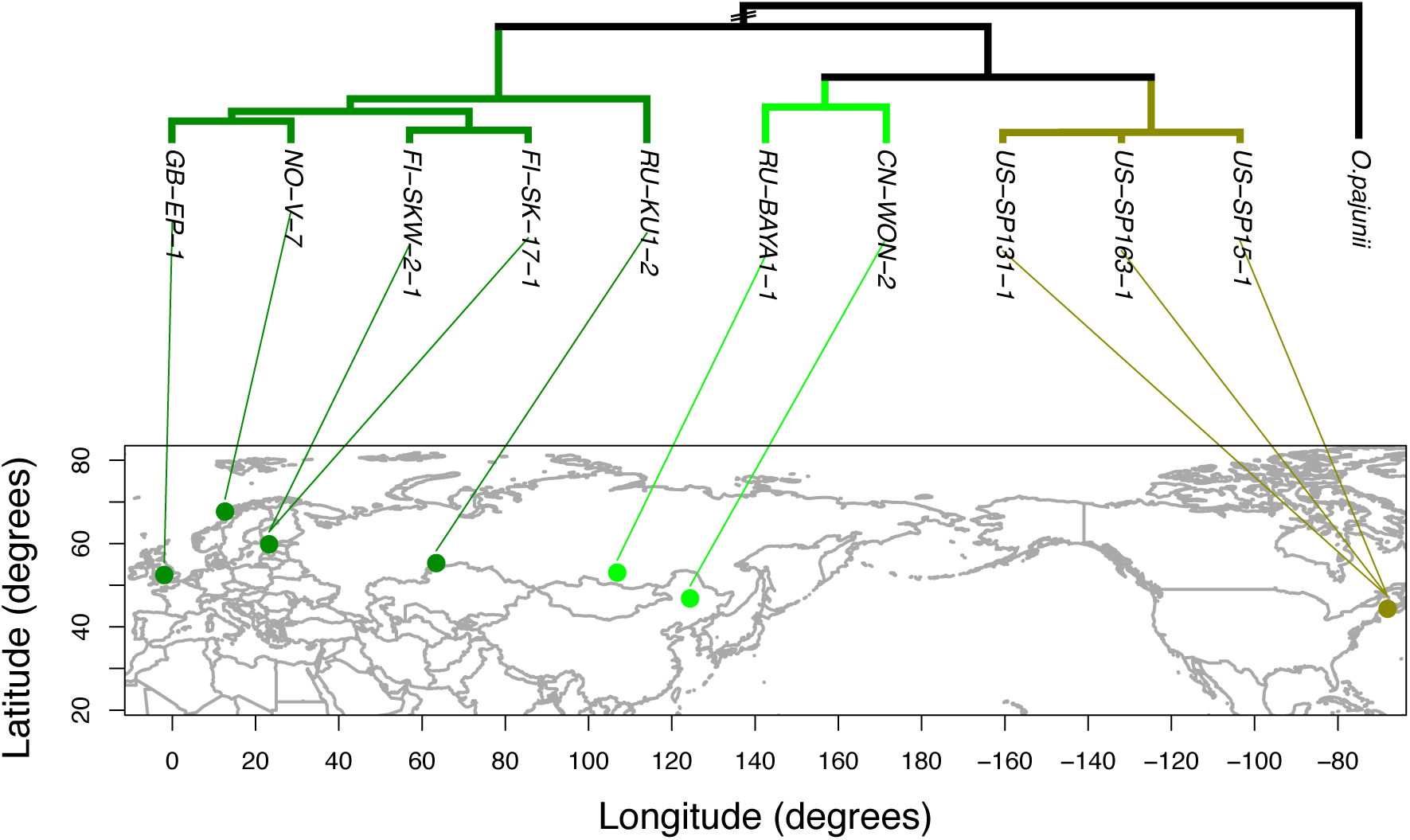
Geophylogeny of *O. colligata*. The tree is based on four-fold degenerate sites of single copy orthologs when compared to the outgroup, *O. pajunii*. Tree tips are connected to the sampling locality by colored lines, whereby colors indicate the main lineages inferred from population structure analyses: Western Eurasian (dark green), East Asian (lime), and North American (olive). See S5 Fig for the correct branch length to *O. pajunii*, as it is out of scale here for better presentation.

**Fig 5.**
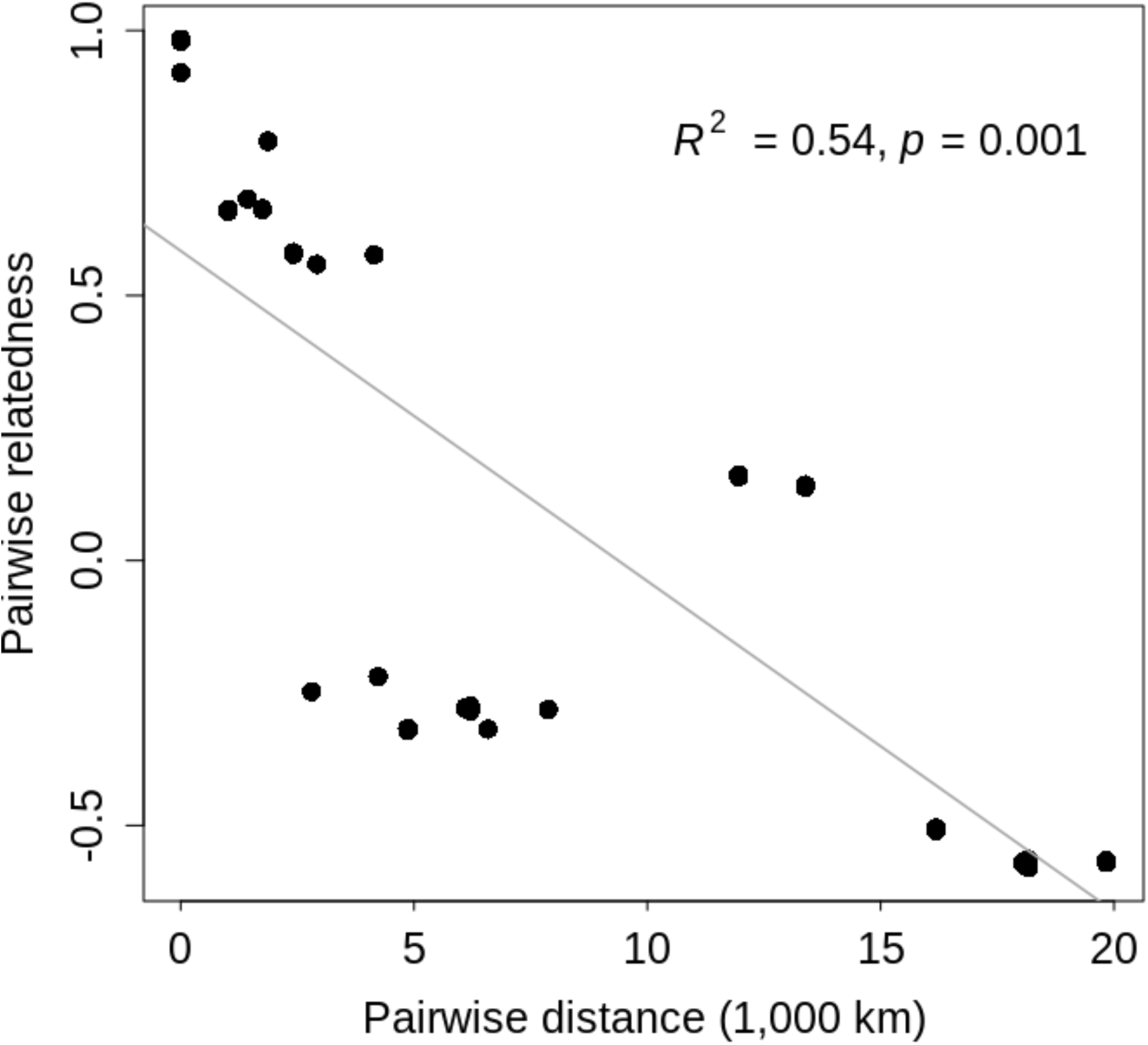
Isolation by distance (IBD). Pairwise relatedness is positively correlated with geographic distance. Each point represents the comparison between two samples. Statistics derive from dbMEM analysis by RDA. The grey line is a regression line based on a linear model.

### Mutation rate estimation

After observing the same population structure in *O. colligata* as previously found in its only host, *D. magna*, we calibrated the molecular clock in the phylogenetic analysis using the fossil-calibrated divergence time between Western Eurasian and East Asian (plus North American) host samples [36]. We found a median divergence time between *O. colligata* Western Eurasian and East Asian (plus North American) lineages of 1.61 Mya (95 % HPD interval: 0.62-4.19 Mya), which is largely overlapping with the host divergence time estimate (1.3-9.3 Mya) [36]. The median clock rate estimate was 1.04 × 10^−9^ per site per year (95 % HPD interval: 2.33 × 10^−10^-2.07 × 10^−9^ per site per year). Given published generation times of microsporidia (i.e., 63 h) [37] this would correspond to a mutation rate of 7.48 × 10^−12^ per site per generation (95 % HPD interval: 1.68 × 10^−12^-1.49 × 10^−11^ per site per generation). However, given its Northern distribution, *O. colligata*, like its host, endures the long winters as resting stage (spore) without reproducing. Assuming a resting time of three to six months, its mutation rate might be between 9.98 × 10^−12^ and 1.50 × 10^−11^ per site per generation (2.24 × 10^−12^-2.98 × 10^−11^). These values are comparable with what has previously been reported for fungi (see e.g., [38–40]).

### Efficient purifying selection

We assessed the efficacy of selection in *O. colligata* to test hypotheses about the evolution of the wide range of genome sizes observed in microsporidia. Therefore, we compared the ratio of nonsynonymous to synonymous polymorphisms, π_N_/π_S_, between BUSCO and non-BUSCO genes. The former are conserved genes among most microsporidia and are therefore expected to be under purifying selection. Indeed, BUSCO genes had a 1.5 times lower median π_N_/π_S_ ratio (0.085) than non-BUSCO genes (0.130). These values are close to an estimate for BUSCO genes in the large genome sized microsporidium *H. tvaerminnensis* (0.102). However, in *H. tvaerminnensis*, non-BUSCO genes have a three times higher median π_N_/π_S_ ratio (0.306), which is significantly different from all other (non-)BUSCO gene values mentioned here (Kruskal-Wallis *χ*^2^(3) = 309.83, *p* < 0.001). This suggests that *H. tvaerminnensis* experiences relaxed purifying selection. By implication, this means that there is relatively efficient purifying selection in *O. colligata*, likely due to its entirely vertical transmission and the consequently high *N*_*e*_.

## Discussion

The study of population genetics aims to understand temporal and spatial variation in genetic diversity, ideally within a framework of the ecology of the focal species. For parasites, this includes host–parasite interactions and the mode of transmission. Here, we take advantage of a model system in host-parasite evolutionary ecology, the planktonic freshwater species *Daphnia magna*, to explain genomic diversity, genome architecture, and population structure of one of its specific parasites, the microsporidium *O. colligata. O. colligata* is a parasite known for its small, compact genome, one of the smallest among all eukaryotic life [14]. Its host, *D. magna*, shows a clear split in its Holarctic distribution into a Western Eurasian and an East Asian (plus North American) lineage, the latter of which is subsequently divided into East Asian and North American lineages [33,35,36]. We find a pattern of co-cladogenesis with overlapping phylogeography between *O. colligata* and its host, which allows us to estimate the mutation rate of a microsporidium for the first time. Furthermore, we observe IBD, providing an explanation for the relationships among populations. We show evidence for efficient genome-wide purifying selection in *O. colligata*, which supports the hypothesis that horizontally transmitted microsporidia were able to evolve and maintain small, streamlined genomes due to high effective population sizes [8].

### Host-parasite co-phylogeography

The close interaction of the intracellular microsporidia and their hosts might shape the phylogeography of the parasites to be congruent to their host’s phylogeography — the within-species equivalent of the Fahrenholz rule [41], which states that host and parasite phylogenies show a pattern of co-cladogenesis. A strict co-phylogeography is expected if the parasite species’ mode of transmission is uniquely vertical [42], consistent with the expectation of co-dispersal as the mechanism of parasite spread [43]. Previous attempts by Pelin et al. [44] and Angst et al. [19] to reconstruct a co-phylogeography based on whole-genome data for two microsporidian species, *N. ceranae* and *H. tvaerminnensis*, respectively, remained unsuccessful, possibly due to high (human-driven) migration or non-simultaneous host-parasite expansions. In contrast, the phylogeography of *O. colligata* is congruent with the host’s phylogeography suggesting a long-term association and co-dispersal with the host. Specifically, both host and parasite phylogenies consisted of the same three geographically distinct lineages; the first lineage encompasses Western Eurasian samples, the second East Asian and the third Northern American samples. Furthermore, in contrast to the previously mentioned microsporidia (*N. ceranae* and *H. tvaerminnensis*), we found a pattern of species-wide IBD in *O. colligata*. IBD has also been described for the host, as well as for a bacterial parasite of *D. magna, Pasteuria ramosa* [32,35]. This pattern further strengthens the assumption of co-dispersal throughout the history of *O. colligata* and *D. magna*.

We have so far not detected *O. colligata* in southern regions. Diagnostic sequencing revealed that parasite samples from the Middle East, which have been identified as *O. colligata* using light microscopy [22], are other species, with very similar morphology and pathology (Fig 1). However, given the growing evidence that *O. colligata* shows an adaptation to colder habitats, it was not surprising to not find it in southern regions [30]. In contrast to *O. colligata*, we confirmed the presence of the *D. magna* specific microsporidia *Glugoides intestinalis* and *Mitosporidium daphniae* in Southern Europe using microscopy and PCR (Fig 1). Based on our results, future research should be careful relying on microscopy only for species identification, as we observed several cases of misidentification of microsporidia infecting *D. magna*.

By calibrating the phylogenetic tree of *O. colligata* with the divergence time of the Western Eurasian and East Asian (plus North American) host lineages, a time which has previously been estimated using fossil data [36], we provide the first estimated mutation rate of a microsporidium. Fungi, to which the microsporidia are phylogenetically associated, have been shown to have low mutation rates (e.g., [38–40]). Our estimate for *O. colligata* falls within the range of previously published mutation rate estimates of fungi. Therefore, we find no evidence that intracellular microsporidia evolve more rapidly than their free-living relatives, which might be expected based on the expectation that parasitism may accelerate the evolutionary process [45]. However, no other approaches to estimate mutation rates, for example, mutation accumulation experiments, have been performed for microsporidia so far.

### Genome evolution

The evolution of different genome sizes in microsporidia is an ongoing research area of great interest [14,15,19,46]. In microsporidia, genome streamlining is associated with the abandonment of entire metabolic pathways and fewer transposable elements.

Furthermore, the maintenance of streamlined microsporidian genomes was possible due to efficient purifying selection, which is only possible with large effective population sizes [8]. In contrast, small effective population sizes, for example, in species that exhibit population bottlenecks during vertical transmission, increase the power of stochastic processes and reduce selection efficacy [8,47]. Also, the proliferation of repetitive elements is less constrained in species with small effective population sizes. Therefore, we wanted to estimate the effective population size and the rate of adaptive substitution, *α* (as a measure for selection efficacy) in the horizontally transmitted *O. colligata* and compare it to another microsporidian parasite of *D. magna, H. tvaerminnensis*, which has a mixed-mode transmission and a small effective population size [19]. However, PSMC expects diploid samples to calculate *N*_*e*_ and *α* estimated using MKTs was unreliable since our data had too few intraspecies polymorphisms as opposed to the many fixed substitutions between *O. colligata* and the outgroup (the closest known relative *O. pajunii*). We would need more diversity by adding more *O. colligata* samples to the dataset or a more closely related outgroup to reliably estimate *α* using MKTs. However, consistent with the expectation of weaker purifying selection in microsporidia of larger genome sizes and additional vertical transmission, we found elevated π_N_/π_S_ ratios in non-BUSCO genes in *H. tvaerminnensis* compared to *O. colligata*. In a previous comparison between the genera *Hamiltosporidium* and *Ordospora*, mixed-mode transmission and larger genome size has been shown to be associated with larger d_N_/d_S_ ratios [8]. We extend the evidential basis with our between-species comparison of population samples.

### Describing genomic variation

We estimated genomic variation of the species-specific parasite *O. colligata* with samples covering a large part of the host’s geographic distribution. The whole-genome SNP density in *O. colligata* is slightly lower than what was found for its close relative *Encephalitozoon cuniculi* [48], which was, in contrast to the samples used in this study, isolated from different host species. The low SNP density in *O. colligata* made certain analyses unreliable, i.e., inferring historical *N*_*e*_ and MKTs. The analyses to determine ploidy levels suggest that *O. colligata* is haploid (we did not conduct cytological analyses). However, two samples were somewhat ambiguous with regard to the haploid pattern (S1 and S2 Figs), but this was in part due to the quantity of segregating polymorphisms or the potential presence of multiple infections of *O. colligata* in the same sample. Our methods to infer the ploidy level of *O. colligata* would fail, however, if *O. colligata* is di-or polyploid with extremely low levels of heterozygosity, an unlikely genomic characteristic.

While low levels of genomic diversity might constrain some population genetic analyses, other important signals might be clearer. For example, because much of the *O. colligata* genome is depauperate in SNP diversity, regions with an increased SNP density stand out from the larger genomic context (Fig 2). Some of these distinctly diverse regions have increased diversity due to host-to-parasite horizontal gene transfer, which Pombert et al. [14] previously described (chromosomes 2, 5, and 8). Specifically, the observed excess diversity may arise as the result of the unintentional mapping of *D. magna* reads to these transferred regions; thus, both the host and parasite variants are conflated. More relevant to the present study is how other outlier regions might be affected differently by evolution. Indeed, some genes with functions relevant to the evolution of microsporidia-specific traits show high sequence variation, e.g., the ribosomal protein L24 (M896_051540), a leucyl-tRNA synthetase (M896_060290), and a putative ABC-like lipid transport protein (M896_121220). The ribosomal protein L24 has been reported by Liu et al. [49] to be differentially expressed before and after the germination of spores of the microsporidium *Nosema bombycis*, suggesting its importance for the regulation of transcriptional and translation activities during the infection process. Like other protein synthetases, Melnikov et al. [50] reported that the leucyl-tRNA synthetase has degenerated in the microsporidium *Vavraia culicis*. Malfunctioning synthetases are hypothesized to produce more diverse proteomes in microsporidia than expected. ABC-like lipid transport proteins are important in parasitic protists, like microsporidia, for their role in nutrient salvage [51]. While most ABC transporters are outward exporters, some are described as importers in parasitic protists. These importers might contribute to the parasite’s nutrient uptake when living inside the host cell. Moreover, the putative ABC-like lipid transport protein M896_121220 has the fourth highest π_N_/π_S_ ratio, which is an indicator for positive selection, especially recognizable when compared directly to the genomic background. Among the proteins with high π_N_/π_S_ ratios (S2 Table), many are annotated as hypothetical proteins. The inability to annotate these genes based on sequence similarity may already be a sign of 1) their fast-evolving nature and 2) their specific function in the context of coevolution between *D. magna* and *O. colligata*.

## Conclusion

The existence of extreme genomic architectures in microsporidia has been known for a while. To understand such large-scale genomic differences, knowledge about the different species’ biology is indispensable. Previously intractable problems in intracellular parasites’ past and present evolutionary dynamics can now be studied using genomic diversity at a fine scale. Furthermore, investigating the evolution of different genome sizes in microsporidia is facilitated by comparing parasites evolving in the same host, where the host factor can be assumed to be similar. Therefore, this study profits from comparisons between two genera of microsporidia, *Hamiltosporidium* and *Ordospora*, which share the host, *D. magna*. The genomes of these microsporidian genera are affected differently by coevolution with the same host. Mainly, we think that the acquisition of vertical transmission in *Hamiltosporidium* led to the expansion of its genome. Contrarily, in the vertically transmitted *Ordospora*, large effective population size and efficient purifying selection helped maintaining its compact genome. This study adds to the growing research of microsporidia supporting the observation that the mode of transmission plays an important role in the evolution of genome size. This hypothesis would benefit from larger comparative studies based on whole genome data across the clade of the microsporidia.

## Material and Methods

### Daphnia magna Diversity Panel

We used parasites derived from material collected within the framework of a large-scale biogeographic study of the host species, *D. magna* [34,35,52,53]. From each population, animals were brought to the laboratory and one iso-female line, i.e., clone, was created. We checked these clones for infections with microsporidia by phase-contrast microscopy, using squash-preparations or samples of the gut. The panel includes whole-genome sequencing (WGS) of *D. magna* clones with Illumina paired-end reads using HiSeq 2500 and NovaSeq 6000 sequencers.

### Samples

Whole-genome sequences of ten *O. colligata* samples, each from a different *D. magna* clone collected from a different population, were obtained from the Illumina sequencing (Table 1). Sequences from clones FI-SK-17-1, NO-V-7, and GB-EP-1 were reused from Haag et al. [8] (NCBI database; SRA accession: SRP211974, Bioproject ID: PRJNA419750), sequences from clone RU-BAYA1-1 were reused from Angst et al. [19] (NCBI database; SRA accession: SRP346323, Bioproject ID: PRJNA780787), and for clone FI-SK-17-1 additional sequencing was done. We used the *D. longispina*-specific *Ordospora pajunii* for analyses that needed an outgroup (NCBI database; Bioproject ID: PRJNA630072 [54]). Some samples with observed microsporidia gut infection (i.e., based on published criteria for identifying *O. colligata* infection) showed less than 1× average whole-genome coverage for *O. colligata*. To verify the presumed parasite at species level, we performed polymerase chain reactions (PCRs) of the small subunit of microsporidian ribosomal DNA followed by Sanger sequencing (S1 Table). For this, we treated one animal per *D. magna* clone with three antibiotics (streptomycin, tetracycline, ampicillin) at a concentration of 50 mg/L each for 24 h to reduce non-focal DNA. To aid gut evacuation, we fed animals after 0 h and 12 h with dextran beads (Sephadex ‘Small’ by Sigma Aldrich: 50 mm diameter) at a concentration of 0.5 g/100 mL [55]. Afterward, we moved individuals into separate wells of an unskirted 96-well PCR plate (Eppendorf) and removed excess fluid with a sterile pipette. Subsequently, we crushed individuals using a customized rack of metallic pestles. To each well, we added 20 µL microLYSIS-Plus DNA release buffer (Microzone, West Sussex, UK) and used the manufacturer’s tough cell lysis protocol (65 °C for 15 min; 96 °C for 2 min; 65 °C for 4 min; 96 °C for 1 min; 65 °C for 1 min; 96 °C for 30 s) for incubation in an Eppendorf Mastercycler Nexus. Following this DNA extraction, we diluted the products to 0.1 × with deionized water and set up the PCR reactions. Each 50 µL PCR took place in an Eppendorf Mastercycler Nexus, with each reaction consisting of 29.5 µL deionized water, 10 µL of the diluted sample, 5 µL 10 × PCR buffer, 2 µL forward and reverse primer, 1 µL dNTPs and 0.5 µL Taq DNA polymerase. We used universal primers targeting the small subunit of microsporidian ribosomal DNA (V1f: 5’-CACCAGGTTGATTCTGCCTGAC-3’; 1342r: 5’-ACGGGCGGTGTGTACAAAGAACAG-3’) [56]. The PCR conditions were 95 °C for 15 min, followed by 40 cycles of 95 °C for 30 s, 67.5 °C for 90 s, and 72 °C for 90 s, and then 72 °C for 7 min. PCR products were sequenced by Microsynth (Balgach, Switzerland) using the Sanger method on ABI 3730/3730xl machines (Applied Biosystems; Thermo Fisher Scientific). We compared the obtained sequence data to the NIH genetic sequence database (GenBank) [57] with the Basic Local Alignment Search Tool for nucleotides (BLASTN) on the NCBI website (https://blast.ncbi.nlm.nih.gov/). The success of the DNA extraction was verified before submission with an additional PCR using a *D. magna*-specific pair of primers and the same PCR conditions, i.e., positive control on PCR conditions. Samples with a microscopically detected microsporidian infection and a positive *D. magna* control but without a PCR product for the microsporidian universal marker were assigned to infection with *M. daphniae*, another gut microsporidium [13] which is phenotypically very similar to *O. colligata*, but for which the primers would not be capable of generating an amplicon because it is a very basal microsporidium.

### Mapping and variant calling

We assessed raw sequencing reads for quality using FastQC v.0.11.7 (http://www.bioinformatics.babraham.ac.uk/projects/fastqc) and subsequently trimmed them to remove low-quality sequences and adapter contamination using Trimmomatic v.0.38 [58]. Unless otherwise stated, we used default parameters for bioinformatics programs. We assayed successful trimming using a second run of FastQC. We downloaded the genome of *O. colligata* (NCBI database; Assembly name: ASM80326v; GenBank assembly accession: GCA_000803265.1, Bioproject accession: PRJNA210314 [14]) from NCBI and used it as a reference for mapping quality-trimmed reads with BWA MEM v.0.7.17 [59], using default parameters. We converted SAM files to BAM files, coordinate-sorted individual BAM files, and removed unmapped reads using SAMtools v.1.9 [60]. Afterward, we added read groups and marked duplicates for individual BAM files using Picard Toolkit v.2.18.16 [61]. To compute the average read depth, we used SAMtools function depth, and for variant calling, we used GATK v.3.8 HaplotypeCaller in the haploid mode [62,63]. Specifically, we first generated GVCFs for individual BAM files and used the GATK function GenotypeGVCF to combine GVCFs into an all-site VCF. We filtered this VCF file to include only SNP variants (i.e., exclude INDEL variants) using VCFtools v.0.1.16 [64].

### Ploidy

We applied two different methods to infer the ploidy of *O. colligata* samples. First, we used ploidyNGS v.3.1.2 [65] to visualize frequencies of putative alleles directly from individual BAM files. Second, we output reads which mapped to the reference genome to FASTQ files using BEDtools v.2.27.1 [66] and subsequently used Kmercountexact.sh, a bash script included in BBMap v.38.22 [67] to generate k-mer frequency histograms. Based on the results of this ploidy assessment, which were consistent across the two methodologies, *O. colligata* was treated as haploid in all analyses requiring ploidy assignment.

### Coding sequences - single-copy orthologs

Phylogenetic analyses and tests for selection (especially its efficacy) mostly rely on protein-coding regions. To characterize variation in nucleotide diversity within the protein-coding sections of the *O. colligata* samples, we needed to extract subsets of the genome-wide VCF using the following approaches. Protein sequences of *O. colligata* (OC4; GenBank accession: JOKQ00000000.1) were downloaded from NCBI and complemented with those of the outgroup, *O. pajunii* (GenBank accession: JACCJH000000000.1 [54]). To find one-to-one orthologs between the species, we used protein datasets of the two as input for OrthoMCL v.2.0.9 [68]). Specifically, we followed the automated pipeline described at the following Github repository: https://github.com/apetkau/orthomcl-pipeline. We aligned the identified orthologous sequences of *O. colligata* and *O. pajunii* with PRANK v.170427 [69] using a custom script adapted from [52]. After an initial survey of pairwise alignment quality, we implemented a masking step, wherein excessively divergent or poorly aligned sequences (divergence > 0.5 %) were excluded from downstream analysis. We used the R package *seqinR* v.3.4-5 [70] for importing FASTA alignments and *PopGenome* v.2.7.1 [71] for the manipulation of the VCF file in the next step. To produce multiple sequence alignments, we generated one alternative reference for each *O. colligata* sample using the GATK-functions SelectVariants and FastaAlternateReferenceMaker. We cut out coding sequences of interest from the alternative references with the script gff2fasta.pl (https://github.com/ISUgenomics/common_scripts). To align the resultant coding sequences of the *O. colligata* ingroup samples to the *O. colligata–O. pajunii* alignments, we used MAFFT v.7.407 [72,73] and its --add option. Next, we masked all positions previously masked in the two species reference alignments in the multiple sequence-alignments using generate_masked_ranges.py (https://gist.github.com/danielecook) and BEDtools function maskfasta.

### Sequence variation and population genetic analyses

We calculated the SNP density of *O. colligata* by dividing the genome lengths by the number of SNPs across the full genome using VCFtools. For calculating per-site nucleotide differences (π) along the *O. colligata* genome, we used pixy v.0.95 [74]. Beforehand, we filtered out alleles with less than half or more than double the average sample coverage.

Finally, we averaged the estimated π-values over one Kilobase pair (Kbp) windows. Similarly, we calculated the non-/synonymous per-site nucleotide differences (π_N_ and π_S_) for the coding sequences of *O. colligata* using the script selectionStats.py (https://github.com/tatumdmortimer/popgen-stats). Both methods are consistent with theoretical expectations and comparable amongst species as they take invariant sites into account for their calculations [74]. The ratio of π_N_ and π_S_ was separately calculated for BUSCO genes (Benchmarking Universal Single-Copy Orthologs) identified using BUSCO v.4.0.1 [75] and its microsporidia_odb10 database (Creation date: 2020-08-05) and compared to published π_N_/π_S_ values of *H. tvaerminnensis* [19].

### Population structure and phylogenetic analyses

To assess population structure in *O. colligata*, we used the R v.3.5.1 [76] packages *SNPRelate* v.1.14.0 and *gdsfmt* v.1.16.0 [77] for principal component analysis (PCA) as well as cluster analysis with the whole-genome polymorphism data. Additionally, to test for IBD, we compared the samples’ pairwise relatedness with their pairwise geographic distance. Therefore, we used the R package *hierfstat* v.0.5-7 [78] to calculate the pairwise relatedness and *geodist* v.0.0.3 [79] to calculate the geographic distance between samples with the geodesic measure. We measured the distance to the North American samples across the Bering Strait to account for the likely dispersal route of host and parasite given their population structure. For file import and format conversions we used *VCFR* and *adegenet* v.2.1.2 [80] in R. We tested for association between relatedness and geographic distance using dbMEM analysis by RDA. Specifically, we transformed the explanatory variable, geographic distance, into dbMEMs using the *adespatial* v.0.3-14 [81] R package and decomposed the response variable, genetic relatedness, into principal components using the R base *stats* function prcomp. We then used the R package *vegan* v.2.5-7 [82] for RDA and assessed significance with 1,000 permutations.

In addition to looking at whole-genome diversity, we also focused directly on the phylogenetic signal in protein-coding regions of the genome. We concatenated previously extracted single-copy ortholog gene alignments into a single sequence and filtered for four-fold degenerate sites with MEGA v.7.16.0617 [83], which we used as input for Bayesian phylogenetic analysis in BEAST2 v.2.6.2 [84,85]. We prepared inputs using the graphical user-interface application BEAUti, which is part of BEAST2. We used the GTR+G substitution model, because this was the most likely model for our data according to jModelTest v.2.1.10 [86], a strict clock, the Yule model as tree prior, and a MCMC chain of 10,000,000 iterations. After observing a clear split between Western Eurasian and East Asian (plus North American) samples, we further used a most recent common ancestor prior with a log-normal distribution for this split and switched to a calibrated Yule model. Specifically, we used M = 1, S = 0.775, and offset = 0.45 for the log-normal distribution which translates to a median of 3.17 Mya and a 95 % CI of 1.21-10.2 Mya and reflects the likely range for this split in the host [36]. We investigated the convergence of the analysis using Tracer v.1.7.1 [87] and ensured that the effective sample size (ESS) of the parameters was greater than 200. For generating the final tree, the posterior sample of trees was summarized to a maximum clade credibility tree using TreeAnnotator, which is part of BEAST2, with the first 10 % of the MCMC chain discarded as burn-in. Lastly, we visualized the obtained tree on a map of the Holarctic using the R package *phytools* v.0.7-20 [88].

### Demographic history and rate of adaptive nucleotide substitutions

We applied the McDonald–Kreitman test (MKT) [89] and a more recent derivate, the asymptotic MKT [90], to estimate *α*. However, the imbalance between low intraspecific diversity and high divergence to the outgroup *O. pajunii* in the multi-sequence alignments of single-copy orthologs impeded reliable estimation of *α* (Jesús Murga-Moreno, pers. comm.). We refrained from estimating past and present *N*_*e*_ using PSMC after determining that *O. colligata* is likely haploid.

## Acknowledgements

We thank Jürgen Hottinger, Urs Stiefel, Michelle Krebs and Andrea Cabalzar for help in the field and laboratory. We thank members of the Ebert group for providing feedback on the study and the manuscript.

## Data availability statement

Analysis scripts are available at https://github.com/pascalangst/Angst_etal_2022_PLOSPathog and raw data is deposited at the NCBI SRA database (BioProject IDs PRJNA814405).

## Author contributions

All authors designed the study. DE collected the samples. PA analyzed the data and wrote the manuscript. All authors reviewed the manuscript.

## Supporting information

**S1 Fig. Putative allele frequency spectrum by ploidyNGS of Ordospora colligata samples**. Histograms of allele frequency of putative alleles in the BAM files found using ploidyNGS, where reads supported a single variant for most positions. Dark and light shadings of colors indicate reference and alternative allele, respectively.

**S2 Fig. K-mer frequency histogram of *Ordospora colligata* by kmercountexact.sh**. K-mer frequency histograms inferred using kmercountexact.sh, with a single peak for homozygous sites and no secondary peaks which would indicate di-/polyploidy.

**S3 Fig. PCA of genomic SNP data**. PC1 vs. PC2 (A) and PC1 vs. PC3 (B).

**S4 Fig. Cluster analysis of genomic SNP data**.

**S5 Fig. Bayesian tree inference of *Ordospora***. Node labels are posterior probabilities and the scale bar unit is average height.

**S1 Table. Further microsporidian infections in the collection**. Gi = *Glugoides intestinalis*, Hm = *Hamiltosporidium magnivora*, Ht = *Hamiltosporidium tvaerminnensis*, Md = *Mitosporidium daphniae*, Oc = *Ordospora colligata*. In some genomic samples, there were no sequencing reads of the parasites that were previously identified by microscopy. This led to try to identify the species with PCR using a universal marker for microsporidia. When there were spores visible under the microscope but no PCR product, the sample was tentatively assigned to be infected with *Mitosporidium daphniae*, an early diverging microsporidium, for which the microsporidian primer pair does not work.

**S2 Table. List of *O. colligata* coding sequence (CDS) alignments with π**_**N**_**/π**_**S**_ **values**. The single copy orthologous protein of *O. pajunii* is denoted in the last column. Per-CDS π and π_N_/π_S_ value were calculated using selectionstats.py. The gene annotation is derived from the original record (NCBI database; Assembly name: ASM80326v; GenBank assembly accession: GCA_000803265.1, Bioproject accession: PRJNA210314)

## References

1. Font-Porterias N, Caro-Consuegra R, Lucas-Sánchez M, Lopez M, Giménez A, Carballo-Mesa A, et al. The Counteracting Effects of Demography on Functional Genomic Variation: The Roma Paradigm. Molecular Biology and Evolution. 2021 Jul 1;38(7):2804–17.

2. Horníková M, Marková S, Lanier HC, Searle JB, Kotlík P. A dynamic history of admixture from Mediterranean and Carpathian glacial refugia drives genomic diversity in the bank vole. Ecology and Evolution. 2021;11(12):8215–25.

3. Marske KA, Thomaz AT, Knowles LL. Dispersal barriers and opportunities drive multiple levels of phylogeographic concordance in the Southern Alps of New Zealand. Molecular Ecology. 2020;29(23):4665–79.

4. Leroy T, Rousselle M, Tilak M-K, Caizergues AE, Scornavacca C, Recuerda M, et al. Island songbirds as windows into evolution in small populations. Current Biology. 2021 Mar 22;31(6):1303-1310.e4.

5. Ebert D, Fields PD. Host–parasite co-evolution and its genomic signature. Nature Reviews Genetics. 2020 Aug 28;21:754–68.

6. Chrostek E, Pelz-Stelinski K, Hurst GDD, Hughes GL. Horizontal Transmission of Intracellular Insect Symbionts via Plants. Front Microbiol. 2017 Nov 28;8:2237.

7. Russell SL, Pepper-Tunick E, Svedberg J, Byrne A, Castillo JR, Vollmers C, et al. Horizontal transmission and recombination maintain forever young bacterial symbiont genomes. PLOS Genetics. 2020 Aug 25;16(8):e1008935.

8. Haag KL, Pombert J-F, Sun Y, de Albuquerque NRM, Batliner B, Fields P, et al. Microsporidia with Vertical Transmission Were Likely Shaped by Nonadaptive Processes. Lynch M, editor. Genome Biology and Evolution. 2020 Jan 1;12(1):3599– 614.

9. Buczek K, Trytek M, Deryło K, Borsuk G, Rybicka-Jasińska K, Gryko D, et al. Bioactivity studies of porphyrinoids against microsporidia isolated from honeybees. Sci Rep. 2020 Jul 14;10(1):11553.

10. Wadi L, Reinke AW. Evolution of microsporidia: An extremely successful group of eukaryotic intracellular parasites. PLOS Pathogens. 2020 Feb 13;16(2):e1008276.

11. Murareanu BM, Sukhdeo R, Qu R, Jiang J, Reinke AW. Generation of a Microsporidia Species Attribute Database and Analysis of the Extensive Ecological and Phenotypic Diversity of Microsporidia. mBio. 2021;12(3):e01490–21.

12. Corradi N. Microsporidia: Eukaryotic Intracellular Parasites Shaped by Gene Loss and Horizontal Gene Transfers. Annu Rev Microbiol. 2015 Oct 15;69(1):167–83.

13. Haag KL, James TY, Pombert J-F, Larsson R, Schaer TMM, Refardt D, et al. Evolution of a morphological novelty occurred before genome compaction in a lineage of extreme parasites. PNAS. 2014 Oct 28;111(43):15480–5.

14. Pombert J-F, Haag KL, Beidas S, Ebert D, Keeling PJ. The Ordospora colligata Genome: Evolution of Extreme Reduction in Microsporidia and Host-To-Parasite Horizontal Gene Transfer. mBio. 2015 Jan 13;6(1):e02400–14.

15. de Albuquerque NRM, Ebert D, Haag KL. Transposable element abundance correlates with mode of transmission in microsporidian parasites. Mobile DNA. 2020 Jun 23;11:19.

16. Altermatt F, Ebert D. Genetic diversity of Daphnia magna populations enhances resistance to parasites. Ecology Letters. 2008;11(9):918–28.

17. Ebert D. Host–parasite coevolution: Insights from the Daphnia–parasite model system. Current Opinion in Microbiology. 2008 Jun 1;11(3):290–301.

18. Orlansky S, Ben-Ami F. Genetic resistance and specificity in sister taxa of Daphnia: insights from the range of host susceptibilities. Parasites Vectors. 2019 Dec;12(1):545.

19. Angst P, Ebert D, Fields PD. Demographic history shapes genomic variation in an intracellular parasite with a wide geographic distribution. Molecular Ecology. 0(0):1– 17.

20. Decaestecker E, Declerck S, De Meester L, Ebert D. Ecological implications of parasites in natural Daphnia populations. Oecologia. 2005 Jul;144(3):382–90.

21. Ebert D, Hottinger JW, Pajunen VI. Temporal and Spatial Dynamics of Parasite Richness in a Daphnia Metapopulation. Ecology. 2001;82(12):3417–34.

22. Goren L, Ben-Ami F. Ecological correlates between cladocerans and their endoparasites from permanent and rain pools: patterns in community composition and diversity. Hydrobiologia. 2013 Jan 1;701(1):13–23.

23. Larsson JIR, Ebert D, Vávra J, Voronin VN. Redescriptjon of Pleistophora intestinalis Chatton, 1907, a microsporidian parasite of Daphnia magna and Daphnia puiex, with establishment of the new genus Glugoides (Microspora, glugeidae). European Journal of Protistology. 1996 May;32(2):251–61.

24. Larsson JIR, Ebert D, Vávra J. Ultrastructural study and description of Ordospora colligata gen. et sp. nov. (microspora, ordosporidae fam. nov.), a new microsporidian parasite of Daphnia magna (Crustacea, Cladocera). European Journal of Protistology. 1997 Dec 17;33(4):432–43.

25. Williams BAP, Hamilton KM, Jones MD, Bass D. Group-specific environmental sequencing reveals high levels of ecological heterogeneity across the microsporidian radiation. Environmental Microbiology Reports. 2018;10(3):328–36.

26. Ebert D, Lipsitch M, Mangin KL. The Effect of Parasites on Host Population Density and Extinction: Experimental Epidemiology with Daphnia and Six Microparasites. The American Naturalist. 2000 Nov 1;156(5):459–77.

27. Kirk D, Luijckx P, Stanic A, Krkošek M. Predicting the Thermal and Allometric Dependencies of Disease Transmission via the Metabolic Theory of Ecology. The American Naturalist. 2019 May 1;193(5):661–76.

28. Manzi F, Halle S, Seemann L, Ben-Ami F, Wolinska J. Sequential infection of Daphnia magna by a gut microsporidium followed by a haemolymph yeast decreases transmission of both parasites. Parasitology. 2021 Nov;148(13):1566–77.

29. Refardt D, Ebert D. Inference of parasite local adaptation using two different fitness components. J Evol Biol. 2007 May;20(3):921–9.

30. Kirk D, Jones N, Peacock S, Phillips J, Molnár PK, Krkošek M, et al. Empirical evidence that metabolic theory describes the temperature dependency of within-host parasite dynamics. PLOS Biology. 2018 Feb 7;16(2):e2004608.

31. Ebert D. Ecology, epidemiology, and evolution of parasitism in Daphnia. Bethesda (MD): National Library of Medicine (US), National Center for Biotechnology Information; 2005.

32. Andras JP, Fields PD, Ebert D. Spatial population genetic structure of a bacterial parasite in close coevolution with its host. Molecular Ecology. 2018;27(6):1371–84.

33. Bekker EI, Karabanov DP, Galimov YR, Haag CR, Neretina TV, Kotov AA. Phylogeography of Daphnia magna Straus (Crustacea: Cladocera) in Northern Eurasia: Evidence for a deep longitudinal split between mitochondrial lineages. PLOS ONE. 2018 Mar 15;13(3):e0194045.

34. Fields PD, Reisser C, Dukić M, Haag CR, Ebert D. Genes mirror geography in Daphnia magna. Molecular Ecology. 2015;24(17):4521–36.

35. Fields PD, Obbard DJ, McTaggart SJ, Galimov Y, Little TJ, Ebert D. Mitogenome phylogeographic analysis of a planktonic crustacean. Molecular Phylogenetics and Evolution. 2018 Dec 1;129:138–48.

36. Cornetti L, Fields PD, Van Damme K, Ebert D. A fossil-calibrated phylogenomic analysis of Daphnia and the Daphniidae. Molecular Phylogenetics and Evolution. 2019 Aug 1;137:250–62.

37. Kramer JP. Generation time of the microsporidian Octosporea muscaedomesticae flu in adult Phormia regina (Meigen) (Diptera, Calliphoridae). Z Parasitenkd. 1965 Mar 3;25:309–13.

38. Bezmenova AV, Zvyagina EA, Fedotova AV, Kasianov AS, Neretina TV, Penin AA, et al. Rapid Accumulation of Mutations in Growing Mycelia of a Hypervariable Fungus Schizophyllum commune. Molecular Biology and Evolution. 2020 Aug 1;37(8):2279–86.

39. Ene IV, Farrer RA, Hirakawa MP, Agwamba K, Cuomo CA, Bennett RJ. Global analysis of mutations driving microevolution of a heterozygous diploid fungal pathogen. Proc Natl Acad Sci U S A. 2018 Sep 11;115(37):E8688–97.

40. Zhu YO, Siegal ML, Hall DW, Petrov DA. Precise estimates of mutation rate and spectrum in yeast. Proc Natl Acad Sci U S A. 2014 Jun 3;111(22):E2310–2318.

41. Fahrenholz H. Ectoparasiten und Abstammungslehre. Zoologischer Anzeiger. 1913;(41):371–4.

42. Werren JH, Baldo L, Clark ME. Wolbachia: master manipulators of invertebrate biology. Nat Rev Microbiol. 2008 Oct;6(10):741–51.

43. Page RDM. Tangled Trees: Phylogeny, Cospeciation, and Coevolution. University of Chicago Press; 2003. 364 p.

44. Pelin A, Selman M, Aris-Brosou S, Farinelli L, Corradi N. Genome analyses suggest the presence of polyploidy and recent human-driven expansions in eight global populations of the honeybee pathogen Nosema ceranae. Environ Microbiol. 2015 Nov;17(11):4443–58.

45. Moran NA. Accelerated evolution and Muller’s rachet in endosymbiotic bacteria. Proc Natl Acad Sci U S A. 1996 Apr 2;93(7):2873–8.

46. Parisot N, Pelin A, Gasc C, Polonais V, Belkorchia A, Panek J, et al. Microsporidian genomes harbor a diverse array of transposable elements that demonstrate an ancestry of horizontal exchange with metazoans. Genome Biol Evol. 2014 Aug 28;6(9):2289–300.

47. Lynch M. The origins of genome architecture. Sunderland (MA): Sinauer Associates.; 2007.

48. Pombert J-F, Xu J, Smith DR, Heiman D, Young S, Cuomo CA, et al. Complete Genome Sequences from Three Genetically Distinct Strains Reveal High Intraspecies Genetic Diversity in the Microsporidian Encephalitozoon cuniculi. Eukaryotic Cell. 2013 Apr 1;12(4):503–11.

49. Liu H, Chen B, Hu S, Liang X, Lu X, Shao Y. Quantitative Proteomic Analysis of Germination of Nosema bombycis Spores under Extremely Alkaline Conditions. Front Microbiol. 2016 Sep 21;7:1459.

50. Melnikov SV, Rivera KD, Ostapenko D, Makarenko A, Sanscrainte ND, Becnel JJ, et al. Error-prone protein synthesis in parasites with the smallest eukaryotic genome. Proc Natl Acad Sci U S A. 2018 Jul 3;115(27):E6245–53.

51. Dean P, Major P, Nakjang S, Hirt RP, Embley TM. Transport proteins of parasitic protists and their role in nutrient salvage. Front Plant Sci. 2014 Apr 29;5.

52. Fields PD, McTaggart SJ, Reisser CMO, Haag C, Palmer WH, Little TJ, et al. Population-genomic analysis identifies a low rate of global adaptive fixation in the proteins of the cyclical parthenogen Daphnia magna. Journal of Molecular Biology and Evolution. 2022;msac048.

53. Seefeldt L, Ebert D. Temperature-versus precipitation-limitation shape local temperature tolerance in a Holarctic freshwater crustacean. Proceedings of the Royal Society B: Biological Sciences. 2019 Jul 24;286(1907):20190929.

54. de Albuquerque N, Haag K, Fields P, Cabalzar A, Ben-Ami F, Pombert J-F, et al. A new microsporidian parasite, Ordospora pajunii sp. nov (Ordosporidae), of Daphnia longispina highlights the value of genomic data for delineating species boundaries. Journal of Eukaryotic Microbiology.

55. Dukić M, Berner D, Roesti M, Haag CR, Ebert D. A high-density genetic map reveals variation in recombination rate across the genome of Daphnia magna. BMC Genetics. 2016 Oct 13;17(1):137.

56. Madyarova EV, Adelshin RV, Dimova MD, Axenov-Gribanov DV, Lubyaga YA, Timofeyev MA. Microsporidian Parasites Found in the Hemolymph of Four Baikalian Endemic Amphipods. PLOS ONE. 2015 Jun 18;10(6):e0130311.

57. Benson DA, Cavanaugh M, Clark K, Karsch-Mizrachi I, Lipman DJ, Ostell J, et al. GenBank. Nucleic Acids Res. 2013 Jan;41(Database issue):D36–42.

58. Bolger AM, Lohse M, Usadel B. Trimmomatic: a flexible trimmer for Illumina sequence data. Bioinformatics. 2014 Aug 1;30(15):2114–20.

59. Li H. Aligning sequence reads, clone sequences and assembly contigs with BWA-MEM. arXiv:13033997 [q-bio]. 2013 May 26;

60. Li H, Handsaker B, Wysoker A, Fennell T, Ruan J, Homer N, et al. The Sequence Alignment/Map format and SAMtools. Bioinformatics. 2009 Aug 15;25(16):2078–9.

61. Broad Institute. “Picard Toolkit.” [Internet]. Broad Institute, GitHub Repository; 2019. Available from: http://broadinstitute.github.io/picard/

62. McKenna A, Hanna M, Banks E, Sivachenko A, Cibulskis K, Kernytsky A, et al. The Genome Analysis Toolkit: A MapReduce framework for analyzing next-generation DNA sequencing data. Genome Res. 2010 Jan 9;20(9):1297–303.

63. Van der Auwera GA, Carneiro MO, Hartl C, Poplin R, del Angel G, Levy-Moonshine A, et al. From FastQ data to high confidence variant calls: the Genome Analysis Toolkit best practices pipeline. Curr Protoc Bioinformatics. 2013 Oct 15;11(1110):11.10.1-11.10.33.

64. Danecek P, Auton A, Abecasis G, Albers CA, Banks E, DePristo MA, et al. The variant call format and VCFtools. Bioinformatics. 2011 Aug 1;27(15):2156–8.

65. Augusto Corrêa dos Santos R, Goldman GH, Riaño-Pachón DM. ploidyNGS: visually exploring ploidy with Next Generation Sequencing data. Bioinformatics. 2017 Aug 15;33(16):2575–6.

66. Quinlan AR. BEDTools: the Swiss-army tool for genome feature analysis. Current protocols in bioinformatics. 2014 Sep 8;47:11.12.1-11.12.34.

67. Bushnell B. BBMap [Internet]. 2016. Available from: https://sourceforge.net/projects/bbmap/

68. Li L, Stoeckert CJ, Roos DS. OrthoMCL: Identification of Ortholog Groups for Eukaryotic Genomes. Genome Res. 2003 Jan 9;13(9):2178–89.

69. Löytynoja A. Phylogeny-aware alignment with PRANK. In: Russell DJ, editor. Multiple Sequence Alignment Methods. Totowa, NJ: Humana Press; 2014. p. 155–70. (Methods in Molecular Biology).

70. Charif D, Lobry JR. SeqinR 1.0-2: A Contributed Package to the R Project for Statistical Computing Devoted to Biological Sequences Retrieval and Analysis. In: Bastolla U, Porto M, Roman HE, Vendruscolo M, editors. Structural Approaches to Sequence Evolution: Molecules, Networks, Populations. Berlin, Heidelberg: Springer; 2007. p. 207–32. (Biological and Medical Physics, Biomedical Engineering).

71. Pfeifer B, Wittelsbürger U, Ramos-Onsins SE, Lercher MJ. PopGenome: An Efficient Swiss Army Knife for Population Genomic Analyses in R. Mol Biol Evol. 2014 Jul 1;31(7):1929–36.

72. Katoh K, Misawa K, Kuma K, Miyata T. MAFFT: a novel method for rapid multiple sequence alignment based on fast Fourier transform. Nucleic Acids Res. 2002 Jul 15;30(14):3059–66.

73. Katoh K, Standley DM. MAFFT Multiple Sequence Alignment Software Version 7: Improvements in Performance and Usability. Mol Biol Evol. 2013 Apr 1;30(4):772–80.

74. Korunes KL, Samuk K. pixy: Unbiased estimation of nucleotide diversity and divergence in the presence of missing data. bioRxiv. 2020 Jun 28;2020.06.27.175091.

75. Seppey M, Manni M, Zdobnov EM. BUSCO: Assessing Genome Assembly and Annotation Completeness. In: Kollmar M, editor. Gene Prediction: Methods and Protocols. New York, NY: Springer; 2019. p. 227–45. (Methods in Molecular Biology).

76. R Core Team. R: The R Project for Statistical Computing [Internet]. Vienna, Austria: R Foundation for Statistical Computing; 2018. Available from: https://www.r-project.org/

77. Zheng X, Levine D, Shen J, Gogarten SM, Laurie C, Weir BS. A high-performance computing toolset for relatedness and principal component analysis of SNP data. Bioinformatics. 2012 Dec 1;28(24):3326–8.

78. Goudet J, Thibaut J. hierfstat: Estimation and Tests of Hierarchical F-Statistics. 2020.

79. Padgham M, Sumner MD. geodist: Fast, Dependency-Free Geodesic Distance Calculations. 2019.

80. Jombart T. adegenet: a R package for the multivariate analysis of genetic markers. Bioinformatics. 2008 Jun 1;24(11):1403–5.

81. Dray S, Bauman D, Blanchet G, Borcard D, Clappe S, Guenard G, et al. adespatial: Multivariate Multiscale Spatial Analysis [Internet]. 2021. Available from: https://CRAN.R-project.org/package=adespatial

82. Oksanen J, Blanchet FG, Friendly M, Kindt R, Legendre P, McGlinn D, et al. vegan: Community Ecology Package [Internet]. 2020. Available from: https://CRAN.R-project.org/package=vegan

83. Kumar S, Stecher G, Tamura K. MEGA7: Molecular Evolutionary Genetics Analysis Version 7.0 for Bigger Datasets. Mol Biol Evol. 2016 Jul 1;33(7):1870–4.

84. Bouckaert R, Vaughan TG, Barido-Sottani J, Duchêne S, Fourment M, Gavryushkina A, et al. BEAST 2.5: An advanced software platform for Bayesian evolutionary analysis. Pertea M, editor. PLoS Comput Biol. 2019 Apr 8;15(4):e1006650.

85. Drummond AJ, Rambaut A, Shapiro B, Pybus OG. Bayesian Coalescent Inference of Past Population Dynamics from Molecular Sequences. Mol Biol Evol. 2005 May 1;22(5):1185–92.

86. Darriba D, Taboada GL, Doallo R, Posada D. jModelTest 2: more models, new heuristics and parallel computing. Nat Methods. 2012 Aug;9(8):772–772.

87. Rambaut A, Drummond AJ, Xie D, Baele G, Suchard MA. Posterior Summarization in Bayesian Phylogenetics Using Tracer 1.7. Syst Biol. 2018 Sep 1;67(5):901–4.

88. Revell LJ. phytools: an R package for phylogenetic comparative biology (and other things). Methods in Ecology and Evolution. 2012;3(2):217–23.

89. McDonald JH, Kreitman M. Adaptive protein evolution at the Adh locus in Drosophila. Nature. 1991 Jun 20;351(6328):652–4.

90. Haller BC, Messer PW. asymptoticMK: A Web-Based Tool for the Asymptotic McDonald–Kreitman Test. G3: Genes, Genomes, Genetics. 2017 May 1;7(5):1569–75.

