## Supplementary figures and images for "Population genetic analysis reveals the role of natural selection and phylogeography on genome-wide diversity in an extremely compact and reduced microsporidian genome"

### Supplemental Figure 1

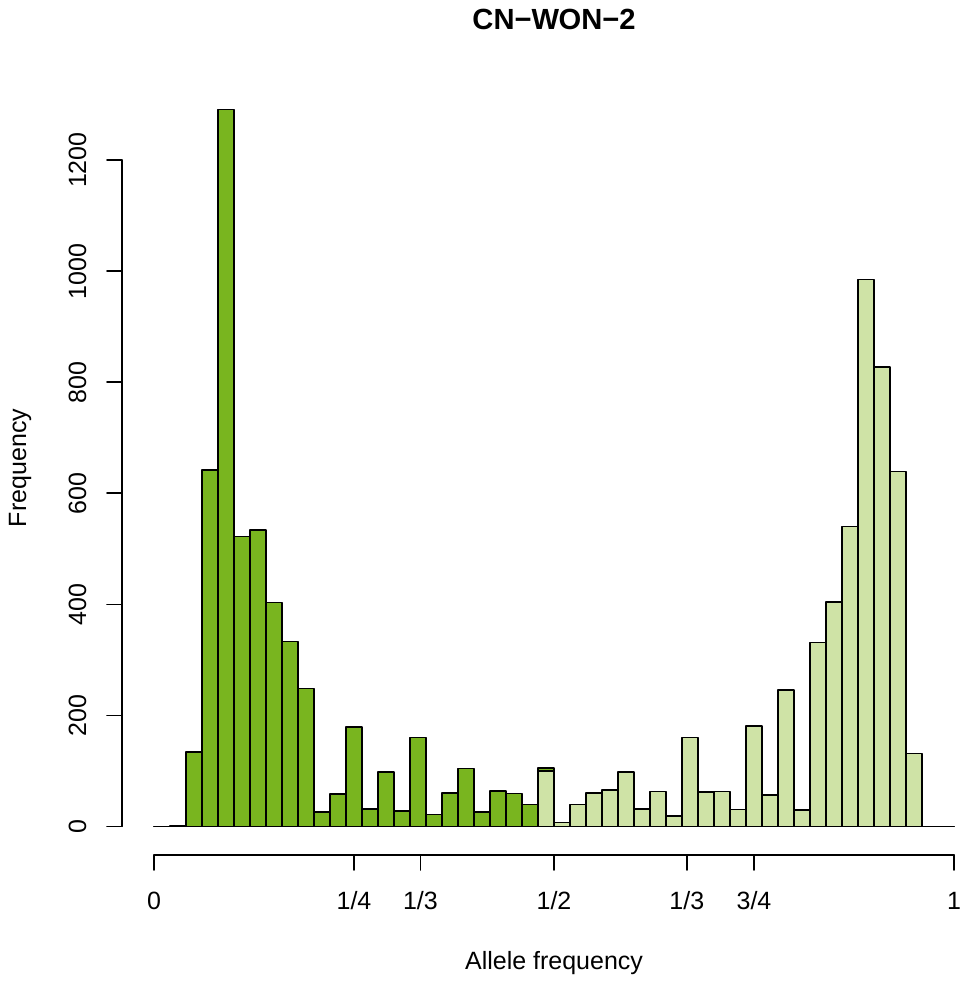

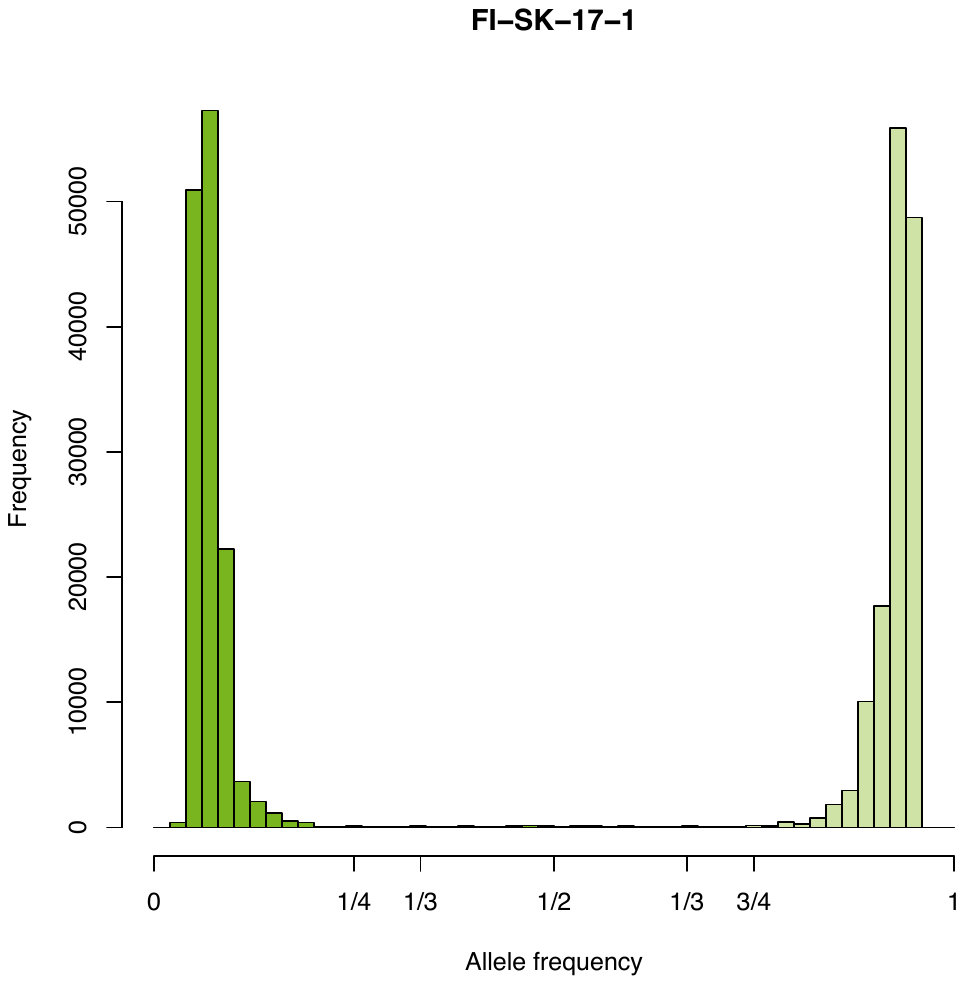

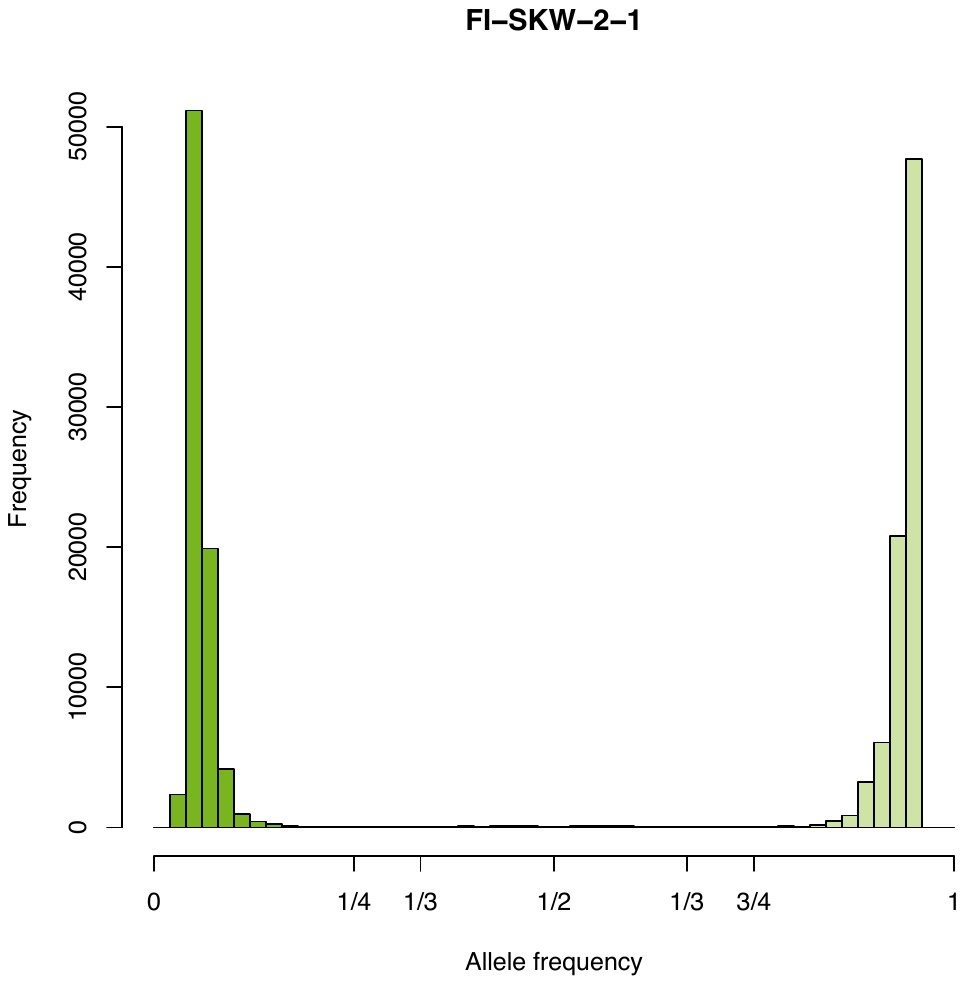

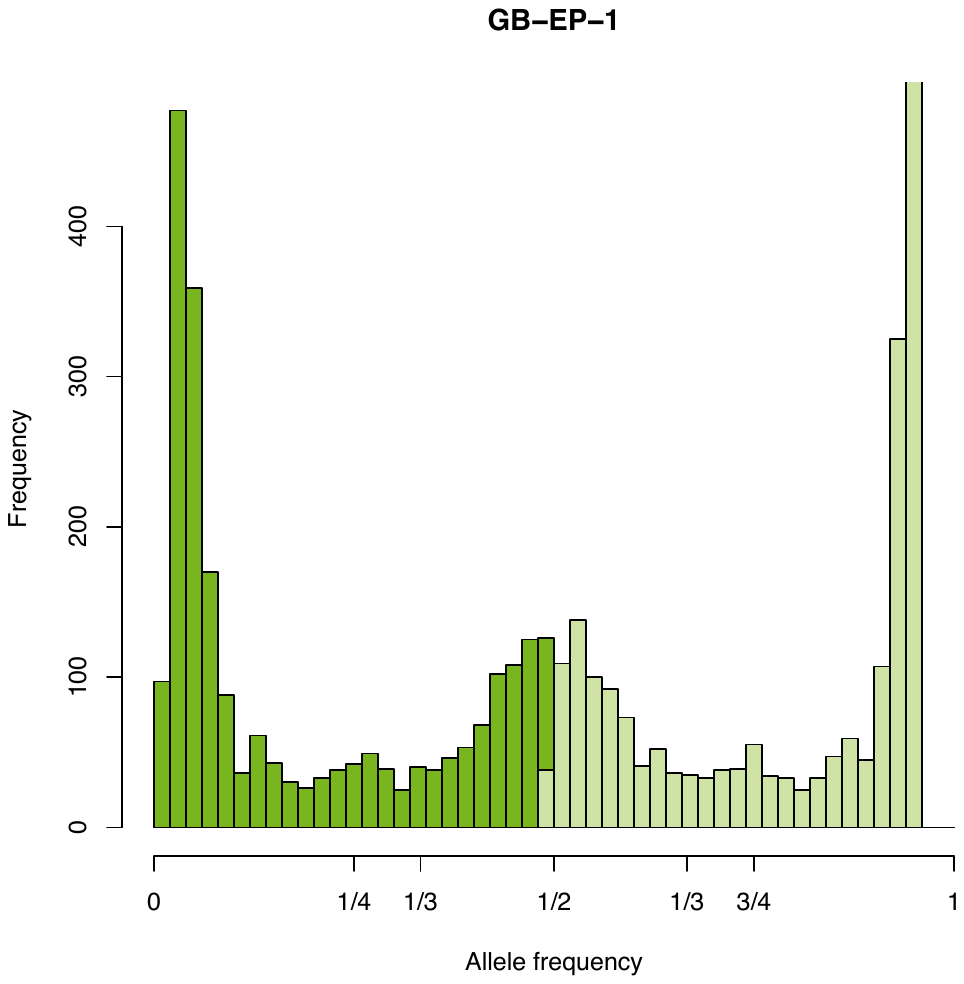

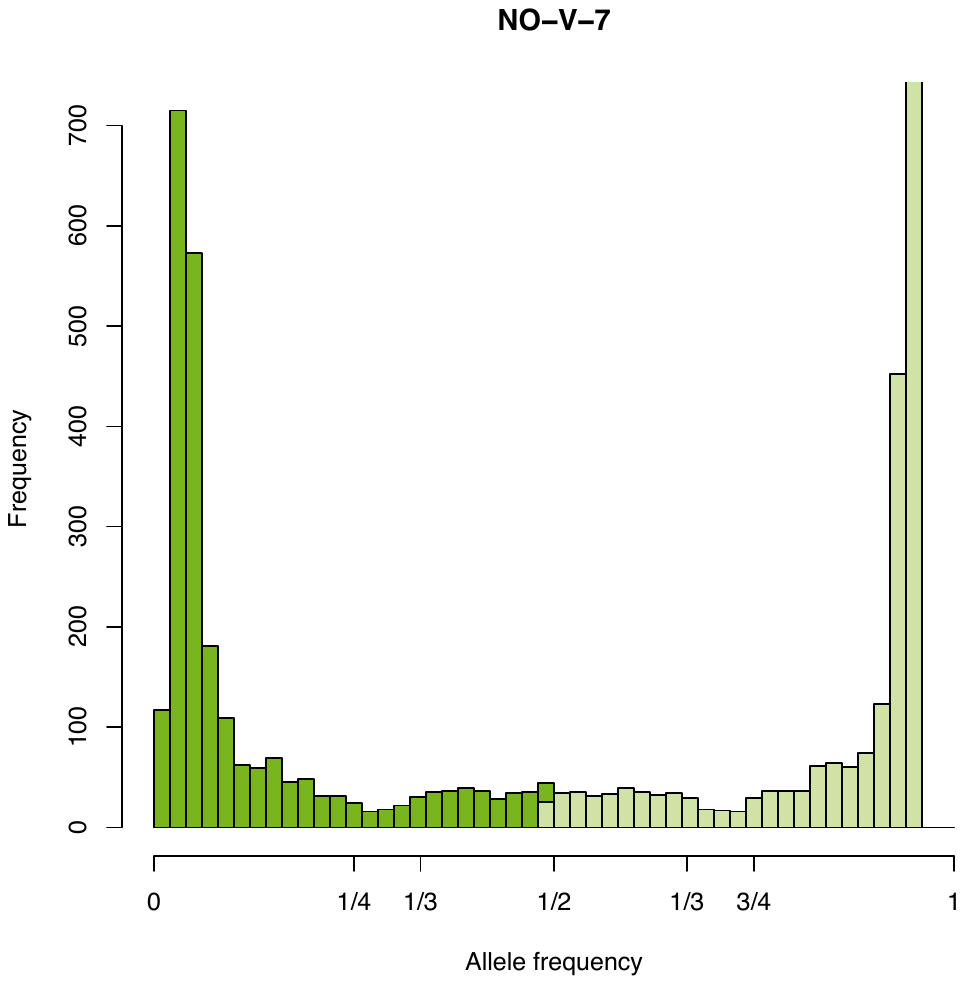

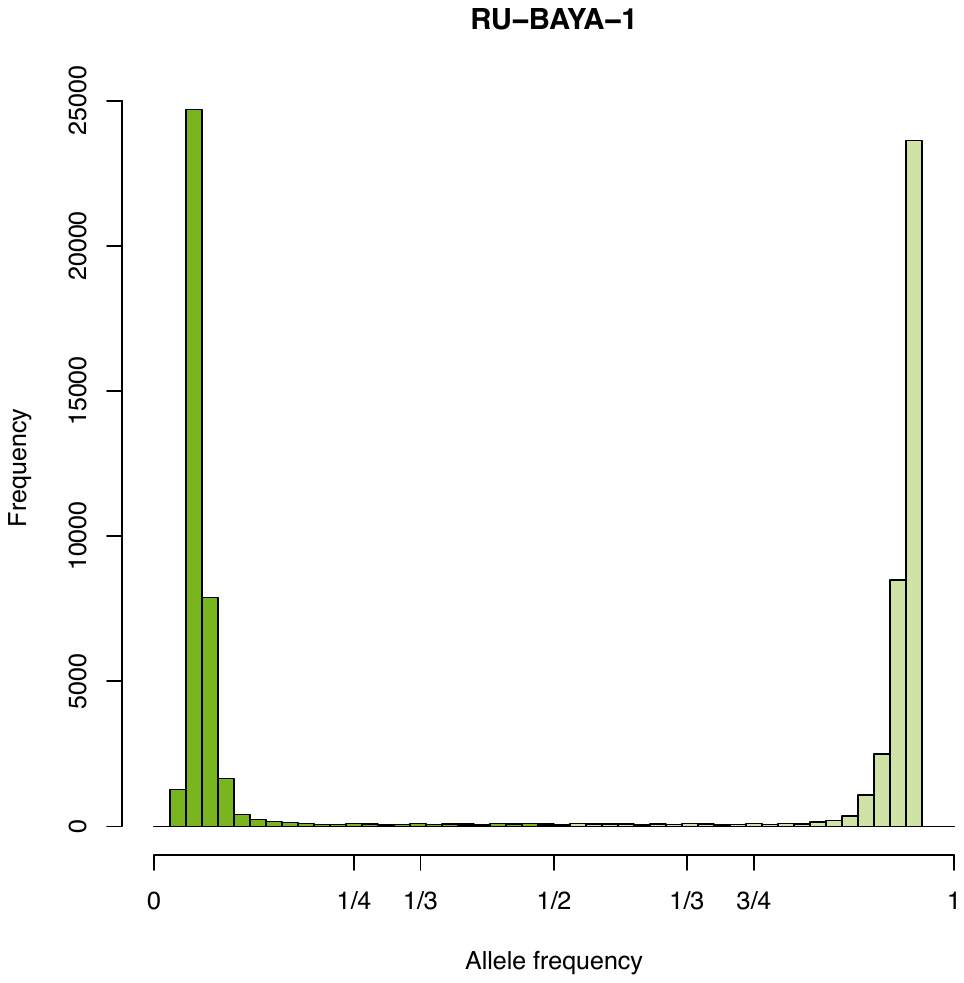

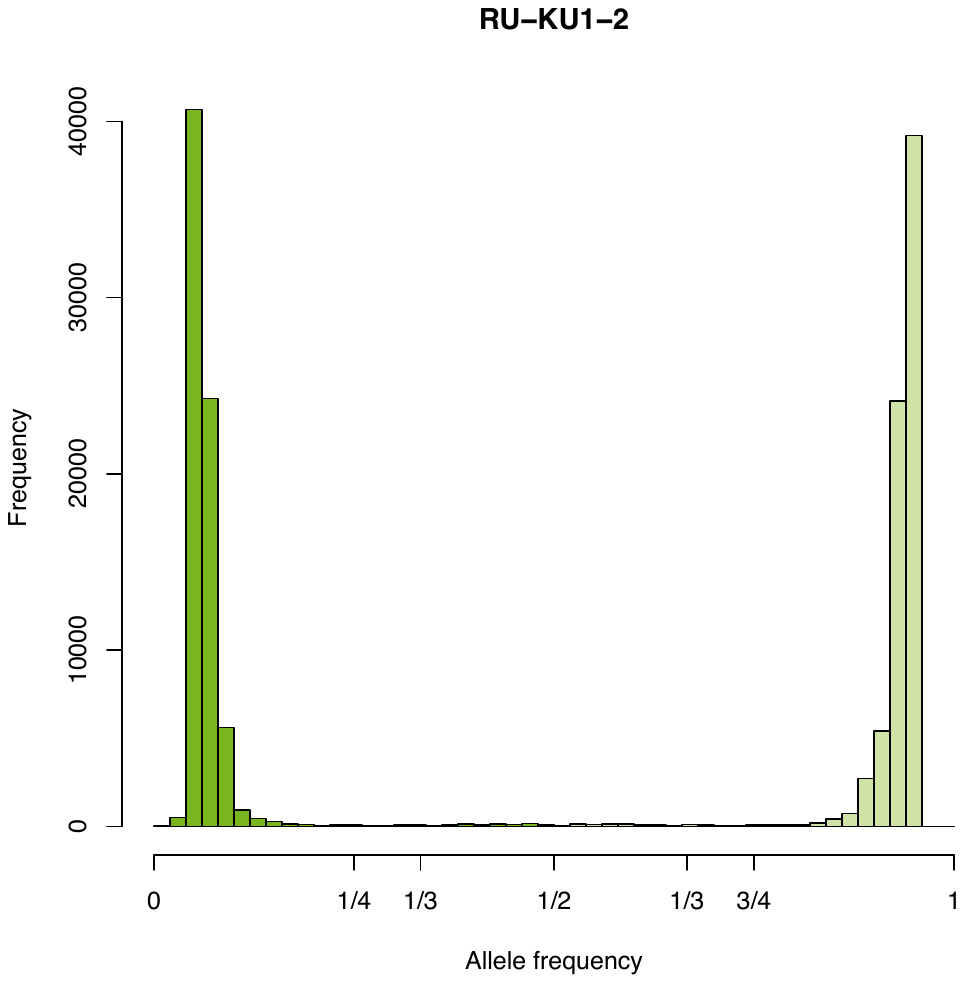

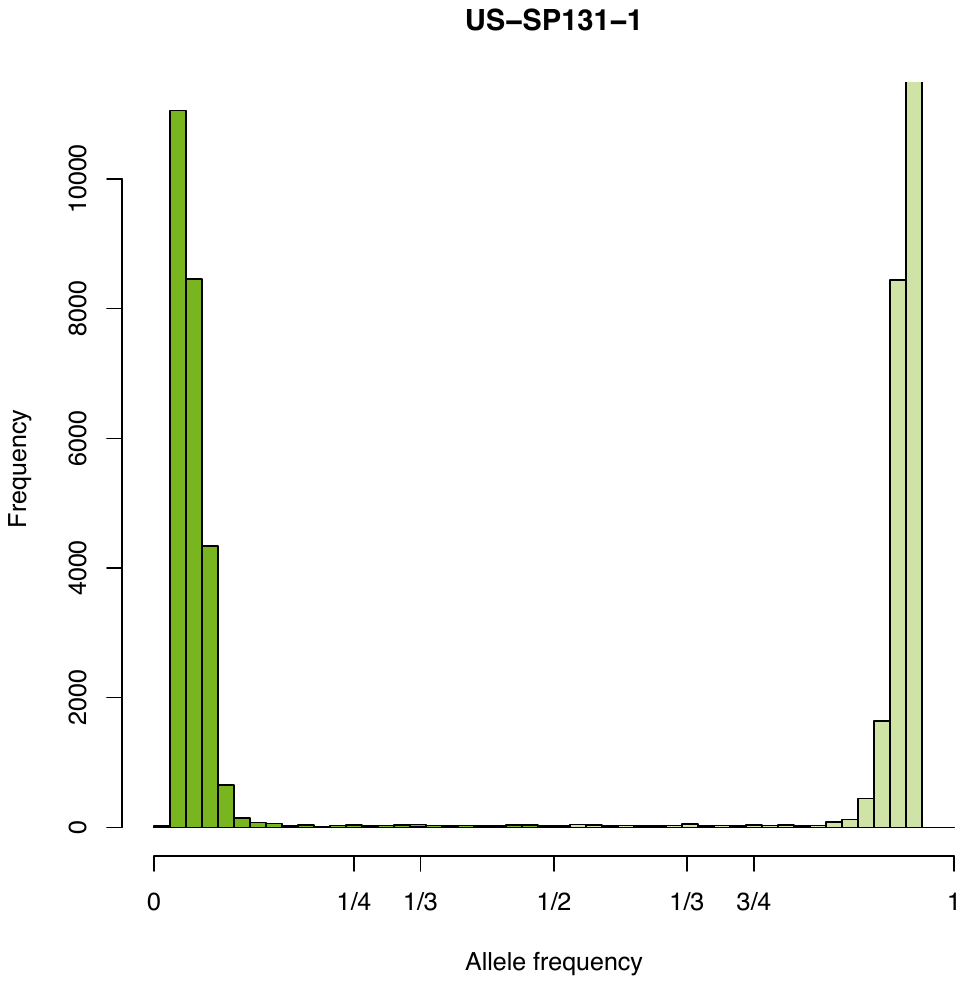

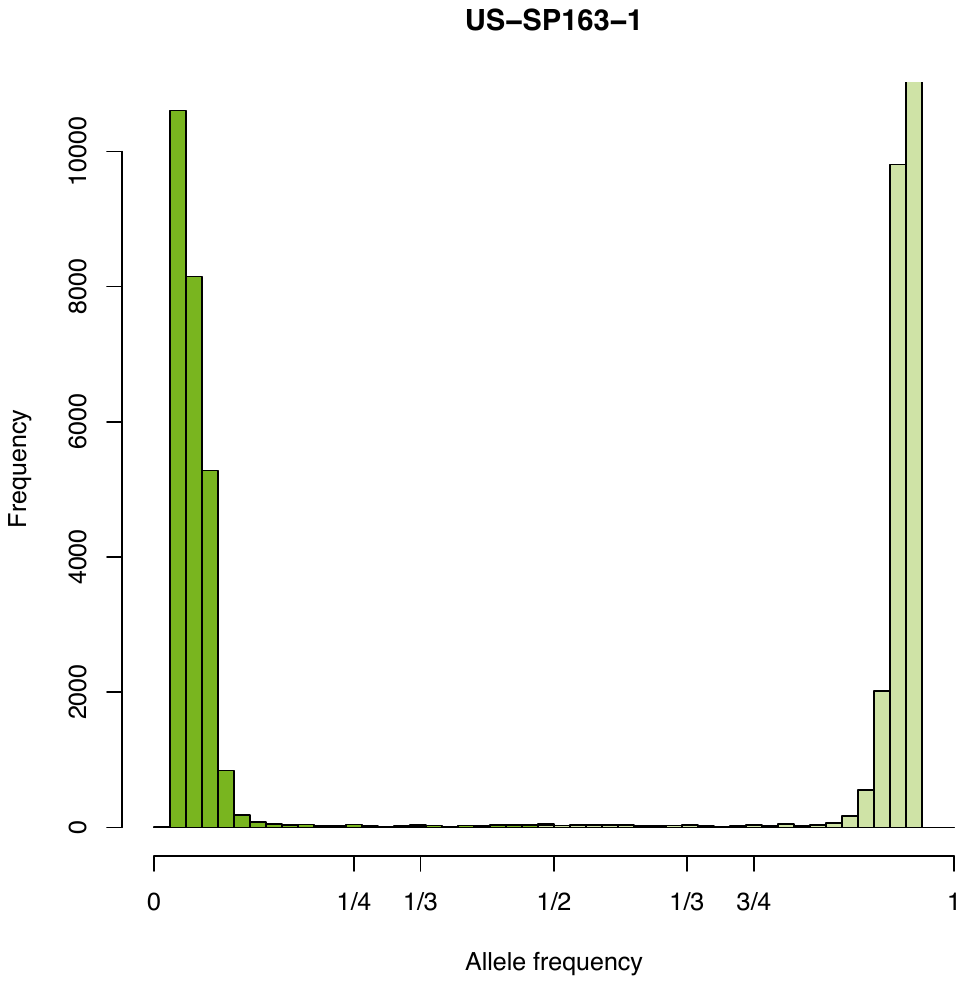

### Supplemental Figure 2

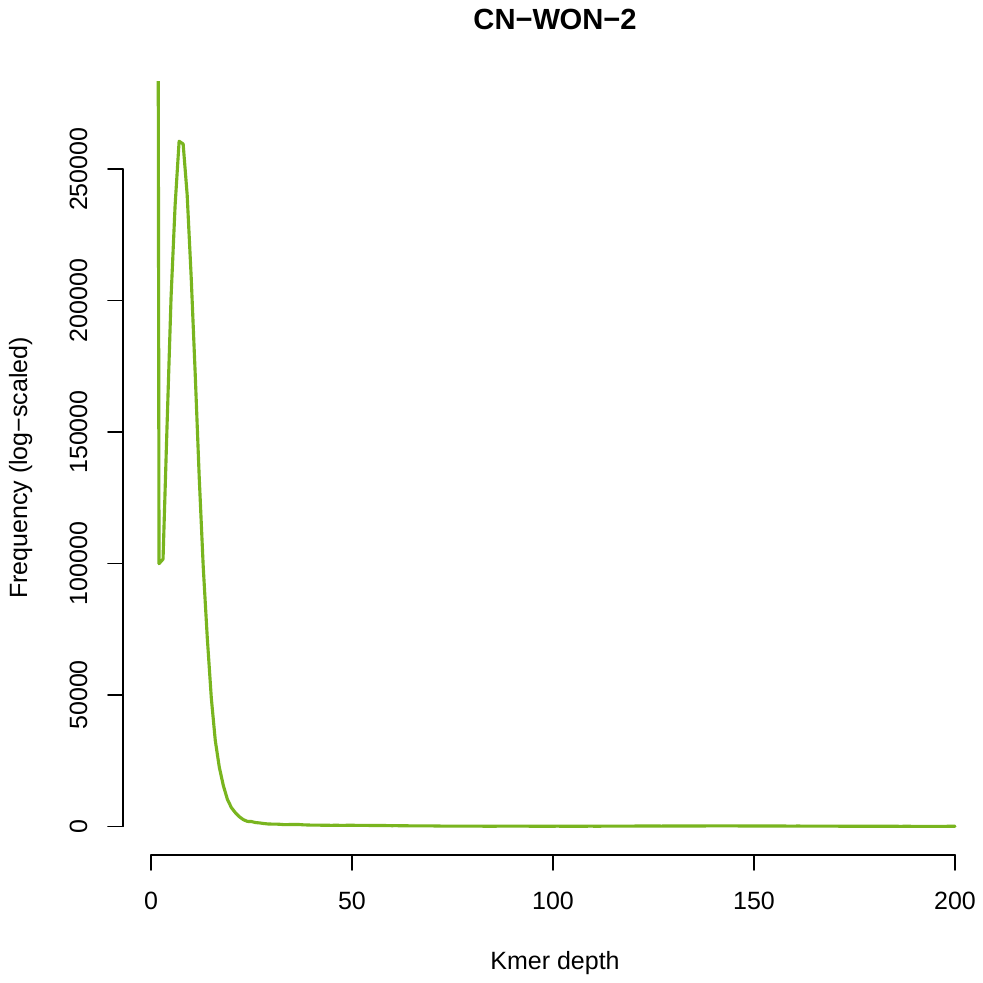

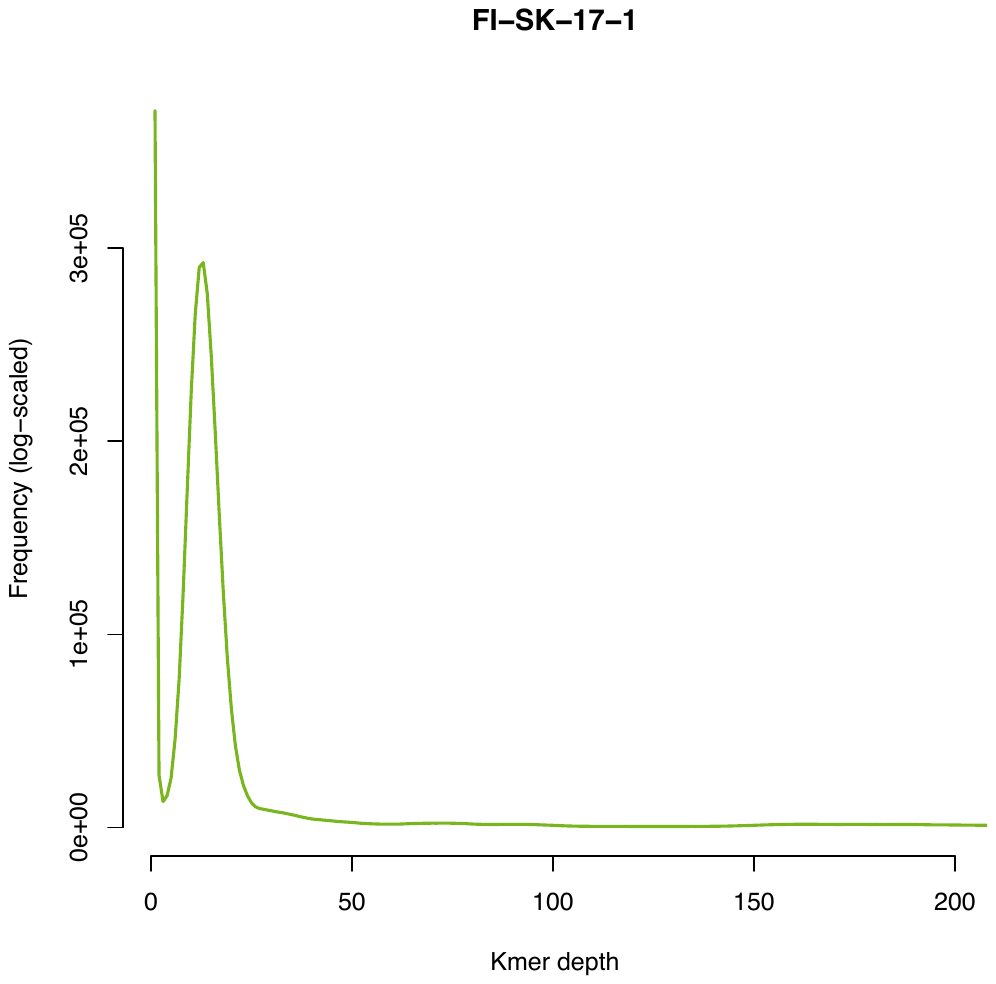

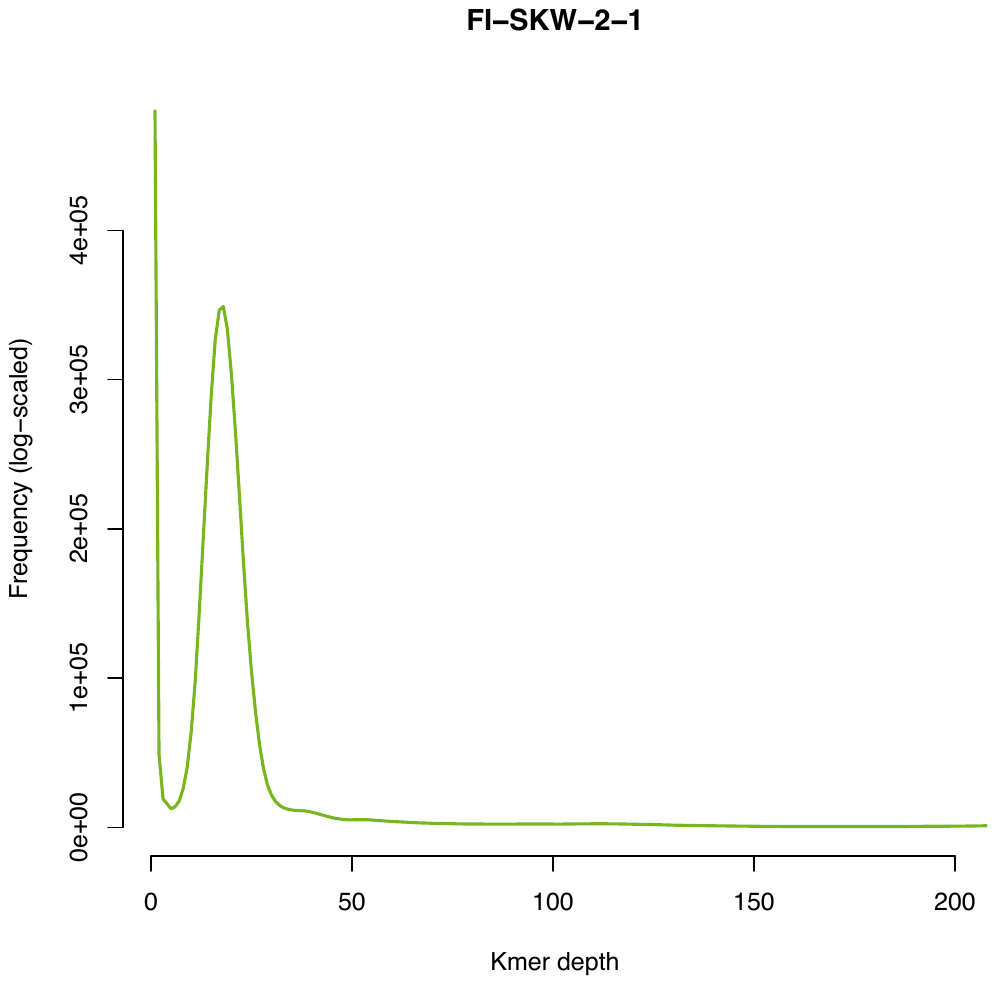

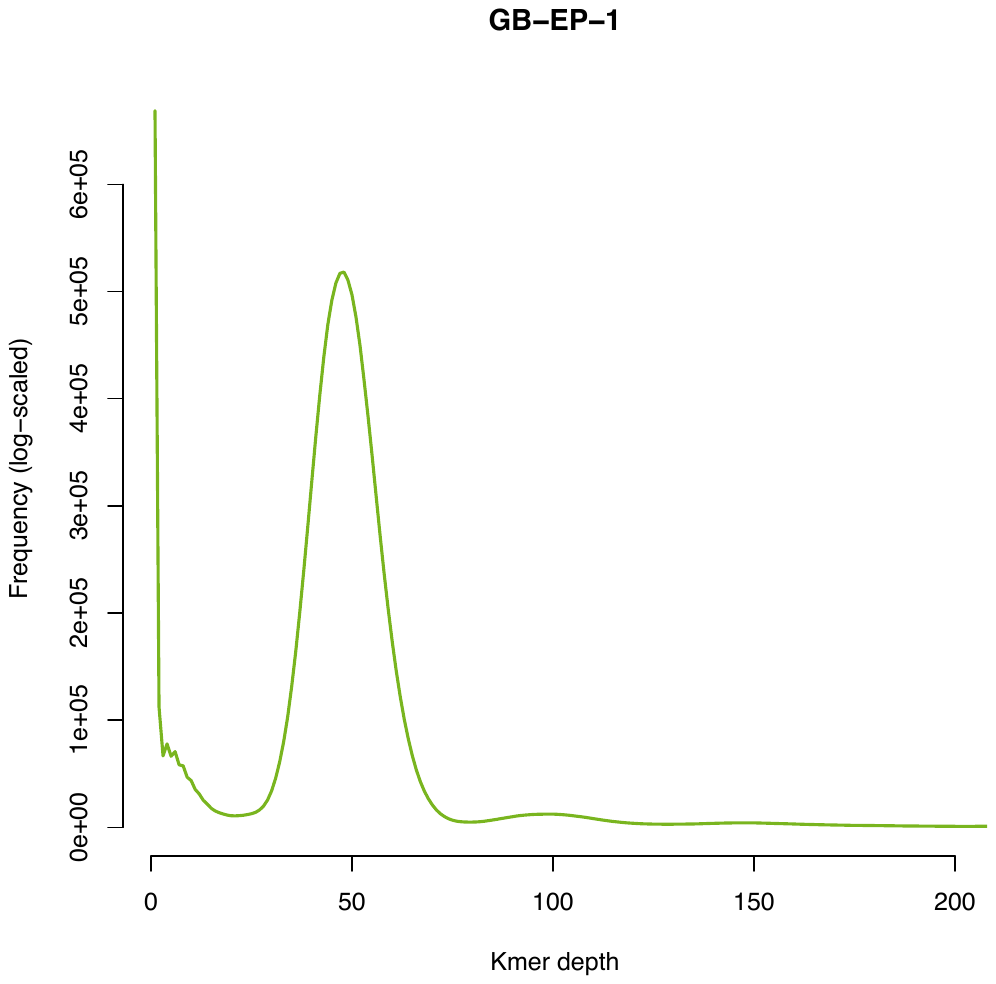

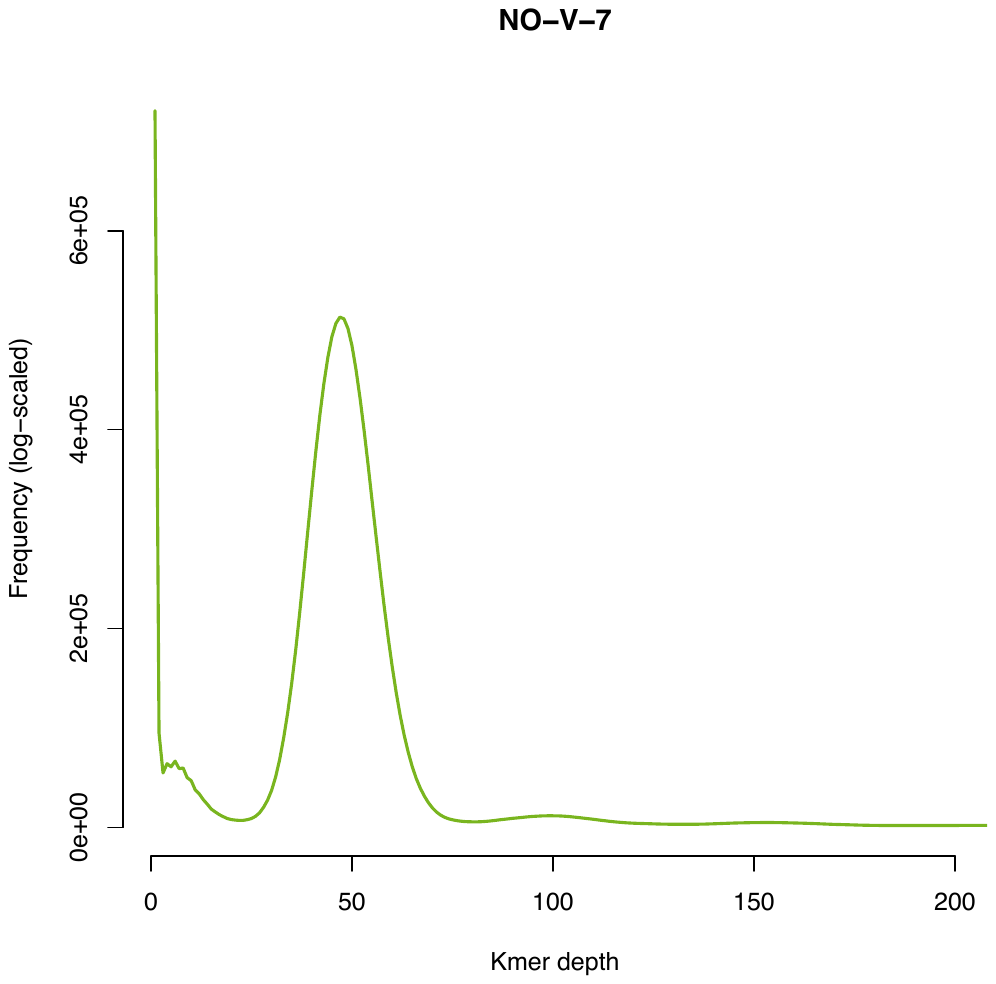

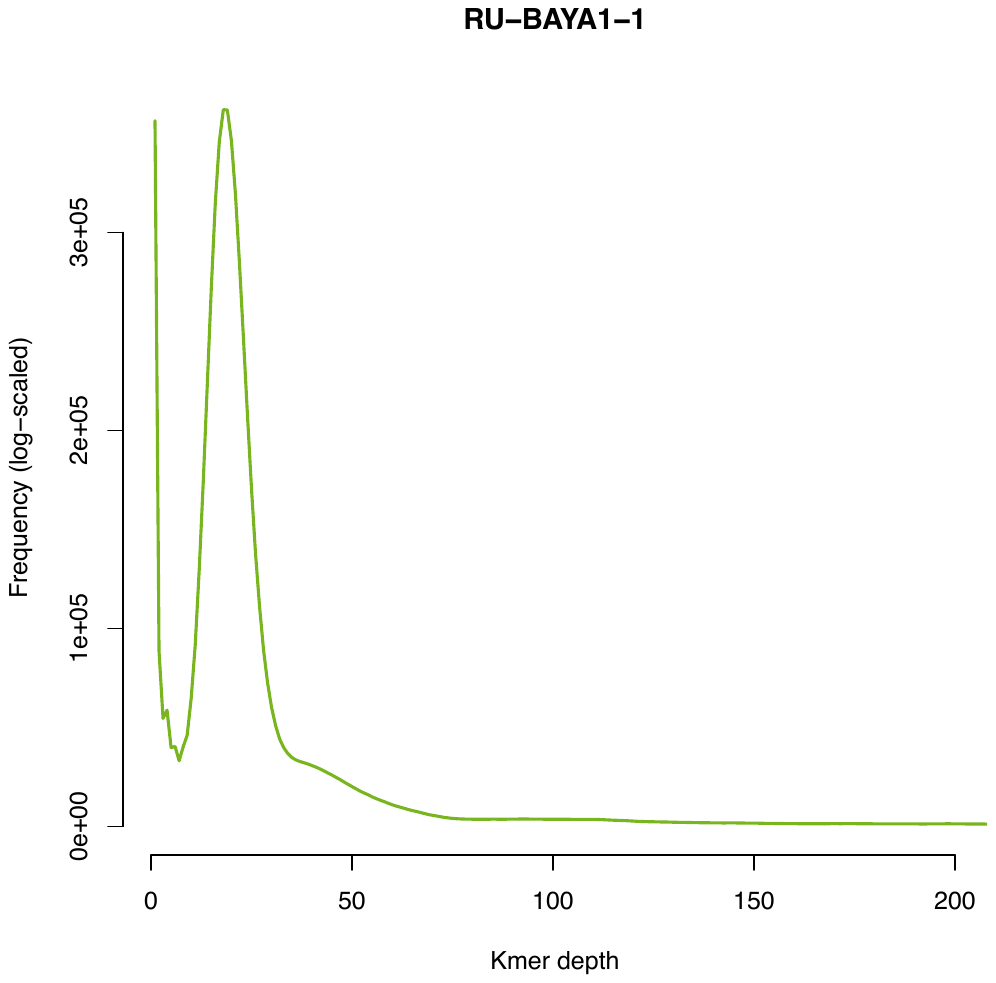

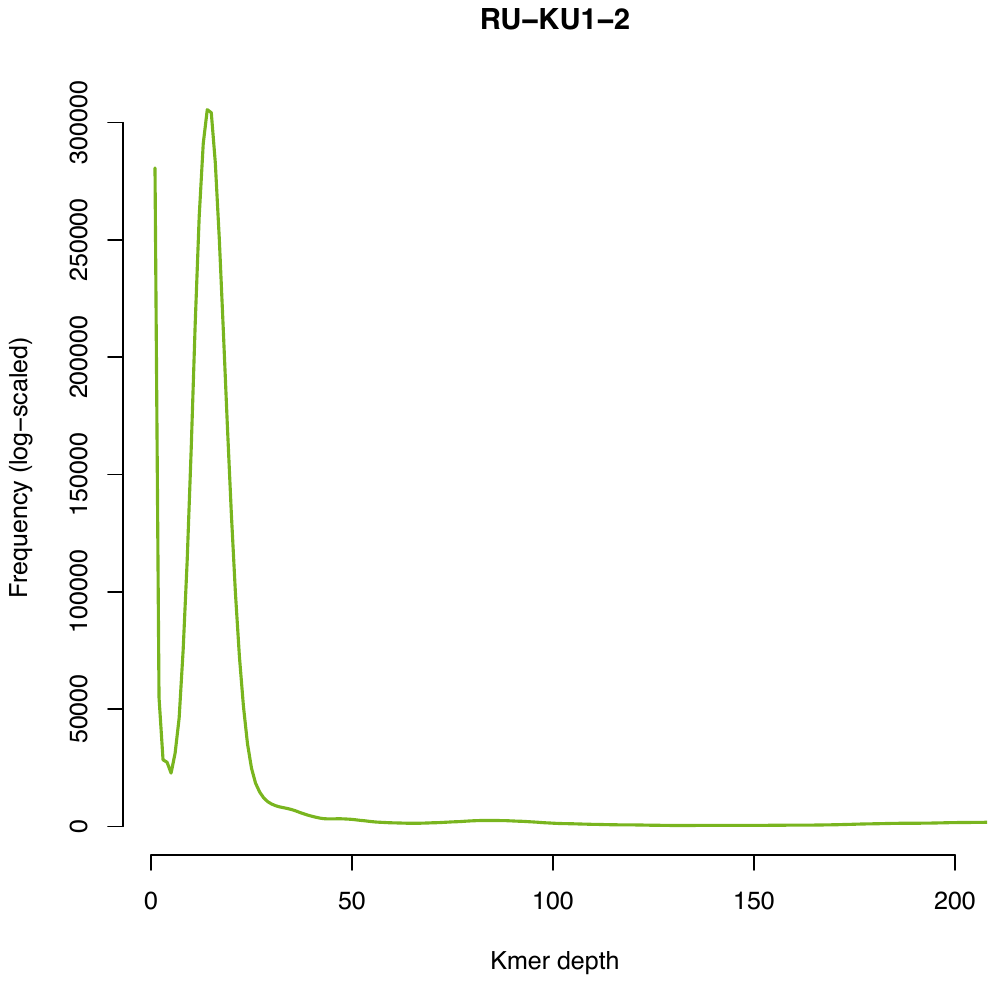

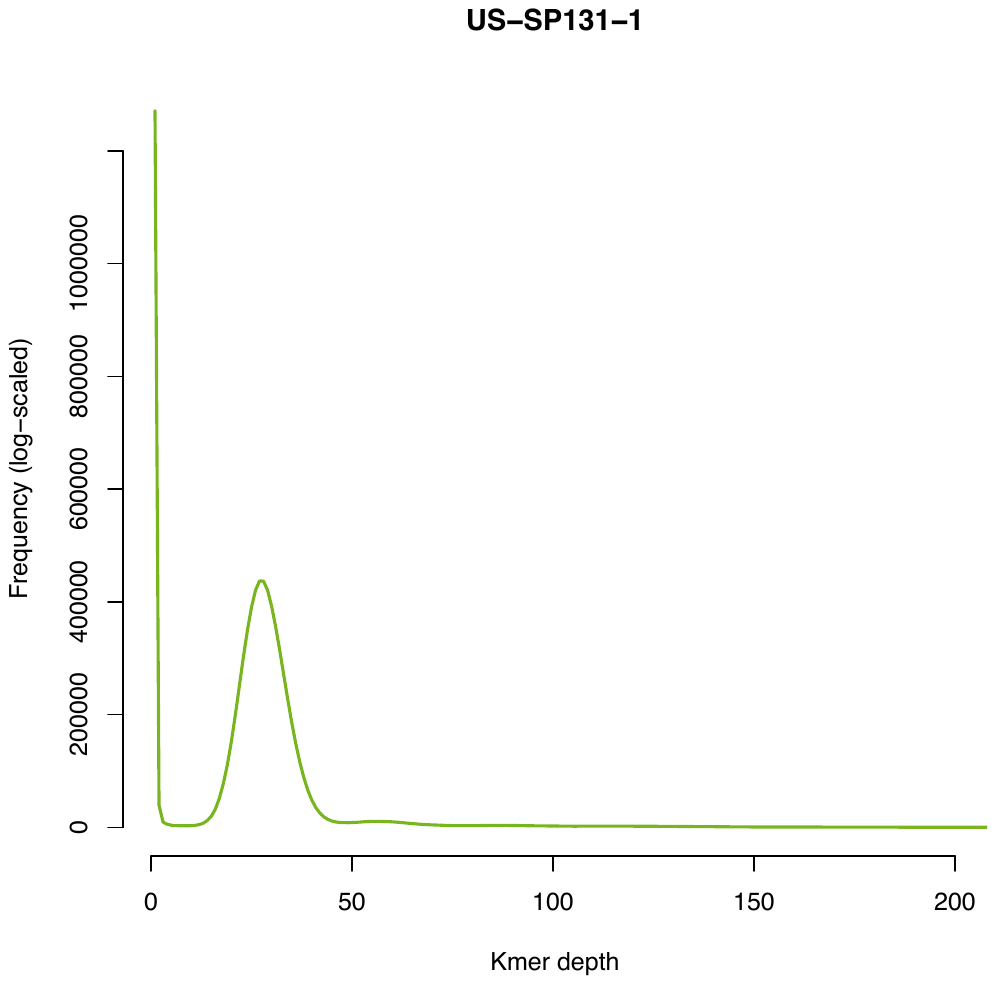

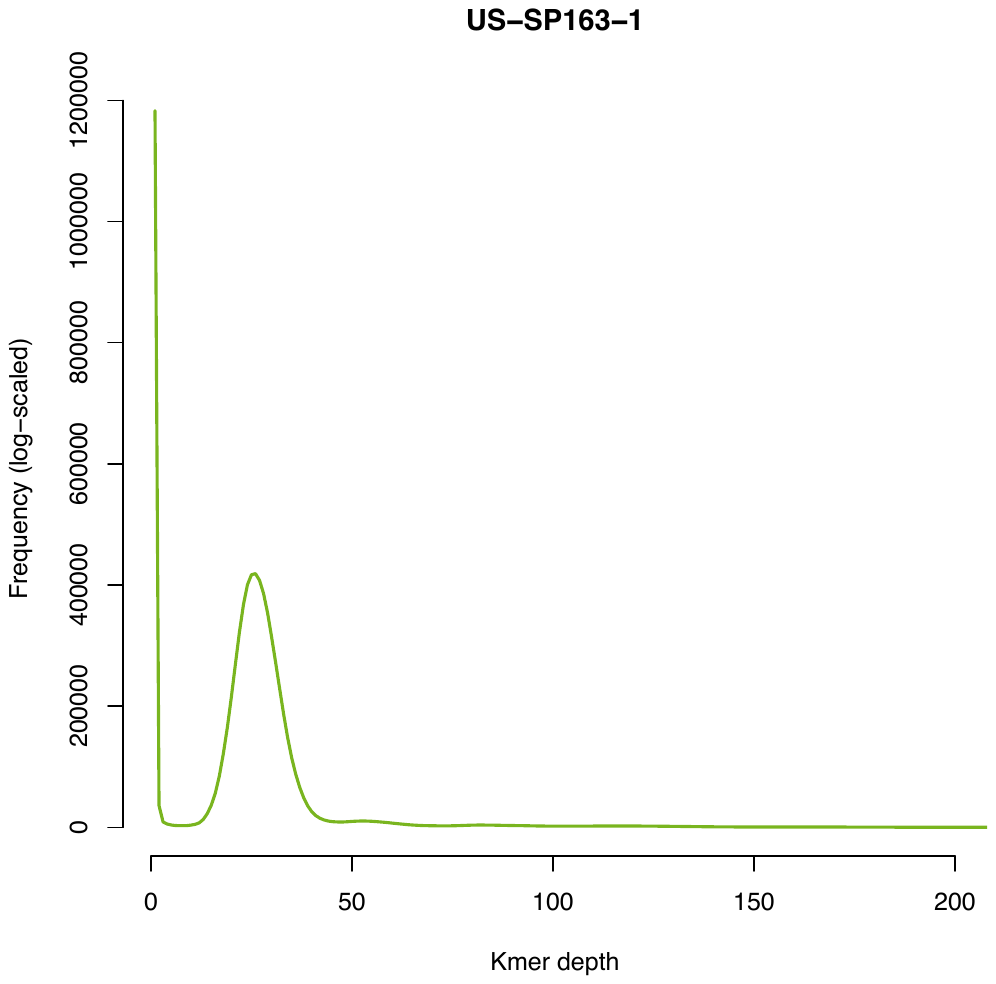

### Supplemental Figure 3

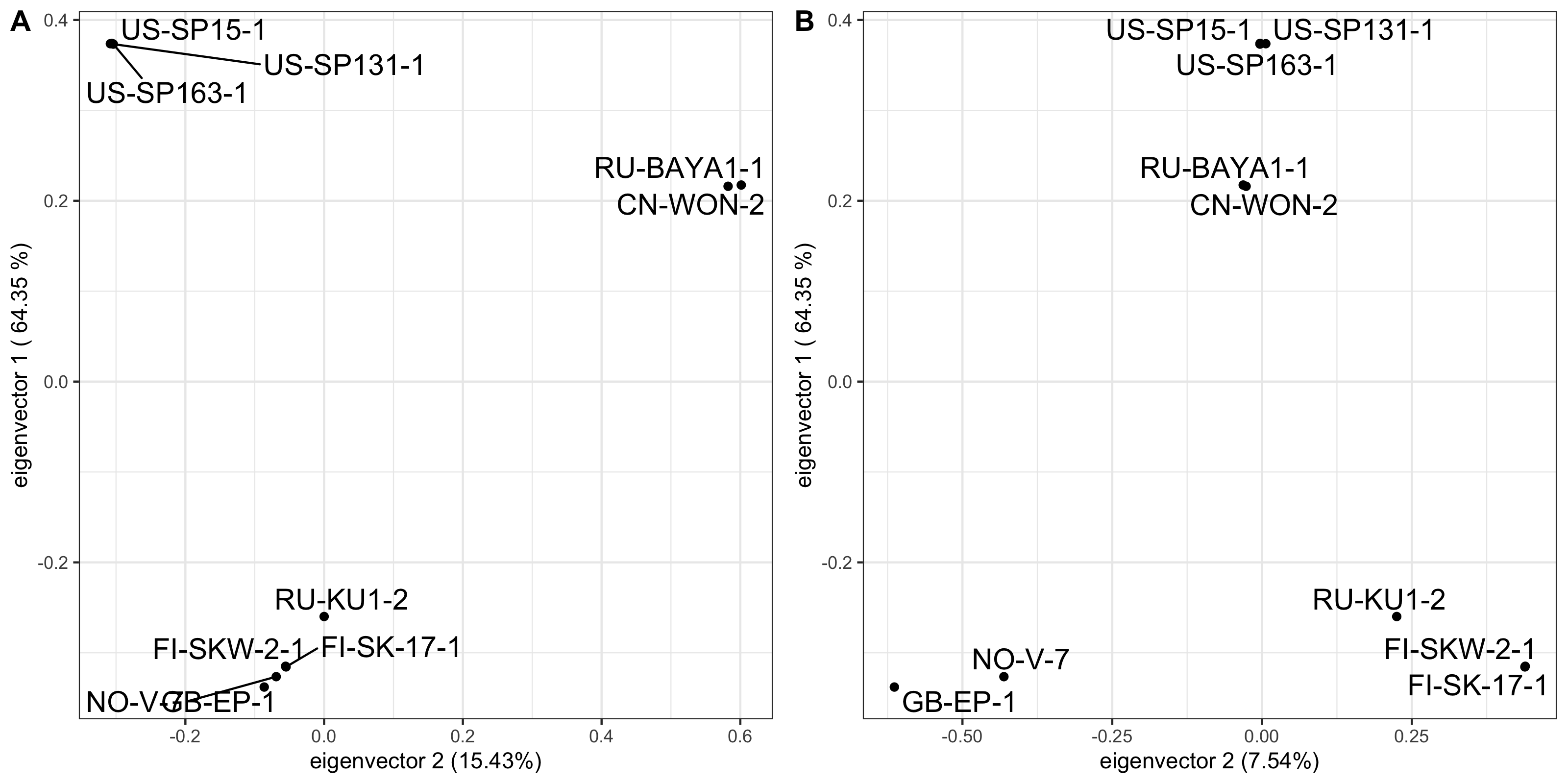

### Supplemental Figure 4

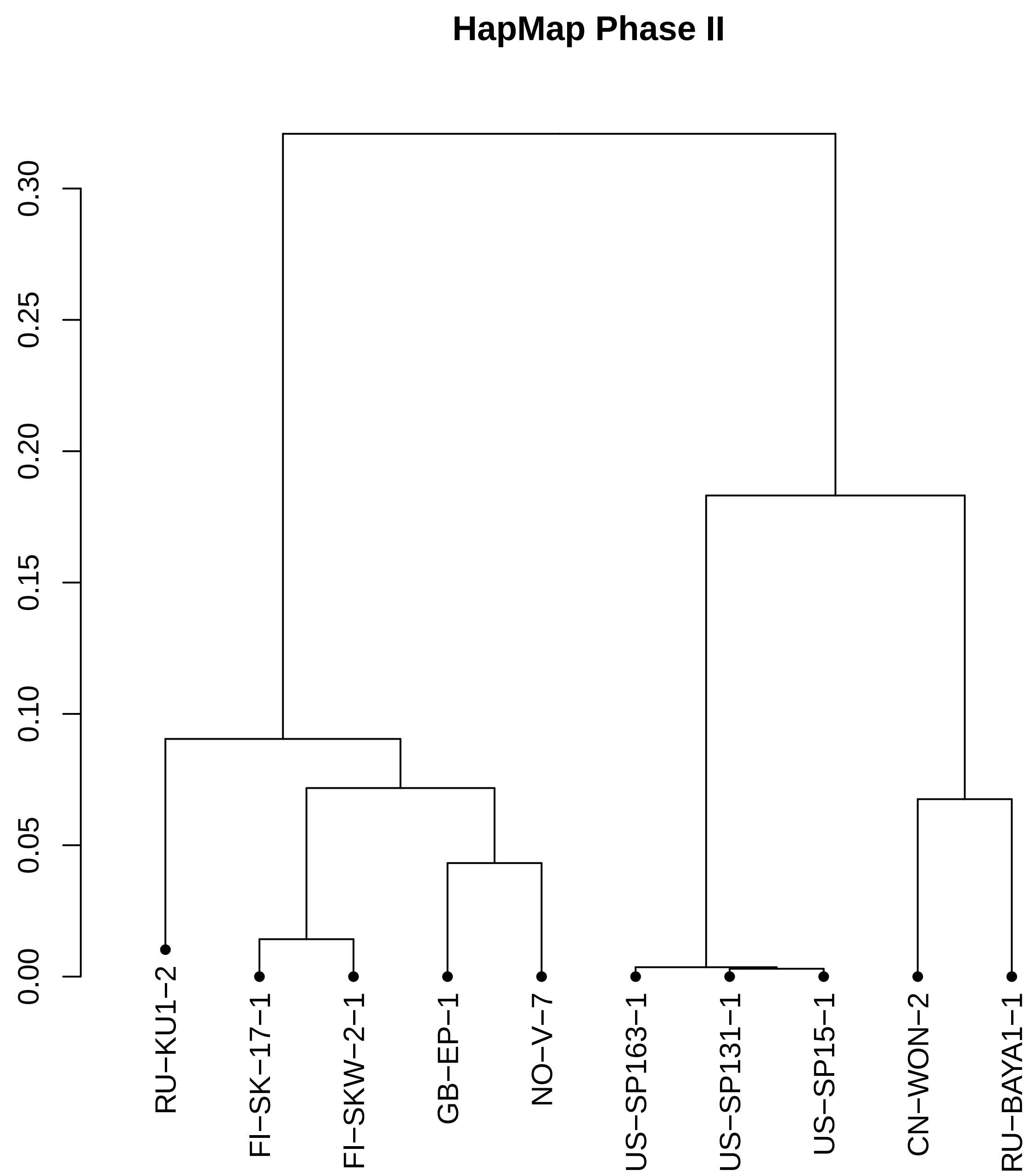

### Supplemental Figure 5

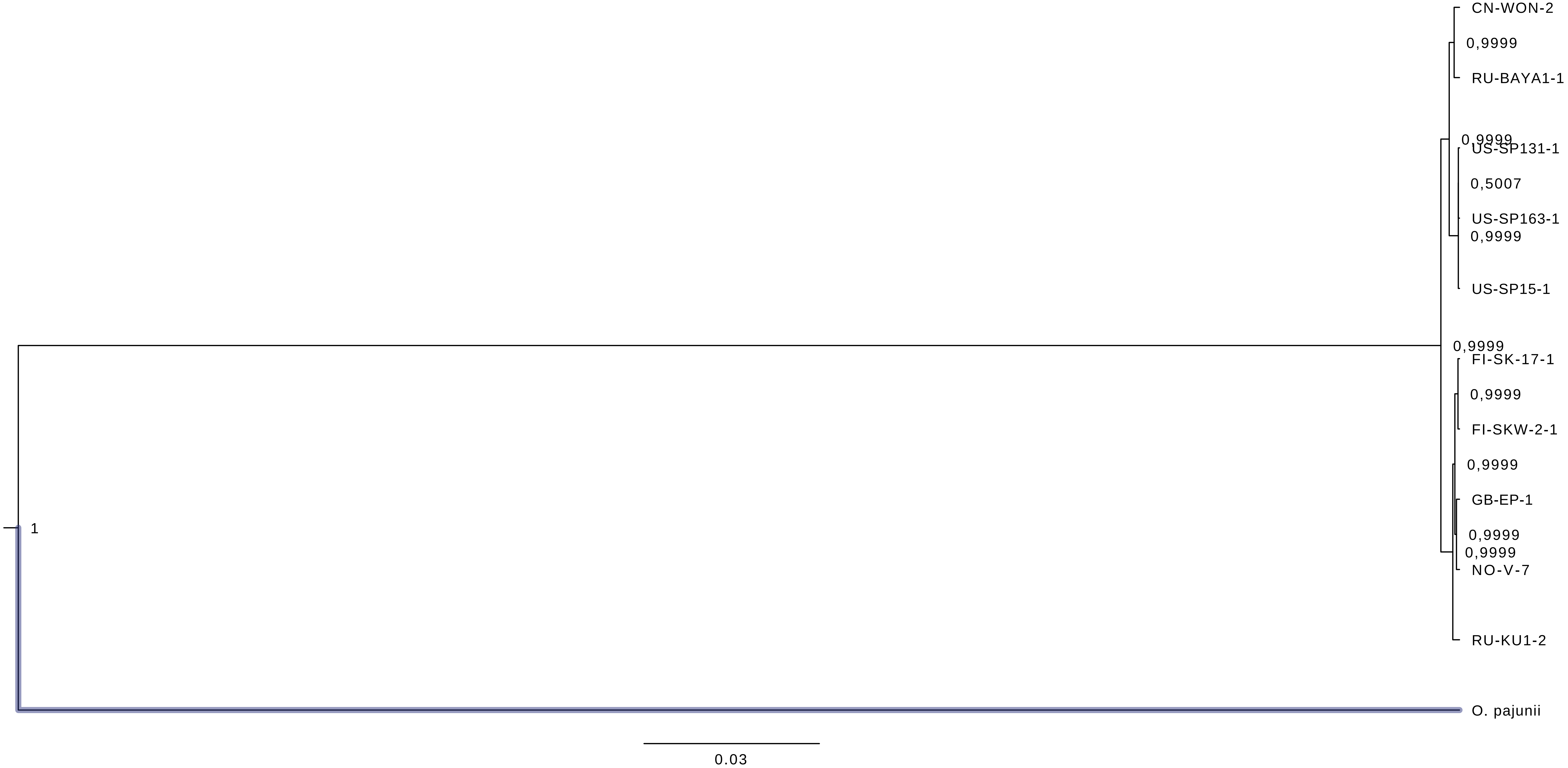
