## Supplemental Table 1 for "Population genetic analysis reveals the role of natural selection and phylogeography on genome-wide diversity in an extremely compact and reduced microsporidian genome"

| **Clone-label** | **Microscopy suggested Infection** | **PCR suggested Infection** |
| --- | --- | --- |
| BE-OM-1 infected O.c.2 | Oc | ? |
| BE-KN2-1 infected G.I. | Gi | Md |
| BE-WH1-2 infected G.i. | Gi | Gi |
| CH-H-1 infected G.i. | Gi | Md |
| CH-Z-1 infected G.i. | Gi | Md |
| CN-WON-1 | Oc | Oc |
| CN-WON-3 | Oc | Oc |
| CN-XIU-1 | Md | Md |
| CN-YUE-1 | Md | Md |
| CZ-N2-1 infected G.i.2 | Gi, Oc | Gi |
| DE-G1-106 infected G.i. | Gi | Md |
| DE-K-35-9 infected G.i. | Gi, Oc | Gi |
| DE-R1-1 infected G.i. | Gi | Gi |
| DE-R1-1 infected M.d. | Md | Md |
| DE-S2-1 infected G.i | Gi | Md |
| DE-S3-3infected G.i | Gi | Md |
| EG-ELIAS-1 G.i. | Gi | Md |
| ES-DO1-1 infected O.c. | Oc, Ht | Gi |
| FI-SK-58-2 | Ht | NA |
| GB-EK1-1 | Md | Md |
| GB-EK1-32 infected G.i. | Gi | Md |
| IE-DUB-1 infected O.c. | Oc | Gi |
| IL-BN-2 | Hm | NA |
| IL-EY-17 infected O.c. | Oc | Gi |
| IL-YERU-16 infected O.c. | Oc | Gi |
| IR-GG-102 | Hm | NA |
| IT-ISR3-2 infected | Oc | Gi |
| IT-PER-2 infected O.c. | Oc | Gi |
| IT-PER-2 | Ht | NA |
| IT-PER-10 | Ht | NA |
| RU-AST2-1 | Gi | NA |
| RU-HA4-2 G.i. | Gi | Md |
| RU-NE1-34 O.c. | Oc | Oc |
| US-ConnecticutValley M.d. | Md | Md |
