## Supplemental Table 2 for "Population genetic analysis reveals the role of natural selection and phylogeography on genome-wide diversity in an extremely compact and reduced microsporidian genome"

| **Locus_tag** | **π** | **π_N_/π_S_** | **Ortholog** | **Annotation** |
| --- | --- | --- | --- | --- |
| M896_010050 | 0.018745939 | 0.85449963 | Cg_02C_49 | hypothetical protein |
| M896_010080 | 0 | Inf | Cg_01_4 | hypothetical protein |
| M896_010090 | 0 | Inf | Cg_01_6 | hypothetical protein |
| M896_010100 | 0.001939479 | 1.10132974 | Cg_01_7 | fructose-1%2C6-bisphosphate aldolase |
| M896_010110 | 0.001010101 | Inf | Cg_01_8 | hypothetical protein |
| M896_010120 | 0.001341922 | Inf | Cg_01_9 | dynein light chain |
| M896_010130 | 0.001410039 | 1.67706168 | Cg_01_10 | hypothetical protein |
| M896_010140 | 0.001786183 | 0.7054238 | Cg_01_11 | hypothetical protein |
| M896_010150 | 0.004151404 | 0.054145 | Cg_01_12 | hypothetical protein |
| M896_010160 | 0 | Inf | Cg_01_13 | ribosomal protein L2P |
| M896_010180 | 0 | Inf | Cg_01_14 | U3 small nucleolar ribonucleoprotein |
| M896_010190 | 0.004188713 | 1.25639364 | Cg_01_15 | hypothetical protein |
| M896_010200 | 0.002078435 | 1.4731405 | Cg_01_16 | DEAD-like helicase |
| M896_010210 | 0.002092488 | Inf | Cg_01_17 | ribosome biogenesis protein BRX1 |
| M896_010220 | 0.002316406 | 0.06993267 | Cg_01_18 | zuotin-like protein |
| M896_010230 | 0.001986044 | 0.27600555 | Cg_01_19 | cell differentiation protein Rcd1 |
| M896_010240 | 0 | Inf | Cg_01_20 | hypothetical protein |
| M896_010250 | 0.000335008 | 0 | Cg_01_21 | hypothetical protein |
| M896_010270 | 0.001202501 | Inf | Cg_01_22 | hypothetical protein |
| M896_010280 | 0 | Inf | Cg_01_23 | hypothetical protein |
| M896_010290 | 0.000408998 | 0 | Cg_01_24 | cleavage and polyadenylation specificity factor |
| M896_010300 | 0.004299404 | Inf | Cg_01_25 | hypothetical protein |
| M896_010310 | 0.00428027 | 2.11268212 | Cg_01_26 | DNA repair protein Rad4 |
| M896_010320 | 0 | Inf | Cg_01_27 | actin |
| M896_010330 | 0 | Inf | Cg_01_28 | forkhead/HNF3 transcription factor |
| M896_010340 | 0.00355209 | 0.10344828 | Cg_01_29 | ribosomal biogenesis protein |
| M896_010350 | 0.003064108 | 1.47857781 | Cg_01_30 | hypothetical protein |
| M896_010360 | 0 | Inf | Cg_01_31 | RNA polymerase II transcriptional regulation |
| M896_010370 | 0.004803241 | 0.62137428 | Cg_01_32 | Fe-S cluster assembly protein IscU |
| M896_010380 | 0.002480308 | 0.4859561 | Cg_01_33 | hypothetical protein |
| M896_010390 | 0.002608242 | 0 | Cg_01_34 | hypothetical protein |
| M896_010400 | 0.003779634 | 0.78539249 | Cg_01_35 | peripheral Golgi membrane protein |
| M896_010410 | 0.002244669 | Inf | Cg_01_36 | putative RNA-binding protein |
| M896_010420 | 0.004008715 | 0.25395567 | Cg_01_37 | small nuclear ribonucleoprotein |
| M896_010430 | 0.006518519 | 1.09413384 | Cg_01_39 | hypothetical protein |
| M896_010440 | 0.002741324 | Inf | Cg_01_40 | hypothetical protein |
| M896_010450 | 0.001496321 | 0.15625141 | Cg_01_42 | hypothetical protein |
| M896_010460 | 0.004644516 | 0.4113428 | Cg_01_43 | hypothetical protein |
| M896_010470 | 0.00331675 | 2.12546125 | Cg_01_44 | hypothetical protein |
| M896_010480 | 0.00379295 | 0.10511826 | Cg_01_45 | hypothetical protein |
| M896_010490 | 0.004472397 | 1.50461636 | Cg_01_46 | DNA-directed RNA polymerase I |
| M896_010500 | 0.003118908 | 1.07469627 | Cg_01_47 | hypothetical protein |
| M896_010510 | 0.001161946 | Inf | Cg_01_48 | hypothetical protein |
| M896_010520 | 0.005359276 | 1.41994849 | Cg_01_49 | serine/threonine protein kinase |
| M896_010530 | 0.002658238 | 0.98038902 | Cg_01_51 | hypothetical protein |
| M896_010540 | 0.002974186 | 0.12404593 | Cg_01_52 | phosphoacetylglucosamine mutase |
| M896_010550 | 0.001698284 | 0.23423818 | Cg_01_53 | subunit of RNA polymerase II transcription factor |
| M896_010560 | 0 | Inf | Cg_01_54 | thioredoxin reductase-like protein |
| M896_010570 | 0.002539683 | 0 | Cg_01_55 | hypothetical protein |
| M896_010580 | 0.004786862 | 0.1450719 | Cg_01_56 | eukaryotic translation initiation factor 2 |
| M896_010590 | 0.002552365 | 0.68646106 | Cg_01_57 | hypothetical protein |
| M896_010600 | 0.000823045 | 0 | Cg_01_58 | thioredoxin |
| M896_010610 | 0.001068153 | 0 | Cg_01_59 | uridine kinase |
| M896_010630 | 0.000979816 | 0.1410386 | Cg_01_60 | thymidine kinase |
| M896_010640 | 0.001763668 | Inf | Cg_01_61 | HRD ubiquitin ligase complex protein |
| M896_010650 | 0.001654081 | Inf | Cg_01_62 | hypothetical protein |
| M896_010670 | 0.002737928 | 0.59051736 | Cg_01_63 | RING Zn-finger domain-containing protein |
| M896_010680 | 0.002079912 | Inf | Cg_01_64 | hypothetical protein |
| M896_010690 | 0.001129944 | Inf | Cg_01_65 | trehalose-6-phosphate synthase |
| M896_010700 | 0.001358025 | Inf | Cg_01_66 | hypothetical protein |
| M896_010710 | 0.00048011 | Inf | Cg_01_67 | hypothetical protein |
| M896_010720 | 0.003040034 | 1.23050006 | Cg_01_68 | CCCH-type Zn-finger protein |
| M896_010730 | 0.004148148 | 2.18560075 | Cg_01_69 | putative ribonucleoprotein |
| M896_010740 | 0.001377018 | Inf | Cg_01_70 | hypothetical protein |
| M896_010750 | 0.002411714 | Inf | Cg_01_71 | hypothetical protein |
| M896_010760 | 0.003400627 | 0.24071869 | Cg_01_72 | trehalose 6-phosphate phosphatase |
| M896_010770 | 0 | Inf | Cg_01_73 | hypothetical protein |
| M896_010780 | 0.001706776 | Inf | Cg_01_74 | hypothetical protein |
| M896_010790 | 0.002533103 | Inf | Cg_01_75 | hypothetical protein |
| M896_010800 | 0.002325581 | 0.33647059 | Cg_01_76 | hypothetical protein |
| M896_010810 | 0.001435544 | Inf | Cg_01_77 | ribosomal protein S12 |
| M896_010820 | 0.00242963 | 0.4886231 | Cg_01_78 | thioredoxin-like protein |
| M896_010830 | 0.006990089 | 3.15791091 | Cg_01_79 | ubiquitin conjugating enzyme E2 |
| M896_010840 | 0.001115573 | Inf | Cg_01_80 | hypothetical protein |
| M896_010850 | 0.002104377 | Inf | Cg_01_81 | hypothetical protein |
| M896_010860 | 0 | Inf | Cg_01_82 | Tat binding protein 1-interacting protein |
| M896_010870 | 0.002093967 | 0.19454274 | Cg_01_83 | aldo-keto reductase |
| M896_010880 | 0.002204586 | Inf | Cg_01_84 | hypothetical protein |
| M896_010890 | 0.003574074 | 2.44253578 | Cg_01_85 | WD domain-containing protein |
| M896_010900 | 0.001966623 | 2.09894662 | Cg_01_86 | hypothetical protein |
| M896_010910 | 0.000793651 | 0 | Cg_01_87 | ubiquitin-conjugating enzyme E2 |
| M896_010920 | 0.003036753 | 0.17901694 | Cg_01_88 | hypothetical protein |
| M896_010930 | 0.001088629 | Inf | Cg_01_89 | cell cycle control microtubule-binding protein |
| M896_010940 | 0.006901311 | 0.24484536 | Cg_01_90 | general negative regulator of transcription |
| M896_010950 | 0.001724476 | 0.13979232 | Cg_01_91 | hypothetical protein |
| M896_010960 | 0.001296017 | 0 | Cg_01_92 | Rad25-like DNA repair helicase |
| M896_010970 | 0.001285304 | 0.50242175 | Cg_01_94 | hypothetical protein |
| M896_010980 | 0.001462304 | 0.10264648 | Cg_01_95 | tRNA/rRNA cytosine-C5-methylase |
| M896_010990 | 0.001721551 | 2.54682884 | Cg_01_96 | hypothetical protein |
| M896_011000 | 0 | Inf | Cg_08A_2 | SCF ubiquitin ligase and anaphase-promoting complex protein |
| M896_011010 | 0.004910053 | 1.38436482 | Cg_01_97 | hypothetical protein |
| M896_011030 | 0.001332549 | 0.67364662 | Cg_01_98 | hypothetical protein |
| M896_011040 | 0.010642827 | Inf | Cg_01_99 | cytochrome B5 |
| M896_011050 | 0.000575004 | 0.7301511 | Cg_01_100 | hypothetical protein |
| M896_011060 | 0.001343747 | 0.56708075 | Cg_01_101 | S8 serine protease |
| M896_011070 | 0.002226631 | Inf | Cg_01_102 | hypothetical protein |
| M896_011080 | 0.002080732 | 0.25009754 | Cg_01_103 | cyclin K-like protein |
| M896_011090 | 0.002943232 | 1.77162301 | Cg_01_104 | chromosome segregation ATPase |
| M896_011100 | 0.001538863 | 0.80761068 | Cg_01_105 | hypothetical protein |
| M896_011110 | 0.00130593 | Inf | Cg_01_106 | large subunit of replication factor C |
| M896_011120 | 0.001308273 | 0.44743429 | Cg_01_107 | hypothetical protein |
| M896_011130 | 0.000945626 | 1.00854857 | Cg_01_108 | hypothetical protein |
| M896_011140 | 0.002041706 | 0.18702264 | Cg_01_110 | dynamin |
| M896_011150 | 0.001304932 | 0 | Cg_01_111 | guanylate kinase |
| M896_011160 | 0.001312224 | 0 | Cg_01_112 | AAA+ ATPase |
| M896_011170 | 0.00130552 | 0.15074788 | Cg_01_113 | hypothetical protein |
| M896_011180 | 0.001109515 | Inf | Cg_01_114 | hypothetical protein |
| M896_011190 | 0.003703704 | 0.82164634 | Cg_01_115 | hypothetical protein |
| M896_011200 | 0.002008788 | 0.30274779 | Cg_01_116 | hypothetical protein |
| M896_011210 | 0.004084967 | 0 | Cg_01_117 | nuclear distribution C-like protein |
| M896_011220 | 0.00318734 | 0.30058828 | Cg_01_118 | WD40 domain-containing protein |
| M896_011230 | 0.003268543 | 0.33030686 | Cg_01_119 | WD40 domain-containing protein |
| M896_011240 | 0 | Inf | Cg_01_120 | frataxin |
| M896_011250 | 0.002877026 | 0.78961309 | Cg_01_121 | Rho-associated coiled-coil domain-containing protein |
| M896_011260 | 0.003843996 | 0.55141551 | Cg_01_122 | hypothetical protein |
| M896_011270 | 0.004224537 | 0.625 | Cg_01_123 | subunit of transcription initiation factor TFIID |
| M896_011280 | 0.00390193 | 0.68674287 | Cg_01_124 | hypothetical protein |
| M896_011290 | 0 | Inf | Cg_01_125 | hypothetical protein |
| M896_011300 | 0.004995693 | 0.29479374 | Cg_01_126 | putative RNA-binding protein |
| M896_011310 | 0.001230969 | Inf | Cg_01_127 | septin |
| M896_011320 | 0.008354866 | Inf | Cg_01_128 | hypothetical protein |
| M896_011330 | 0.00285658 | 3.4852768 | Cg_01_129 | chitin synthase |
| M896_011340 | 0 | Inf | Cg_01_130 | hypothetical protein |
| M896_011350 | 0.00139237 | 0.26599683 | Cg_01_131 | putative serine/threonine protein kinase |
| M896_011360 | 0 | Inf | Cg_01_132 | hypothetical protein |
| M896_011370 | 0.00239899 | 0.90475551 | Cg_01_133 | hypothetical protein |
| M896_011380 | 0.001987353 | 0.96621647 | Cg_01_134 | putative Cdc48 ATPase |
| M896_011390 | 0.001432981 | Inf | Cg_01_135 | subunit of pre-mRNA cleavage GTPase |
| M896_011400 | 0.002373247 | Inf | Cg_01_136 | tyrosinyl-tRNA synthetase |
| M896_011410 | 0.002257572 | 0.65058808 | Cg_01_137 | hypothetical protein |
| M896_011420 | 0.002754036 | Inf | Cg_01_138 | hypothetical protein |
| M896_011430 | 0.002469136 | Inf | Cg_01_140 | putative proteasome regulatory complex protein |
| M896_011440 | 0.000757576 | Inf | Cg_01_141 | hypothetical protein |
| M896_011450 | 0.00409636 | 0.43985994 | Cg_01_142 | putative RIO kinase |
| M896_011460 | 0.003276353 | 1.17750901 | Cg_01_143 | RNA pol Rpb4 domain-containing protein |
| M896_011470 | 0.000877461 | Inf | Cg_01_144 | hypothetical protein |
| M896_011480 | 0.001508916 | 0.15600364 | Cg_01_145 | hypothetical protein |
| M896_011490 | 0.002572802 | 0.09987333 | Cg_01_146 | putative proliferating cell nuclear antigen |
| M896_011500 | 0.001394091 | 0.61201781 | Cg_01_147 | putative centromere/microtubule binding protein |
| M896_011510 | 0.002493108 | 0.97502812 | Cg_01_148 | WD G-beta repeat domain-containing protein |
| M896_011520 | 0.002122122 | 0.15831407 | Cg_01_149 | ADP-ribosylation factor family domain-containing protein |
| M896_011530 | 0.002751323 | 0.18122644 | Cg_01_150 | type-B DNA-directed DNA polymerase |
| M896_011540 | 0.003760684 | 1.69570557 | Cg_01_151 | subunit 10 of anaphase-promoting complex |
| M896_011550 | 0.002844281 | 0.08844973 | Cg_01_152 | hypothetical protein |
| M896_011560 | 0.001203704 | 0.88978826 | Cg_01_153 | putative SAM dependent methyltransferase |
| M896_011570 | 0.001730038 | 0.11754634 | Cg_01_154 | putative superfamily II RNA helicase |
| M896_011580 | 0.002523047 | 0.67199931 | Cg_01_155 | phosphatidylinositol transfer protein |
| M896_011590 | 0.004587542 | 0.38894797 | Cg_01_156 | ribosomal protein S9 |
| M896_011600 | 0.002614379 | Inf | Cg_01_157 | small nuclear ribonucleoprotein |
| M896_011610 | 0.005705706 | Inf | Cg_01_158 | ribosomal protein L21 |
| M896_011620 | 0.007520786 | 1.78087997 | Cg_01_159 | hypothetical protein |
| M896_011630 | 0.002402402 | Inf | Cg_01_160 | SecE/Sec61-gamma subunit of protein translocation complex |
| M896_011650 | 0.004746788 | 0.3402747 | Cg_01_161 | hypothetical protein |
| M896_011660 | 0.000705467 | Inf | Cg_01_162 | HAM1 domain-containing protein |
| M896_011670 | 0.002558635 | 0.57333016 | Cg_01_163 | 6-phosphogluconate dehydrogenase |
| M896_011680 | 0.003703704 | 0.1283164 | Cg_01_164 | hypothetical protein |
| M896_011690 | 0.001458151 | Inf | Cg_01_165 | PX domain-containing protein |
| M896_011700 | 0.003802733 | 0.246482 | Cg_01_166 | hypothetical protein |
| M896_011710 | 0.000750117 | Inf | Cg_01_167 | hypothetical protein |
| M896_011720 | 0.0018107 | 0.08248233 | Cg_01_168 | serine/threonine protein phosphatase 2A-like protein |
| M896_011730 | 0.002539683 | 0.16925422 | Cg_01_169 | putative GTPase |
| M896_011740 | 0.001711672 | 0.04708031 | Cg_01_171 | hypothetical protein |
| M896_011750 | 0.001730703 | 0 | Cg_01_172 | hypothetical protein |
| M896_011760 | 0.002651515 | 0.10824321 | Cg_01_173 | minichromosome maintenance protein |
| M896_011770 | 0.00331384 | 0.08689655 | Cg_01_174 | hypothetical protein |
| M896_011780 | 0.000746714 | 0 | Cg_01_175 | peptidase C48 domain-containing protein |
| M896_011790 | 0.007726692 | 0.90826483 | Cg_01_176 | hypothetical protein |
| M896_011800 | 0.003977611 | 0.12328915 | Cg_01_177 | hypothetical protein |
| M896_011810 | 0.001229314 | 0 | Cg_01_178 | serine palmitoyltransferase |
| M896_011820 | 0.003534392 | 0.1894303 | Cg_01_179 | hypothetical protein |
| M896_011830 | 0.00210887 | 0.86473191 | Cg_01_180 | RhoGAP domain-containing protein |
| M896_011840 | 0.001858788 | 0.10797392 | Cg_01_181 | hypothetical protein |
| M896_011850 | 0.001361807 | 0.25876578 | Cg_01_182 | putative membrane protein |
| M896_011860 | 0.004004711 | Inf | Cg_01_183 | putative pseudouridine synthase |
| M896_011870 | 0.001561365 | 0.18042913 | Cg_01_184 | ribosomal protein S6e |
| M896_011880 | 0.000877655 | Inf | Cg_01_185 | hypothetical protein |
| M896_011890 | 0.002011696 | Inf | Cg_01_186 | glucose-6-phosphate isomerase |
| M896_011900 | 0.004277517 | Inf | Cg_01_187 | hypothetical protein |
| M896_011910 | 0.003441595 | 3.43073342 | Cg_01_188 | putative serine/threonine kinase |
| M896_011920 | 0 | Inf | Cg_01_189 | Ran GTPase-activating protein |
| M896_011930 | 0.004523608 | 0.71771351 | Cg_01_190 | hypothetical protein |
| M896_011940 | 0.000335008 | Inf | Cg_01_191 | putative ribosomal protein L1 |
| M896_011960 | 0 | Inf | Cg_01_192 | hypothetical protein |
| M896_011970 | 0.003472222 | 0.70340796 | Cg_01_193 | putative amino acid transporter |
| M896_011980 | 0.003892028 | 0.1875635 | Cg_01_194 | hypothetical protein |
| M896_011990 | 0.001082954 | 0.44444444 | Cg_01_195 | Fcf1 domain-containing protein |
| M896_012000 | 0.003844901 | 0.53945753 | Cg_01_196 | hypothetical protein |
| M896_012010 | 0.003000686 | 0.36648091 | Cg_01_197 | FAT domain-containing protein |
| M896_012020 | 0.00617284 | Inf | Cg_01_198 | hypothetical protein |
| M896_012030 | 0.003779912 | 0.17087064 | Cg_01_199 | hypothetical protein |
| M896_012040 | 0.003753086 | 0.14084459 | Cg_01_200 | hypothetical protein |
| M896_012050 | 0.000642479 | 0 | Cg_01_201 | putative transcription initiation factor TFIIIB |
| M896_012060 | 0.002153316 | Inf | Cg_01_202 | hypothetical protein |
| M896_012070 | 0.00280713 | 1.59236889 | Cg_01_203 | hypothetical protein |
| M896_012080 | 0.003724166 | 0.76611437 | Cg_01_204 | ribosomal protein L10-like protein |
| M896_012090 | 0.007215007 | 0.48609865 | Cg_01_205 | hypothetical protein |
| M896_012100 | 0.002604803 | Inf | Cg_01_206 | putative phosphoesterase |
| M896_012110 | 0.001489533 | Inf | Cg_01_207 | subunit C of CCAAT-binding factor |
| M896_012120 | 0.002621929 | 1.16381788 | Cg_01_208 | calcineurin-like phosphoesterase |
| M896_012130 | 0.004675398 | 0.68594541 | Cg_01_209 | hypothetical protein |
| M896_012140 | 0.001932367 | Inf | Cg_01_210 | hypothetical protein |
| M896_012150 | 0.001046244 | 0.55310522 | Cg_01_211 | hypothetical protein |
| M896_012160 | 0.003903305 | Inf | Cg_01_212 | putative GTPase |
| M896_012170 | 0.003130511 | Inf | Cg_01_213 | putative subunit E of vacuolar ATP synthase |
| M896_012180 | 0.003246274 | 0.84061396 | Cg_01_214 | hypothetical protein |
| M896_012190 | 0.00201788 | 0.40929513 | Cg_01_215 | beta-CASP domain-containing protein |
| M896_012200 | 0.003828548 | 0.07998347 | Cg_01_216 | hypothetical protein |
| M896_012210 | 0.002344116 | Inf | Cg_01_217 | hypothetical protein |
| M896_012220 | 0.002830992 | 0.45363525 | Cg_01_218 | eukaryotic peptide chain release factor eRF1 |
| M896_012230 | 0.002292769 | 0.24328445 | Cg_01_219 | subunit RSC8 of RSC chromatin remodeling complex |
| M896_012240 | 0.004004004 | 0.22561649 | Cg_01_220 | hypothetical protein |
| M896_012250 | 0.00653046 | 0.33631234 | Cg_01_221 | DNA polymerase alpha/epsilon-like protein |
| M896_012260 | 0.002866763 | 0.55277567 | Cg_01_222 | phosphoglycerate kinase |
| M896_012270 | 0.002262713 | 0.12306085 | Cg_01_223 | acetyl-CoA synthetase |
| M896_012280 | 0.002275225 | 0.54023621 | Cg_01_224 | putative DNA mismatch repair protein |
| M896_012290 | 0.00456621 | 1.48472849 | Cg_01_225 | beta type-4 subunit of proteasome |
| M896_012300 | 0 | Inf | Cg_01_226 | hypothetical protein |
| M896_012310 | 0.005372405 | 0.57260008 | Cg_01_227 | subunit Rpb11 of RNA polymerase II |
| M896_012320 | 0.002159594 | 0.16097527 | Cg_01_228 | glycerol 3 phosphate dehydrogenase |
| M896_012330 | 0.001333333 | 0.21150662 | Cg_01_229 | phosphomannomutase |
| M896_012340 | 0.000510856 | 0 | Cg_01_230 | ribosomal protein S3AE |
| M896_012350 | 0.002664324 | 0.44835523 | Cg_01_231 | putative nitric-oxide synthase |
| M896_012360 | 0.004719764 | 0.03512364 | Cg_01_232 | hypothetical protein |
| M896_012370 | 0.002554278 | 0.38788231 | Cg_01_233 | WD40 domain-containing protein |
| M896_012380 | 0.003483682 | 0.54597084 | Cg_01_235 | hypothetical protein |
| M896_012390 | 0.004553734 | Inf | Cg_01_236 | hypothetical protein |
| M896_012400 | 0.00071817 | 0.11654908 | Cg_01_237 | hypothetical protein |
| M896_012410 | 0.003367003 | Inf | Cg_01_238 | protein phosphatase inhibitor |
| M896_012420 | 0.003037687 | 0.62382626 | Cg_01_240 | hypothetical protein |
| M896_012430 | 0.002880658 | 0.62722689 | Cg_01_241 | hypothetical protein |
| M896_012440 | 0.002301994 | 0.74834356 | Cg_01_242 | hypothetical protein |
| M896_012450 | 0.00203828 | 0.08524519 | Cg_01_243 | hypothetical protein |
| M896_012460 | 0.002270884 | 2.17280164 | Cg_01_244 | hypothetical protein |
| M896_012470 | 0.001095462 | 0 | Cg_01_245 | hypothetical protein |
| M896_012480 | 0.001563031 | 0.31595031 | Cg_01_246 | putative Sar1 GTPase |
| M896_012500 | 0.00222475 | 0.72859528 | Cg_01_247 | hypothetical protein |
| M896_012510 | 0.001884253 | 1.90905218 | Cg_01_248 | kinesin motor domain-containing protein |
| M896_012520 | 0.007267645 | 0.85092913 | Cg_01_250 | hypothetical protein |
| M896_020020 | 0.001302932 | Inf | Cg_02A_1 | hypothetical protein |
| M896_020030 | 0.001725178 | 0.98563066 | Cg_02A_2 | hypothetical protein |
| M896_020040 | 0.000938967 | Inf | Cg_02A_3 | hypothetical protein |
| M896_020050 | 0.000814625 | Inf | Cg_02A_4 | aminotransferase |
| M896_020060 | 0.002259887 | 1.29421719 | Cg_02A_5 | thioredoxin domain-containing protein |
| M896_020070 | 0.000857055 | Inf | Cg_02A_6 | hypothetical protein |
| M896_020080 | 0.00316358 | Inf | Cg_02A_7 | hypothetical protein |
| M896_020090 | 0.003420167 | 1.9242766 | Cg_02A_8 | hypothetical protein |
| M896_020110 | 0.001058201 | Inf | Cg_02B_1 | ribosomal protein L6 |
| M896_020120 | 0.003524229 | 0.06099234 | Cg_02B_2 | beta subunit of transcription initiation factor IIF |
| M896_020130 | 0.002987862 | 0 | Cg_02B_3 | ribosomal protein S17 |
| M896_020140 | 0.002962963 | 0 | Cg_02B_4 | S26 type I signal peptidase |
| M896_020150 | 0.005163818 | 0.69874052 | Cg_02B_5 | ribosomal protein L7a |
| M896_020160 | 0 | Inf | Cg_02B_6 | ubiquitin |
| M896_020170 | 0.001364522 | 0.08122931 | Cg_02B_8 | poly(A) polymerase |
| M896_020180 | 0.001364522 | Inf | Cg_02B_9 | putative histone-like transcription factor |
| M896_020190 | 0.000846922 | Inf | Cg_02B_10 | hypothetical protein |
| M896_020210 | 0.000914495 | Inf | Cg_02B_11 | hypothetical protein |
| M896_020220 | 0.001119105 | 0 | Cg_02B_12 | HscB-like chaperone |
| M896_020230 | 0.002515202 | 0.32415793 | Cg_02B_13 | hypothetical protein |
| M896_020240 | 0.002769864 | 0.79524564 | Cg_02B_14 | dead box helicase |
| M896_020250 | 0.00130527 | 0.37249795 | Cg_02B_15 | hypothetical protein |
| M896_020260 | 0.001062733 | Inf | Cg_02B_16 | hypothetical protein |
| M896_020270 | 0.001022431 | Inf | Cg_02B_17 | hypothetical protein |
| M896_020280 | 0.001431493 | 0.69494389 | Cg_02B_18 | hypothetical protein |
| M896_020290 | 0.001929012 | Inf | Cg_02B_19 | hypothetical protein |
| M896_020300 | 0 | Inf | Cg_02B_20 | ribosomal protein L5 |
| M896_020310 | 0.002923977 | 0.13333333 | Cg_02B_21 | hypothetical protein |
| M896_020320 | 0.001952695 | 0.31286979 | Cg_02B_22 | hypothetical protein |
| M896_020330 | 0.002746567 | 0.31752447 | Cg_02B_23 | hydroxyacylglutathione hydrolase |
| M896_020340 | 0.003311735 | 2.42754934 | Cg_02B_24 | hypothetical protein |
| M896_020350 | 0.002498282 | 0.44778049 | Cg_02B_26 | hypothetical protein |
| M896_020360 | 0.001757444 | 0.2524731 | Cg_02B_27 | catalytic protein kinase |
| M896_020370 | 0.00138001 | Inf | Cg_02B_28 | hypothetical protein |
| M896_020390 | 0.005072464 | 0.43454385 | Cg_02B_29 | hypothetical protein |
| M896_020410 | 0.00134959 | 0 | Cg_02B_30 | subunit delta of TCP-1 chaperonin |
| M896_020420 | 0.003003003 | 0.20060904 | Cg_02B_31 | mitogen-activated protein kinase |
| M896_020430 | 0.000758808 | 0 | Cg_02B_32 | hypothetical protein |
| M896_020440 | 0.001661475 | Inf | Cg_02B_33 | hypothetical protein |
| M896_020450 | 0.002283557 | 0.03217879 | Cg_02B_34 | 26S proteasome regulatory complex protein |
| M896_020460 | 0.001218535 | 0.16074627 | Cg_02B_35 | hypothetical protein |
| M896_020470 | 0.001223242 | 1.19732137 | Cg_02B_36 | nucleoporin autopeptidase domain-containing protein |
| M896_020480 | 0.002079272 | 0 | Cg_02B_37 | gamma subunit of transcription initiation factor IIA |
| M896_020490 | 0.001023123 | 0.4095334 | Cg_02B_38 | glutathione peroxidase |
| M896_020500 | 0.001219326 | 0.09015589 | Cg_02B_39 | Sec1-like intracellular trafficking protein |
| M896_020510 | 0 | Inf | Cg_02B_41 | hypothetical protein |
| M896_020520 | 0.001460616 | 0 | Cg_02B_42 | Ras-like GTP binding protein |
| M896_020530 | 0.001040129 | 0.06894734 | Cg_02B_43 | hypothetical protein |
| M896_020540 | 0.000794786 | Inf | Cg_02B_44 | hypothetical protein |
| M896_020550 | 0.002425201 | 0.09078422 | Cg_02B_45 | hypothetical protein |
| M896_020560 | 0.00225802 | 0.27966594 | Cg_02B_47 | hypothetical protein |
| M896_020570 | 0.001271979 | 0.80658138 | Cg_02B_48 | beta type-3 subunit of proteasome |
| M896_020580 | 0.001560758 | 2.61579864 | Cg_02B_49 | putative E1-E2 ATPase |
| M896_020590 | 0.000882585 | Inf | Cg_02B_50 | hypothetical protein |
| M896_020610 | 0.004542278 | 1.11639377 | Cg_02B_51 | hypothetical protein |
| M896_020620 | 0 | Inf | Cg_02B_52 | small subunit of replication factor C |
| M896_020630 | 0.002383253 | 0.36582654 | Cg_02B_53 | hypothetical protein |
| M896_020640 | 0.001252885 | 0.18056655 | Cg_02B_54 | hypothetical protein |
| M896_020650 | 0.001143425 | 0.46940559 | Cg_02B_55 | GATA zinc finger domain-containing protein |
| M896_020660 | 0.003794778 | 0.41067285 | Cg_02B_56 | putative RNA-binding protein |
| M896_020670 | 0.002904145 | 0.10488197 | Cg_02B_57 | Sof1 domain-containing U3 snoRNP protein |
| M896_020680 | 0.000524934 | 0 | Cg_02B_58 | hypothetical protein |
| M896_020690 | 0.001362282 | 0 | Cg_02B_59 | hypothetical protein |
| M896_020700 | 0.002331265 | Inf | Cg_02B_60 | hypothetical protein |
| M896_020710 | 0.00114899 | 0.03937497 | Cg_02B_61 | Smp2-like plasmid maintenance protein |
| M896_020730 | 0.006466466 | 0.28326446 | Cg_02B_63 | hypothetical protein |
| M896_020740 | 0 | Inf | Cg_02B_65 | hypothetical protein |
| M896_020750 | 0.001410935 | Inf | Cg_02B_66 | hypothetical protein |
| M896_020760 | 0.000861326 | Inf | Cg_02B_67 | putative Ran GTPase binding protein |
| M896_020770 | 0.004276441 | Inf | Cg_02B_68 | hypothetical protein |
| M896_020780 | 0.002717003 | 0.87216806 | Cg_02B_69 | hypothetical protein |
| M896_020790 | 0.004454626 | 0.32999905 | Cg_02B_70 | hypothetical protein |
| M896_020800 | 0.001619626 | 0.35015252 | Cg_02B_71 | Hsp70-like protein |
| M896_020840 | 0.003240741 | 0.6440703 | Cg_02C_54 | ribosomal protein S8 |
| M896_020860 | 0.000653595 | Inf | Cg_02C_56 | ribosomal protein L35Ae |
| M896_020870 | 0.00505698 | Inf | Cg_02C_57 | hypothetical protein |
| M896_020880 | 0.002948222 | 0.59245018 | Cg_02C_58 | hypothetical protein |
| M896_020890 | 0.001328751 | 0.10105195 | Cg_02C_59 | Suf domain-containing protein |
| M896_020900 | 0.001016703 | Inf | Cg_02C_60 | thioredoxin reductase |
| M896_020910 | 0 | Inf | Cg_02C_61 | hypothetical protein |
| M896_020920 | 0.00136876 | Inf | Cg_02C_62 | Emp24/gp25L domain-containing protein |
| M896_020930 | 0.0004329 | 0 | Cg_02C_63 | transport protein particle complex protein |
| M896_020940 | 0.004884595 | Inf | Cg_02C_64 | hypothetical protein |
| M896_020950 | 0.002642844 | 0.46128492 | Cg_02C_65 | hypothetical protein |
| M896_020960 | 0.001642772 | 1.34502924 | Cg_02C_66 | hypothetical protein |
| M896_020970 | 0.003544214 | 0.29260116 | Cg_02C_67 | hypothetical protein |
| M896_020980 | 0.004004004 | Inf | Cg_02C_68 | LSM domain-containing protein |
| M896_020990 | 0.000650329 | 0.39879121 | Cg_02C_69 | hypothetical protein |
| M896_021000 | 0.002204586 | Inf | Cg_02C_70 | hypothetical protein |
| M896_021020 | 0 | Inf | Cg_02C_71 | putative nascent polypeptide-associated complex protein |
| M896_021030 | 0.001676677 | Inf | Cg_02C_72 | subunit alpha of protein prenyltransferase |
| M896_021040 | 0.002382118 | 0.86590853 | Cg_02C_73 | hypothetical protein |
| M896_021050 | 0 | Inf | Cg_02C_74 | ubiquitin domain-containing protein |
| M896_021060 | 0.002210275 | 0.94133632 | Cg_02C_75 | Rad3-like DNA-binding helicase |
| M896_021070 | 0.001436008 | Inf | Cg_02C_76 | heat shock protein 90 |
| M896_021080 | 0.00303757 | 0.91521659 | Cg_02C_77 | hypothetical protein |
| M896_021090 | 0.001960077 | 0.18870268 | Cg_02C_78 | protein kinase domain-containing protein |
| M896_021100 | 0.001143118 | Inf | Cg_02C_79 | microsomal signal peptidase |
| M896_021110 | 0.001380328 | 0.09116657 | Cg_02C_80 | DNA replication licensing factor |
| M896_021120 | 0.000935279 | Inf | Cg_02C_81 | hypothetical protein |
| M896_021130 | 0.002763385 | Inf | Cg_02C_82 | nuclear transport factor 2 |
| M896_021140 | 0.002429943 | 0.52634722 | Cg_02C_83 | hypothetical protein |
| M896_021150 | 0.00337132 | 0.14209539 | Cg_02C_84 | hypothetical protein |
| M896_021160 | 0.012913799 | 1.78067583 | Cg_02C_85 | homeobox domain-containing protein |
| M896_021170 | 0.001339906 | 0.40966864 | Cg_02C_86 | tRNA synthetase class I |
| M896_021180 | 0.002380952 | Inf | Cg_02C_87 | DNA ligase |
| M896_021190 | 0.001642315 | 2.20377907 | Cg_02C_88 | Sec23/Sec24-like protein |
| M896_021200 | 0.002121583 | 0.90472382 | Cg_02C_89 | chromosome condensation complex Condensin |
| M896_021210 | 0.004269547 | 0.34653795 | Cg_02C_90 | transcription initiation factor TFIID |
| M896_021220 | 0.0004004 | Inf | Cg_02C_91 | hypothetical protein |
| M896_021230 | 0.001169591 | Inf | Cg_02C_92 | mitosis protein Dim1 |
| M896_021240 | 0.005934343 | 0.40806975 | Cg_02C_93 | hypothetical protein |
| M896_021250 | 0.0018107 | 0.75900621 | Cg_02C_94 | hypothetical protein |
| M896_021260 | 0.001959379 | 4.43704438 | Cg_02C_95 | dolichyl-phosphate-mannose-protein |
| M896_021270 | 0.001583468 | 0.31862857 | Cg_02C_96 | hypothetical protein |
| M896_021280 | 0.001424501 | 2.55077479 | Cg_02C_97 | hypothetical protein |
| M896_021290 | 0.001725998 | Inf | Cg_02C_98 | hypothetical protein |
| M896_021300 | 0.001146601 | 1.54505819 | Cg_02C_99 | hypothetical protein |
| M896_021310 | 0 | Inf | Cg_02C_100 | transcription factor E2F |
| M896_021320 | 0.001827802 | Inf | Cg_02C_101 | prolyl-tRNA synthetase |
| M896_021330 | 0.002136526 | 0.66665362 | Cg_02C_102 | trehalase |
| M896_021340 | 0.003024213 | 0.55431973 | Cg_02C_103 | peptidase M48 domain-containing protein |
| M896_021350 | 0.005334692 | 0.27218973 | Cg_02C_104 | hypothetical protein |
| M896_021360 | 0 | Inf | Cg_02C_105 | hypothetical protein |
| M896_021370 | 0.003550182 | 0.76717358 | Cg_02C_106 | hypothetical protein |
| M896_021380 | 0.002015504 | 0.16683854 | Cg_02C_107 | subunit A of RAB protein geranylgeranyltransferase |
| M896_021390 | 0.001851852 | Inf | Cg_02C_108 | hypothetical protein |
| M896_021400 | 0.000474074 | Inf | Cg_02C_109 | hypothetical protein |
| M896_021410 | 0.002162162 | 0.22339473 | Cg_02C_110 | hypothetical protein |
| M896_021420 | 0.004302652 | 0.29710459 | Cg_02C_111 | serine/threonine kinase |
| M896_021430 | 0.001957071 | 0.47691321 | Cg_02C_112 | Sec23 domain-containing protein |
| M896_021440 | 0.003433897 | 0 | Cg_02C_113 | prenylated RAB acceptor 1 |
| M896_021450 | 0.002616747 | 0.4118451 | Cg_02C_114 | hypothetical protein |
| M896_021470 | 0.002296469 | 0.13573342 | Cg_02C_115 | alanyl-tRNA synthetase |
| M896_021480 | 0.003809524 | 0.31643681 | Cg_02C_117 | subunit A of transcription initiation factor TFIID |
| M896_021490 | 0.0039675 | 0.24585153 | Cg_02C_118 | hypothetical protein |
| M896_021500 | 0.002175476 | 0.1073574 | Cg_02C_119 | putative DEAD-box helicase |
| M896_021510 | 0.003708972 | 0.41818705 | Cg_02C_120 | hypothetical protein |
| M896_021520 | 0.00043573 | Inf | Cg_02C_121 | putative Per1-like membrane protein |
| M896_021550 | 0.001270879 | 0 | Cg_02C_124 | ATP-dependent RNA helicase |
| M896_021560 | 0.002113776 | 0.07677599 | Cg_02C_125 | glycosyltransferase |
| M896_021570 | 0.001351717 | Inf | Cg_02C_126 | Ca2+-binding protein |
| M896_021580 | 0.002063492 | Inf | Cg_02C_127 | hypothetical protein |
| M896_021590 | 0.001169591 | Inf | Cg_02C_128 | RNA-binding protein |
| M896_021600 | 0.002176584 | 0.1684744 | Cg_02C_129 | ribosomal protein S3 |
| M896_021610 | 0.001294498 | 1.24410616 | Cg_02C_130 | RIO protein kinase |
| M896_021620 | 0.001975309 | 0.68408473 | Cg_02C_131 | hypothetical protein |
| M896_021630 | 0.004601571 | Inf | Cg_02C_132 | ribosomal protein S28 |
| M896_021640 | 0.004383081 | 1.37181773 | Cg_02C_133 | hypothetical protein |
| M896_021650 | 0.002926383 | 1.41867846 | Cg_02C_134 | hypothetical protein |
| M896_021660 | 0.006070732 | 0.21252856 | Cg_02C_135 | DNA repair protein Mms21 |
| M896_021670 | 0.002460222 | 0.10195448 | Cg_02C_136 | histone acetyltransferase |
| M896_021680 | 0.003292181 | 0.84983515 | Cg_02C_137 | endochitinase |
| M896_021690 | 0.002617458 | Inf | Cg_02C_138 | C-terminal domain of replication factor C |
| M896_021700 | 0.001786183 | 0 | Cg_02C_139 | hypothetical protein |
| M896_021710 | 0.001998824 | Inf | Cg_02C_140 | ribosomal protein S15A |
| M896_021720 | 0.001102984 | 0.71422951 | Cg_02C_141 | minichromosome maintenance protein |
| M896_021730 | 0.002502503 | Inf | Cg_02C_142 | hypothetical protein |
| M896_021740 | 0.004032922 | Inf | Cg_02C_143 | glutaredoxin |
| M896_021750 | 0.000861326 | 0.29261759 | Cg_02C_144 | ribosome stability and mRNA decay protein |
| M896_021760 | 0.001815182 | Inf | Cg_02C_145 | DNA helicase TIP49 |
| M896_021780 | 0.000513067 | Inf | Cg_02C_146 | hypothetical protein |
| M896_021790 | 0.004487475 | 1.52558852 | Cg_02C_147 | hypothetical protein |
| M896_021800 | 0.002324037 | 1.48912188 | Cg_02C_148 | P-type ATPase |
| M896_021810 | 0.002394314 | Inf | Cg_02C_149 | Rab GTPase |
| M896_021820 | 0.001026495 | Inf | Cg_02C_150 | hypothetical protein |
| M896_021830 | 0 | Inf | Cg_02C_151 | hypothetical protein |
| M896_021840 | 0.000255428 | Inf | Cg_02C_153 | hypothetical protein |
| M896_021850 | 0 | Inf | Cg_02C_154 | hypothetical protein |
| M896_021860 | 0 | Inf | Cg_02C_155 | hypothetical protein |
| M896_021870 | 0.002203857 | Inf | Cg_02C_156 | hypothetical protein |
| M896_021880 | 0.000798403 | 0.25354462 | Cg_02C_157 | hypothetical protein |
| M896_030080 | 0.0043409 | 0.68284812 | Cg_03_105 | chromatin organization modifier protein |
| M896_030090 | 0.002469136 | Inf | Cg_03_106 | hypothetical protein |
| M896_030100 | 0.002893519 | Inf | Cg_03_107 | hypothetical protein |
| M896_030110 | 0.001373737 | Inf | Cg_03_108 | nucleotide sugar transporter |
| M896_030120 | 0.005196241 | Inf | Cg_03_109 | hypothetical protein |
| M896_030130 | 0.001722653 | Inf | Cg_03_110 | putative transcription initiation factor IIF auxiliary protein |
| M896_030140 | 0.002741467 | Inf | Cg_03_111 | ankyrin-like protein |
| M896_030150 | 0.010441292 | 0.42946165 | Cg_03_112 | hypothetical protein |
| M896_030160 | 0.001080247 | Inf | Cg_03_113 | Ca2+-binding protein |
| M896_030170 | 0.002153316 | Inf | Cg_03_114 | Rho-like GTPase |
| M896_030180 | 0.003122731 | Inf | Cg_03_115 | hypothetical protein |
| M896_030190 | 0.00292583 | 1.97664328 | Cg_03_116 | Snf2-like DNA/RNA helicase |
| M896_030200 | 0.003233912 | Inf | Cg_03_117 | WD40 domain-containing protein |
| M896_030210 | 0 | Inf | Cg_03_118 | subunit B of CCAAT-binding transcription factor |
| M896_030220 | 0.000904722 | Inf | Cg_03_119 | hypothetical protein |
| M896_030230 | 0.004320988 | 0.48911652 | Cg_03_120 | hypothetical protein |
| M896_030240 | 0.002889703 | 0.84044028 | Cg_03_121 | ribosomal protein L18 |
| M896_030250 | 0.003186908 | Inf | Cg_03_122 | hypothetical protein |
| M896_030260 | 0.003328645 | 0.44813293 | Cg_03_123 | hypothetical protein |
| M896_030270 | 0.00047619 | Inf | Cg_03_124 | histone H3 |
| M896_030280 | 0.001731948 | Inf | Cg_03_125 | hypothetical protein |
| M896_030290 | 0.001614885 | 0.679398 | Cg_03_126 | tRNA nucleotidyltransferase/poly(A) polymerase |
| M896_030300 | 0.002939447 | Inf | Cg_03_127 | Rab GTPase |
| M896_030310 | 0.002341201 | 1.5542255 | Cg_03_128 | hypothetical protein |
| M896_030320 | 0.001562692 | Inf | Cg_03_129 | Cdc48-like ATPase |
| M896_030330 | 0.000296956 | 0.2529844 | Cg_03_130 | hypothetical protein |
| M896_030350 | 0.000675858 | Inf | Cg_03_131 | putative sugar kinase |
| M896_030360 | 0.000993413 | 0.29918737 | Cg_03_132 | PelA-like RNA-binding protein |
| M896_030370 | 0.000534188 | 0.61572652 | Cg_03_133 | histone deacetylase |
| M896_030380 | 0.001839588 | 0.63313028 | Cg_03_134 | Myg1-like protein |
| M896_030390 | 0.001570101 | 3.91324622 | Cg_03_135 | hypothetical protein |
| M896_030400 | 0.00220817 | Inf | Cg_03_136 | regulatory subunit of ATP-dependent 26S proteasome |
| M896_030420 | 0.003025561 | 0.56817533 | Cg_03_137 | hypothetical protein |
| M896_030430 | 0.003139856 | Inf | Cg_03_138 | molybdopterin biosynthesis protein MoeB |
| M896_030440 | 0.00139693 | Inf | Cg_03_139 | hypothetical protein |
| M896_030450 | 0.001786711 | Inf | Cg_03_140 | cytidylate kinase |
| M896_030460 | 0 | Inf | Cg_03_141 | hypothetical protein |
| M896_030470 | 0.000646465 | Inf | Cg_03_142 | hypothetical protein |
| M896_030480 | 0.00229455 | Inf | Cg_03_143 | adenine nucleotide alpha hydrolase |
| M896_030490 | 0.000613196 | Inf | Cg_03_144 | hypothetical protein |
| M896_030510 | 0.00326147 | 0.3248074 | Cg_03_1 | hypothetical protein |
| M896_030520 | 0.002527233 | 0 | Cg_03_2 | peroxiredoxin |
| M896_030530 | 0.004098584 | 0.32321933 | Cg_03_3 | peptidase S8-like protein |
| M896_030550 | 0.003132648 | 0.06923077 | Cg_03_4 | homeodomain-containing transcription factor |
| M896_030560 | 0.001033592 | 0.15866655 | Cg_03_5 | Sac-like phosphoinositide polyphosphatase |
| M896_030570 | 0.002040364 | Inf | Cg_03_6 | calponin |
| M896_030580 | 0.002595758 | Inf | Cg_03_7 | hypothetical protein |
| M896_030590 | 0.001760176 | 0.54443667 | Cg_03_8 | DNA repair Rad51-like protein |
| M896_030600 | 0.002308802 | 0.16 | Cg_03_9 | putative subunit of V-type ATP synthase |
| M896_030610 | 0 | Inf | Cg_03_10 | hypothetical protein |
| M896_030620 | 0.003271077 | 0.54530664 | Cg_03_11 | N-myristoyl transferase |
| M896_030630 | 0.002645503 | 0.27556185 | Cg_03_12 | hypothetical protein |
| M896_030640 | 0.003462812 | 0.21345807 | Cg_03_13 | phosphatidylinositol 4-kinase |
| M896_030660 | 0.001556811 | 0.14099574 | Cg_03_14 | hypothetical protein |
| M896_030670 | 0.001266774 | 0.19944006 | Cg_03_15 | flap endonuclease 1 |
| M896_030680 | 0 | Inf | Cg_03_16 | large chain of transcription initiation factor IIA |
| M896_030690 | 0.001101101 | 0.11605387 | Cg_03_18 | hypothetical protein |
| M896_030700 | 0.000659134 | 0 | Cg_03_19 | hypothetical protein |
| M896_030710 | 0 | Inf | Cg_03_20 | putative subunit of transcription initiation factor TFIIIB |
| M896_030720 | 0.001082954 | 0.24918831 | Cg_03_21 | Cdc6-like protein |
| M896_030730 | 0.000920447 | Inf | Cg_03_22 | hypothetical protein |
| M896_030740 | 0.001249092 | 0.19212188 | Cg_03_23 | 14-3-3-like signal transduction protein |
| M896_030750 | 0.000774107 | 0 | Cg_03_24 | hypothetical protein |
| M896_030760 | 0.001916681 | 2.46837349 | Cg_03_25 | SCP/PR1 domain-containing protein |
| M896_030770 | 0.002642713 | 0.06383632 | Cg_03_26 | serine/threonine protein kinase |
| M896_030780 | 0.003606663 | 1.01401049 | Cg_03_27 | hypothetical protein |
| M896_030790 | 0.001611653 | 0.18689785 | Cg_03_28 | hypothetical protein |
| M896_030800 | 0.002303143 | 0 | Cg_03_29 | ribosomal protein L7 |
| M896_030810 | 0.001768933 | 0.25764739 | Cg_03_30 | putative G2/M transition transcriptional factor |
| M896_030820 | 0.002354354 | 2.81597527 | Cg_03_31 | ubiquitin-activating enzyme E1 |
| M896_030830 | 0.002600473 | 0.08611919 | Cg_03_32 | putative tRNA-intron endonuclease |
| M896_030840 | 0.004038066 | 0.49900784 | Cg_03_33 | protein kinase |
| M896_030850 | 0.000995237 | 0.04071951 | Cg_03_34 | Sec1-like intracellular trafficking protein |
| M896_030860 | 0.002264511 | 0.35213329 | Cg_03_35 | haspin serine/threonine kinase |
| M896_030870 | 0.000893092 | 0.7331784 | Cg_03_36 | hypothetical protein |
| M896_030880 | 0.003220612 | 0.27110838 | Cg_03_37 | hypothetical protein |
| M896_030890 | 0.001638807 | 0 | Cg_03_39 | hypothetical protein |
| M896_030900 | 0.002126075 | 0.49933264 | Cg_03_40 | hypothetical protein |
| M896_030910 | 0.002962963 | 0.76388889 | Cg_03_41 | small nuclear ribonucleoprotein |
| M896_030920 | 0.002541757 | 0.59049959 | Cg_03_42 | hypothetical protein |
| M896_030930 | 0.0026046 | Inf | Cg_03_43 | beta-tubulin |
| M896_030940 | 0.004861111 | 0.18064601 | Cg_03_44 | putative sedlin |
| M896_030950 | 0.001247563 | Inf | Cg_03_45 | ubiquitin-related modifier-like protein |
| M896_030960 | 0.003449528 | 0.66042003 | Cg_03_46 | hypothetical protein |
| M896_030970 | 0.003227881 | 0.19663889 | Cg_03_47 | hypothetical protein |
| M896_030980 | 0.003327056 | 0.30386323 | Cg_03_48 | hypothetical protein |
| M896_030990 | 0 | Inf | Cg_03_49 | RNA-binding protein |
| M896_031000 | 0.002164502 | Inf | Cg_03_50 | RNA-binding protein |
| M896_031010 | 0.002180194 | 2.25525933 | Cg_03_51 | RNA binding domain-containing protein |
| M896_031020 | 0.001139601 | Inf | Cg_03_53 | ribosomal protein L34 |
| M896_031030 | 0.002220257 | 5.91549879 | Cg_03_54 | exoribonuclease R |
| M896_031050 | 0.001840234 | 2.0231777 | Cg_03_55 | ATP-dependent 6-phosphofructokinase |
| M896_031060 | 0.000819403 | Inf | Cg_03_56 | ribonuclease HII |
| M896_031070 | 0.001722995 | 0.516649 | Cg_03_57 | isopeptidase T |
| M896_031080 | 0.001481481 | 0.13137336 | Cg_03_58 | ribosomal protein S14 |
| M896_031090 | 0.001668335 | 0.12784091 | Cg_03_59 | hypothetical protein |
| M896_031100 | 0.000793651 | 0.20480405 | Cg_03_60 | protein kinase |
| M896_031110 | 0.001867414 | Inf | Cg_03_61 | homeodomain-containing transcription factor |
| M896_031120 | 0.001028807 | Inf | Cg_03_62 | Rab GTPase-interacting Golgi membrane protein |
| M896_031140 | 0.002580108 | 0.79170647 | Cg_03_64 | vacuolar import and degradation protein |
| M896_031150 | 0.001618837 | Inf | Cg_03_65 | putative GTPase-activating protein |
| M896_031160 | 0.00256021 | 3.87616822 | Cg_03_66 | Rho/Rac/Cdc42-like GTPase guanine nucleotide exchange factor |
| M896_031170 | 0.001456513 | 0.63482892 | Cg_03_67 | MutS-like mismatch repair ATPase |
| M896_031180 | 0.001765719 | 0 | Cg_03_68 | synaptobrevin/VAMP-like protein |
| M896_031190 | 0.001411456 | 0 | Cg_03_69 | heat shock protein 70 |
| M896_031200 | 0.002316081 | 1.78826605 | Cg_03_70 | hypothetical protein |
| M896_031210 | 0.002341551 | 0.98767814 | Cg_03_71 | subunit C of vacuolar H+-ATPase V1 |
| M896_031220 | 0 | Inf | Cg_03_72 | subunit M of DNA-directed RNA polymerase |
| M896_031230 | 0.003959935 | 1.07060145 | Cg_03_73 | subunit 2 of splicing factor 3a |
| M896_031240 | 0.001108893 | Inf | Cg_03_74 | ADP-ribosylation factor-like GTPase |
| M896_031250 | 0.000955043 | Inf | Cg_03_75 | sirtuin-like protein |
| M896_031260 | 0.002350427 | Inf | Cg_03_76 | hypothetical protein |
| M896_031270 | 0.000329048 | Inf | Cg_03_77 | subunit B of DNA-directed RNA polymerase |
| M896_031280 | 0.001388889 | 0.58769561 | Cg_03_78 | WD40 domain-containing protein |
| M896_031290 | 0.00169714 | Inf | Cg_03_79 | Spt16/Cdc68-like protein |
| M896_031300 | 0.001843158 | 0.40128792 | Cg_03_80 | hypothetical protein |
| M896_031310 | 0.000880301 | 0.88459888 | Cg_03_81 | hypothetical protein |
| M896_031320 | 0.00119431 | 2.36063346 | Cg_03_82 | ABCG-like transporter |
| M896_031330 | 0.006530214 | 0.48815094 | Cg_03_83 | hypothetical protein |
| M896_031340 | 0.000854701 | 0 | Cg_03_84 | hypothetical protein |
| M896_031350 | 0.001456157 | 0 | Cg_03_85 | hypothetical protein |
| M896_031360 | 0.004906205 | 1.41571061 | Cg_03_86 | U6 snRNA associated Sm-like protein |
| M896_031380 | 0 | Inf | Cg_03_87 | hypothetical protein |
| M896_031390 | 0.005206656 | 1.69680902 | Cg_03_88 | phosphatidylinositol-4-phosphate 5-kinase |
| M896_031410 | 0.001667893 | 0.74696545 | Cg_03_89 | ribosomal protein L13 |
| M896_031420 | 0.000486618 | 0 | Cg_03_90 | ribosomal protein S16 |
| M896_031430 | 0.002043423 | Inf | Cg_03_91 | subunit F of V-type ATPase |
| M896_031440 | 0.004921005 | 0.72120383 | Cg_03_92 | SCF ubiquitin ligase Skp1-like protein |
| M896_031460 | 0.001510721 | 2.42826583 | Cg_03_94 | UTP-glucose-1-phosphate uridylyltransferase |
| M896_031470 | 0.002149518 | 5.5592077 | Cg_03_95 | subunit gamma of vesicle coat complex |
| M896_031480 | 0 | Inf | Cg_03_96 | hypothetical protein |
| M896_040020 | 0.001807643 | 0.71883382 | Cg_05F_3 | PCI domain-containing protein |
| M896_040030 | 0.005754183 | 0.33839177 | Cg_04_212 | hypothetical protein |
| M896_040040 | 0.001423761 | 0.49895941 | Cg_04_211 | hypothetical protein |
| M896_040050 | 0.002361615 | Inf | Cg_04_210 | translation initiation factor IF-2 |
| M896_040060 | 0.001446759 | Inf | Cg_04_209 | hypothetical protein |
| M896_040070 | 0.0013225 | Inf | Cg_04_208 | hypothetical protein |
| M896_040080 | 0.005158449 | 0.53226758 | Cg_04_207 | hypothetical protein |
| M896_040090 | 0.005544467 | 0.39772727 | Cg_04_206 | hypothetical protein |
| M896_040100 | 0.003005904 | 0.10359702 | Cg_04_205 | hypothetical protein |
| M896_040110 | 0.003654756 | Inf | Cg_04_204 | ER translocation protein Sec61 |
| M896_040120 | 0.003222703 | Inf | Cg_04_203 | O-sialoglycoprotein endopeptidase |
| M896_040130 | 0.00351311 | 2.99419921 | Cg_04_202 | HrpA-like helicase |
| M896_040140 | 0.002764084 | 2.96730311 | Cg_04_201 | hypothetical protein |
| M896_040150 | 0.002401302 | Inf | Cg_04_200 | Rab GTPase |
| M896_040160 | 0.003241754 | Inf | Cg_04_199 | hypothetical protein |
| M896_040170 | 0.003004535 | 0.43953451 | Cg_04_198 | hypothetical protein |
| M896_040180 | 0.001505249 | 0.89562584 | Cg_04_197 | hypothetical protein |
| M896_040190 | 0.000925926 | 0.38816456 | Cg_04_196 | hypothetical protein |
| M896_040200 | 0.001931114 | Inf | Cg_04_195 | hypothetical protein |
| M896_040210 | 0 | Inf | Cg_04_194 | hypothetical protein |
| M896_040220 | 0.000991543 | Inf | Cg_04_193 | hypothetical protein |
| M896_040230 | 0.002147947 | 0.99031166 | Cg_04_192 | hypothetical protein |
| M896_040240 | 0.000855503 | 0.90214339 | Cg_04_191 | hypothetical protein |
| M896_040250 | 0.00262674 | 1.28482099 | Cg_04_190 | TATA binding protein associated factor 4-like protein |
| M896_040260 | 0.002548785 | 0.24664879 | Cg_04_189 | hypothetical protein |
| M896_040270 | 0.003047351 | 0.22691766 | Cg_04_188 | hypothetical protein |
| M896_040280 | 0.001674969 | 0.3179821 | Cg_04_187 | Ca2+-binding actin-bundling fimbrin/plastin |
| M896_040290 | 0.003242148 | 1.26786759 | Cg_04_186 | hypothetical protein |
| M896_040300 | 0.001525054 | 3.94695854 | Cg_04_185 | telomerase reverse transcriptase |
| M896_040310 | 0.001856738 | 0.8822441 | Cg_04_184 | hypothetical protein |
| M896_040320 | 0.001374225 | Inf | Cg_04_183 | subunit alpha of proteasome |
| M896_040330 | 0.00132716 | Inf | Cg_04_182 | DWNN ubiquitin-like domain-containing protein |
| M896_040340 | 0.003854694 | 2.00255845 | Cg_04_181 | pre-ribosome nuclear export protein |
| M896_040350 | 0.001738473 | 0.27702905 | Cg_04_180 | hypothetical protein |
| M896_040360 | 0.003006967 | Inf | Cg_04_179 | hypothetical protein |
| M896_040370 | 0.002973978 | 0.39107418 | Cg_04_178 | hypothetical protein |
| M896_040390 | 0.002188552 | 0.28644435 | Cg_04_177 | subunit alpha of mRNA capping enzyme |
| M896_040410 | 0.002578959 | 2.3392881 | Cg_04_176 | superfamily II DNA/RNA helicase |
| M896_040420 | 0.002200527 | Inf | Cg_04_175 | hypothetical protein |
| M896_040430 | 0.008112875 | 0.11226872 | Cg_04_174 | hypothetical protein |
| M896_040440 | 0.002505202 | 1.39214713 | Cg_04_173 | type-B delta catalytic subunit of DNA polymerase |
| M896_040450 | 0.001851852 | Inf | Cg_04_172 | histone H4 |
| M896_040460 | 0.007626886 | 1.10966844 | Cg_04_171 | histone H3 |
| M896_040470 | 0.000940623 | Inf | Cg_04_170 | hypothetical protein |
| M896_040480 | 0 | Inf | Cg_04_168 | hypothetical protein |
| M896_040490 | 0.001423886 | Inf | Cg_04_167 | subunit beta of T complex protein 1 |
| M896_040500 | 0.002386831 | 1.05284068 | Cg_04_166 | hypothetical protein |
| M896_040510 | 0.001362567 | 0.0461944 | Cg_04_165 | hypothetical protein |
| M896_040520 | 0.001959379 | 0.76876877 | Cg_04_164 | hypothetical protein |
| M896_040530 | 0.001297879 | Inf | Cg_04_163 | hypothetical protein |
| M896_040540 | 0.001921042 | Inf | Cg_04_162 | hypothetical protein |
| M896_040550 | 0.004204204 | Inf | Cg_04_160 | ribosomal protein S29 |
| M896_040560 | 0 | Inf | Cg_04_159 | hypothetical protein |
| M896_040570 | 0 | Inf | Cg_04_158 | ribosomal protein S5 |
| M896_040580 | 0.002659069 | 0.265625 | Cg_04_157 | hypothetical protein |
| M896_040590 | 0.001916602 | 0.31062626 | Cg_04_155 | hypothetical protein |
| M896_040610 | 0.003116385 | 3.42480298 | Cg_04_154 | hypothetical protein |
| M896_040620 | 0.000819252 | Inf | Cg_04_153 | transcription elongation factor S-II |
| M896_040630 | 0.000927472 | Inf | Cg_04_152 | hypothetical protein |
| M896_040650 | 0.003744856 | 0.34888602 | Cg_04_151 | protein phosphatase 2A |
| M896_040660 | 0.001186624 | 0.40462376 | Cg_04_150 | putative glucose transporter |
| M896_040670 | 0.001545103 | 0.41454023 | Cg_04_149 | SAGA/ADA complex histone acetyltransferase |
| M896_040680 | 0.003560149 | 1.95512985 | Cg_04_148 | hypothetical protein |
| M896_040690 | 0.002147469 | 0.5280638 | Cg_04_147 | HrpA-like helicase |
| M896_040700 | 0.001634738 | 0.85954927 | Cg_04_146 | DHHC-type Zn finger domain-containing protein |
| M896_040710 | 0.002134338 | Inf | Cg_04_145 | hypothetical protein |
| M896_040720 | 0.002222222 | Inf | Cg_04_144 | RNA polymerase II transcription factor |
| M896_040740 | 0.00140802 | Inf | Cg_04_143 | protein tyrosine phosphatase |
| M896_040750 | 0.00141759 | 1.46713313 | Cg_04_142 | 26S proteasome regulatory complex protein |
| M896_040760 | 0.001378122 | Inf | Cg_04_141 | hypothetical protein |
| M896_040770 | 0.00667334 | 0.83003195 | Cg_04_140 | ribosomal protein L27 |
| M896_040780 | 0.00359147 | 0.28275266 | Cg_04_139 | hypothetical protein |
| M896_040790 | 0.002731567 | 1.86831108 | Cg_04_138 | DNA topoisomerase II |
| M896_040800 | 0.000416667 | Inf | Cg_04_137 | hypothetical protein |
| M896_040810 | 0.001453224 | 0.1277081 | Cg_04_136 | hypothetical protein |
| M896_040820 | 0.002334267 | Inf | Cg_04_135 | hypothetical protein |
| M896_040840 | 0.001267057 | Inf | Cg_04_130 | heat shock transcription factor |
| M896_040850 | 0.00301619 | 0.56260042 | Cg_04_129 | hypothetical protein |
| M896_040860 | 0.000888889 | Inf | Cg_04_128 | hypothetical protein |
| M896_040870 | 0.002704779 | Inf | Cg_04_126 | non-ATPase subunit 7 of 26S proteasome |
| M896_040880 | 0.001952862 | 0.61713228 | Cg_04_125 | hypothetical protein |
| M896_040890 | 0.000333667 | Inf | Cg_04_124 | hypothetical protein |
| M896_040900 | 0.000815794 | Inf | Cg_04_123 | ribosomal protein SA |
| M896_040920 | 0.002432691 | 0.68861602 | Cg_04_122 | dimethyladenosine transferase |
| M896_040930 | 0.003731343 | 1.16044757 | Cg_04_121 | hypothetical protein |
| M896_040940 | 0.000457081 | 0.08975059 | Cg_04_120 | MdlB-type ABC transporter |
| M896_040960 | 0.001405358 | 0.47366091 | Cg_04_119 | E3 ubiquitin-protein ligase |
| M896_040970 | 0.00342241 | 0.51423614 | Cg_04_118 | hypothetical protein |
| M896_040980 | 0.000663746 | 0.13662266 | Cg_04_117 | putative RNA-binding protein |
| M896_040990 | 0.0015273 | 1.33975388 | Cg_04_116 | hypothetical protein |
| M896_041010 | 0.000771605 | Inf | Cg_04_115 | uridine diphosphate-N-acetylglucosamine |
| M896_041020 | 0 | Inf | Cg_04_114 | hypothetical protein |
| M896_041030 | 0.002085994 | Inf | Cg_04_113 | hypothetical protein |
| M896_041040 | 0.002496329 | 0.73675944 | Cg_04_112 | lysyl-tRNA synthetase |
| M896_041050 | 0.003024575 | 0.63961461 | Cg_04_111 | X-prolyl aminopeptidase |
| M896_041060 | 0.003735632 | 0.42403249 | Cg_04_110 | hypothetical protein |
| M896_041070 | 0.001576504 | Inf | Cg_04_109 | putative methyltransferase |
| M896_041080 | 0.001910086 | Inf | Cg_04_108 | ubiquitin-conjugating enzyme E2 |
| M896_041090 | 0.00237037 | 0.23853211 | Cg_04_107 | ribosomal protein S11 |
| M896_041100 | 0.003342693 | 1.02291845 | Cg_04_106 | putative lipase |
| M896_041110 | 0.001092459 | 0.15657745 | Cg_04_105 | putative dolichol kinase |
| M896_041120 | 0.002868555 | Inf | Cg_04_104 | Rab11-like GTPase |
| M896_041130 | 0.002149871 | 0.46836677 | Cg_04_103 | vacuolar protein sorting-associated protein |
| M896_041140 | 0.002007869 | 0.14252772 | Cg_04_102 | PP2A-like phosphoprotein phosphatase |
| M896_041150 | 0.002080378 | 0.42932347 | Cg_04_101 | mitotic phase inducer phosphatase |
| M896_041160 | 0.001897019 | Inf | Cg_04_100 | peptidyl-tRNA hydrolase |
| M896_041170 | 0.002429943 | 0.51053806 | Cg_04_99 | putative 6-phosphogluconolactonase |
| M896_041180 | 0.007149758 | 0.76205358 | Cg_04_98 | ribosomal protein L22 |
| M896_041190 | 0.003816993 | 0.35022845 | Cg_04_97 | seryl-tRNA synthetase |
| M896_041200 | 0.002031122 | 0.58634839 | Cg_04_96 | subunit of U5 snRNP spliceosome |
| M896_041210 | 0.000495893 | Inf | Cg_04_95 | hypothetical protein |
| M896_041220 | 0.001399556 | Inf | Cg_04_94 | translation initiation factor 6 |
| M896_041230 | 0.001559454 | Inf | Cg_04_93 | Sm-like ribonucleoprotein |
| M896_041240 | 0.002678385 | 0.13758957 | Cg_04_92 | hypothetical protein |
| M896_041260 | 0.004575163 | 0.92915811 | Cg_04_89 | Nop56p-like protein |
| M896_041270 | 0.004063986 | Inf | Cg_04_88 | hypothetical protein |
| M896_041280 | 0.002012882 | 1.09930716 | Cg_04_87 | asparagine synthase |
| M896_041290 | 0.00151005 | 1.31535173 | Cg_04_86 | minichromosome maintenance protein |
| M896_041300 | 0.005291005 | 0.79615967 | Cg_04_85 | hypothetical protein |
| M896_041310 | 0.002512179 | 0.44256414 | Cg_04_84 | cohesin |
| M896_041320 | 0.003006967 | Inf | Cg_04_83 | hypothetical protein |
| M896_041330 | 0.002155716 | Inf | Cg_04_82 | pseudouridine synthase |
| M896_041340 | 0.003653942 | 0.51168871 | Cg_04_81 | subunit beta of phenylalanyl-tRNA synthetase |
| M896_041350 | 0.002024804 | 0.66373703 | Cg_04_79 | superfamily II RNA helicase |
| M896_041370 | 0.007380174 | 2.41681575 | Cg_04_78 | hypothetical protein |
| M896_041380 | 0.002446103 | 0.9068609 | Cg_04_77 | chromosome segregation ATPase |
| M896_041390 | 0.000984529 | Inf | Cg_04_76 | subunit of endoplasmic reticulum translocation complex |
| M896_041400 | 0.001742919 | Inf | Cg_04_75 | subunit of transcription initiation factor TFIID |
| M896_041410 | 0.004840484 | Inf | Cg_04_74 | hypothetical protein |
| M896_041420 | 0.006887589 | 0.96466067 | Cg_04_73 | homeodomain-containing protein |
| M896_041430 | 0.001804589 | 2.84915994 | Cg_04_72 | signal recognition particle GTPase |
| M896_041440 | 0.000522108 | 0 | Cg_04_71 | hypothetical protein |
| M896_041450 | 0.002253997 | 1.6988106 | Cg_04_70 | myosin heavy chain |
| M896_041460 | 0 | Inf | Cg_04_69 | ribosomal protein L29e |
| M896_041470 | 0.001458151 | Inf | Cg_04_68 | hypothetical protein |
| M896_041480 | 0.002469136 | Inf | Cg_04_67 | ribosomal protein S27 |
| M896_041490 | 0.001021115 | Inf | Cg_04_66 | subunit theta of T-complex protein 1 |
| M896_041500 | 0.000560224 | Inf | Cg_04_65 | homeodomain-containing protein |
| M896_041510 | 0.004208099 | Inf | Cg_04_64 | hypothetical protein |
| M896_041520 | 0.004206676 | 0.24946439 | Cg_04_63 | hypothetical protein |
| M896_041530 | 0.003128371 | Inf | Cg_04_62 | dolichol-phosphate mannosyltransferase |
| M896_041540 | 0.001649732 | 0.55547804 | Cg_04_61 | topoisomerase IA |
| M896_041550 | 0.000486618 | Inf | Cg_04_59 | hypothetical protein |
| M896_041560 | 0.005201966 | 0.39446158 | Cg_04_58 | actin-like protein |
| M896_041570 | 0.001879729 | 0.86292611 | Cg_04_56 | translation elongation factor EF-1 alpha |
| M896_041580 | 0.002161506 | 0.17543388 | Cg_04_55 | subunit A of DNA topoisomerase VI |
| M896_041590 | 0.004670913 | 0.77369041 | Cg_04_54 | hypothetical protein |
| M896_041600 | 0.002418226 | 0.67363933 | Cg_04_53 | alpha-1%2C2 mannosyl-transferase |
| M896_041610 | 0.002539182 | 0.66975432 | Cg_04_52 | valyl-tRNA synthetase |
| M896_041620 | 0.002039721 | 0.19669692 | Cg_04_51 | hypothetical protein |
| M896_041630 | 0.004541591 | 0.29956319 | Cg_04_50 | subunit beta of pyruvate dehydrogenase |
| M896_041640 | 0 | Inf | Cg_04_49 | translation initiation factor IF1A |
| M896_041650 | 0.003414352 | 0.49215792 | Cg_04_48 | amino acid permease |
| M896_041660 | 0.001863499 | 0.11052669 | Cg_04_47 | SWIB domain-containing protein |
| M896_041670 | 0.003128594 | 0.27003078 | Cg_04_46 | ubiquitin fusion degradation protein 2 |
| M896_041680 | 0.001284958 | Inf | Cg_04_45 | thymidylate kinase |
| M896_041690 | 0.00302204 | 0.25655053 | Cg_04_44 | hypothetical protein |
| M896_041700 | 0.001311359 | Inf | Cg_04_43 | hypothetical protein |
| M896_041710 | 0.001079797 | Inf | Cg_04_42 | WD40 domain-containing protein |
| M896_041720 | 0.003420875 | 2.74933679 | Cg_04_41 | nuclear protein export factor |
| M896_041730 | 0.002432825 | Inf | Cg_04_40 | hypothetical protein |
| M896_041740 | 0.001295592 | Inf | Cg_04_39 | lysophospholipid acyltransferase |
| M896_041750 | 0.001531295 | 2.33281493 | Cg_04_38 | superfamily II RNA helicase |
| M896_041760 | 0.000883691 | 0.62480102 | Cg_04_37 | putative RNA-binding protein |
| M896_041770 | 0.001670844 | Inf | Cg_04_36 | ribosomal protein L32 |
| M896_041780 | 0.001044216 | Inf | Cg_04_35 | hypothetical protein |
| M896_041790 | 0.003333333 | 0.11269959 | Cg_04_33 | hypothetical protein |
| M896_041800 | 0.000979968 | 0.09126816 | Cg_04_32 | hypothetical protein |
| M896_041810 | 0.002276867 | 0.12599772 | Cg_04_31 | hypothetical protein |
| M896_041820 | 0.002035002 | 0 | Cg_04_30 | ribosomal protein S10 |
| M896_041840 | 0.001593999 | 0.23743637 | Cg_04_29 | hypothetical protein |
| M896_041850 | 0.00308642 | Inf | Cg_04_28 | ribosomal protein L13a |
| M896_041860 | 0.001970055 | Inf | Cg_04_27 | hypothetical protein |
| M896_041870 | 0.00174526 | 1.08449703 | Cg_04_26 | subunit beta' of DNA-directed RNA polymerase |
| M896_041890 | 0.002682895 | 1.30147277 | Cg_04_25 | hypothetical protein |
| M896_041910 | 0.002934564 | Inf | Cg_04_24 | hypothetical protein |
| M896_041920 | 0.005372405 | 0.7063997 | Cg_04_23 | hypothetical protein |
| M896_041930 | 0.003106332 | 3.45880462 | Cg_04_22 | hypothetical protein |
| M896_041940 | 0.002939447 | 0 | Cg_04_21 | hypothetical protein |
| M896_041950 | 0.00198059 | Inf | Cg_04_20 | TATA-box binding protein |
| M896_041960 | 0.001446759 | Inf | Cg_04_19 | hypothetical protein |
| M896_041970 | 0.001486126 | Inf | Cg_04_18 | hypothetical protein |
| M896_041980 | 0.001063727 | Inf | Cg_04_17 | Na+/H+ antiporter |
| M896_041990 | 0.002660913 | 0.25626779 | Cg_04_16 | Zn-finger protein |
| M896_042000 | 0.001832917 | 0.76631942 | Cg_04_15 | hypothetical protein |
| M896_042010 | 0.001128748 | 0.3680418 | Cg_04_14 | regulator of chromosome condensation |
| M896_042020 | 0.000251572 | 0 | Cg_04_13 | tRNA-binding domain-containing CsaA-like protein |
| M896_042030 | 0.001171257 | Inf | Cg_04_12 | hypothetical protein |
| M896_042040 | 0.002762084 | 0.27719767 | Cg_04_11 | GTPase activating protein |
| M896_042050 | 0.001514221 | 0.19164691 | Cg_04_10 | hypothetical protein |
| M896_042060 | 0.004901961 | 1.41350803 | Cg_04_9 | subunit beta of mRNA capping enzyme |
| M896_042070 | 0.003537866 | Inf | Cg_04_8 | GTP-binding nuclear protein |
| M896_042080 | 0.000555556 | Inf | Cg_04_7 | hypothetical protein |
| M896_042090 | 0.000424798 | 0 | Cg_04_6 | hypothetical protein |
| M896_042100 | 0.001011941 | Inf | Cg_04_5 | hypothetical protein |
| M896_042110 | 0 | Inf | Cg_04_4 | hypothetical protein |
| M896_042120 | 0.001572723 | 1.13738237 | Cg_04_3 | hypothetical protein |
| M896_042130 | 0.001814059 | 0.1280262 | Cg_04_2 | hypothetical protein |
| M896_042140 | 0 | Inf | Cg_04_1 | hypothetical protein |
| M896_050040 | 0 | Inf | Cg_05A_60 | subunit D of vacuolar-type H+-ATPase |
| M896_050050 | 0.000627746 | Inf | Cg_05A_59 | hypothetical protein |
| M896_050080 | 0.002239906 | 1.02774247 | Cg_05A_56 | signal recognition particle GTPase |
| M896_050090 | 0.003808328 | Inf | Cg_05A_55 | ribose 5-phosphate isomerase |
| M896_050100 | 0.003568202 | Inf | Cg_05A_54 | subunit Rpn7 of 26S proteasome regulatory complex |
| M896_050120 | 0.002968961 | 0.82949551 | Cg_05A_53 | hypothetical protein |
| M896_050130 | 0.002247547 | 0.11092437 | Cg_05A_52 | subunit C34 of DNA-directed RNA polymerase III |
| M896_050140 | 0.000361337 | Inf | Cg_05A_51 | hypothetical protein |
| M896_050150 | 0.002743484 | 0 | Cg_05A_50 | hypothetical protein |
| M896_050160 | 0.002018822 | Inf | Cg_05A_49 | subunit gamma of T-complex protein 1 |
| M896_050180 | 0 | Inf | Cg_05A_47 | histone H3/H4-like protein |
| M896_050190 | 0.002524312 | 0.36207778 | Cg_05A_46 | RNase PH-like protein |
| M896_050200 | 0.000896735 | 0.69095255 | Cg_05A_45 | putative P-loop ATPase/acetyltransferase fusion protein |
| M896_050210 | 0.002760943 | 0.43682144 | Cg_05A_44 | U1 snRNP-specific protein C |
| M896_050230 | 0.001099537 | Inf | Cg_05A_43 | acyl-CoA cholesterol acyltransferase |
| M896_050240 | 0.002159704 | Inf | Cg_05A_42 | 1-acyl-SN-glycerol-3-phosphate acyltransferase |
| M896_050250 | 0.001571268 | 1.26338924 | Cg_05A_41 | kinesin-like protein |
| M896_050260 | 0.001185185 | 0.2345679 | Cg_05A_40 | hypothetical protein |
| M896_050270 | 0.000768403 | 0 | Cg_05A_39 | inorganic pyrophosphatase |
| M896_050280 | 0.001427469 | Inf | Cg_05A_37 | Rho GTPase |
| M896_050290 | 0.001944444 | 0 | Cg_05A_36 | hypothetical protein |
| M896_050300 | 0 | Inf | Cg_05A_35 | putative actin depolymerization factor |
| M896_050310 | 0.002867384 | Inf | Cg_05A_34 | hypothetical protein |
| M896_050320 | 0.002145002 | 0.50040225 | Cg_05A_33 | mRNA capping enzyme |
| M896_050330 | 0.000973007 | 0.38836256 | Cg_05A_32 | hypothetical protein |
| M896_050340 | 0.006688778 | 1.30876875 | Cg_05A_31 | ribosomal protein S23 |
| M896_050350 | 0.000620516 | Inf | Cg_05A_30 | regulatory complex subunit of 26S proteasome |
| M896_050360 | 0.002225209 | Inf | Cg_05A_29 | ATP/ADP translocase |
| M896_050370 | 0.001224847 | Inf | Cg_05A_28 | hypothetical protein |
| M896_050380 | 0.002279202 | 0.90564593 | Cg_05A_27 | ATP-dependent rRNA helicase |
| M896_050390 | 0.003697125 | 2.64838989 | Cg_05A_26 | hypothetical protein |
| M896_050410 | 0.003652263 | Inf | Cg_05A_25 | fragile histidine family hydrolase |
| M896_050420 | 0.001172928 | Inf | Cg_05A_24 | hypothetical protein |
| M896_050430 | 0.00285748 | 0.93231785 | Cg_05A_23 | hypothetical protein |
| M896_050440 | 0.003355371 | 0.43757375 | Cg_05A_22 | ATP/ADP translocase |
| M896_050450 | 0.002603969 | 0.38895317 | Cg_05A_21 | uracil-DNA glycosylase |
| M896_050460 | 0.001851852 | 1.05547445 | Cg_05A_20 | ATP/ADP translocase |
| M896_050470 | 0.000881834 | 0 | Cg_05A_19 | subunit alpha of 20S proteasome |
| M896_050480 | 0.003244838 | 1.37634137 | Cg_05A_18 | putative short chain dehydrogenase |
| M896_050490 | 0.000750117 | Inf | Cg_05A_17 | AGC serine/threonine kinase |
| M896_050500 | 0.00333855 | 0.27896841 | Cg_05A_16 | putative TFIIF-interacting CTD phosphatase |
| M896_050510 | 0.003120796 | 0.5671125 | Cg_05A_15 | phosphoinositide 3-kinase |
| M896_050520 | 0.000864895 | 1.51794341 | Cg_05A_13 | replication factor A protein 1 |
| M896_050530 | 0 | Inf | Cg_05A_12 | hypothetical protein |
| M896_050540 | 0.000887166 | 0.46113687 | Cg_05A_11 | importin |
| M896_050550 | 0.00281294 | 1.48291408 | Cg_05A_10 | subunit eta of T-complex protein 1 |
| M896_050570 | 0.003552251 | 1.03536676 | Cg_05A_7 | hypothetical protein |
| M896_050580 | 0.000532141 | Inf | Cg_05A_6 | MYST histone acetyltransferase |
| M896_050590 | 0 | Inf | Cg_05A_5 | ribosomal biogenesis protein |
| M896_050600 | 0.000874349 | 0.62142295 | Cg_05A_4 | HrpA-like helicase |
| M896_050610 | 0.003292181 | 0.34450356 | Cg_05A_3 | ATP-dependent RNA helicase |
| M896_050620 | 0.00045819 | Inf | Cg_05A_2 | subunit of translation initiation factor 2B |
| M896_050630 | 0.001965056 | 0.48955817 | Cg_05D_1 | MutS-like mismatch repair ATPase |
| M896_050640 | 0.003125 | 0.78765761 | Cg_05D_2 | hypothetical protein |
| M896_050650 | 0.00328673 | 0.5132996 | Cg_05D_3 | hypothetical protein |
| M896_050660 | 0.002862986 | 1.27862187 | Cg_05D_4 | methionine aminopeptidase 2 |
| M896_050670 | 0.003890924 | 1.15798005 | Cg_05D_5 | hypothetical protein |
| M896_050680 | 0.002991823 | 1.42208001 | Cg_05D_6 | subunit of transcription initiation factor TFIID |
| M896_050690 | 0.002373247 | 0 | Cg_05D_7 | hypothetical protein |
| M896_050700 | 0.002030344 | 0.67762628 | Cg_05D_8 | small subunit of replication factor C |
| M896_050710 | 0.002540705 | 0.63141725 | Cg_05D_9 | hypothetical protein |
| M896_050720 | 0 | Inf | Cg_05D_10 | putative methyltransferase |
| M896_050740 | 0.001851852 | 2.9941245 | Cg_05D_12 | chromosome segregation protein |
| M896_050750 | 0.002851153 | Inf | Cg_05D_13 | fibrillarin |
| M896_050760 | 0.00109979 | 0.08508698 | Cg_05D_14 | HMG-like nuclear protein |
| M896_050770 | 0.002310858 | 0.50527682 | Cg_05D_15 | subunit of U1 snRNP |
| M896_050780 | 0.001415146 | 0.27428768 | Cg_05D_16 | hypothetical protein |
| M896_050790 | 0.002362708 | Inf | Cg_05D_17 | hypothetical protein |
| M896_050800 | 0.00216644 | 0.58934642 | Cg_05D_18 | glycerol-3-phosphate dehydrogenase |
| M896_050810 | 0.001595026 | Inf | Cg_05D_19 | hypothetical protein |
| M896_050820 | 0.002350932 | 1.69732892 | Cg_05D_20 | putative RNA-processing beta-lactamase |
| M896_050830 | 0.00191752 | Inf | Cg_05D_21 | long-chain acyl-CoA synthetase |
| M896_050840 | 0.000883838 | Inf | Cg_05D_22 | ribonucleoside-diphosphate reductase |
| M896_050850 | 0.000441501 | Inf | Cg_05D_23 | ubiquitin-conjugating enzyme |
| M896_050860 | 0.002555772 | Inf | Cg_05D_24 | hypothetical protein |
| M896_050870 | 0.006814815 | Inf | Cg_05D_25 | putative kinase |
| M896_050880 | 0.002116402 | 1.03490816 | Cg_05D_26 | hypothetical protein |
| M896_050890 | 0.000243309 | Inf | Cg_05D_27 | hypothetical protein |
| M896_050900 | 0.001564945 | Inf | Cg_05D_28 | ribosomal protein L27 |
| M896_050910 | 0.003713176 | 1.99819147 | Cg_05D_29 | kinesin-like protein |
| M896_050920 | 0.004002389 | Inf | Cg_05D_30 | hypothetical protein |
| M896_050930 | 0.001880053 | 0.53012685 | Cg_05D_31 | hypothetical protein |
| M896_050940 | 0.002558666 | 0.81698184 | Cg_05D_32 | hypothetical protein |
| M896_050960 | 0.000580304 | Inf | Cg_05D_33 | subunit B of V-type ATP synthase |
| M896_050970 | 0.010185185 | 0.35235096 | Cg_05D_34 | DNA directed RNA polymerase |
| M896_050980 | 0.002743484 | Inf | Cg_05D_35 | hypothetical protein |
| M896_050990 | 0.001392111 | 1.88469134 | Cg_05D_36 | phosphoglyceromutase |
| M896_051000 | 0.003703704 | Inf | Cg_05D_37 | hypothetical protein |
| M896_051010 | 0.003632819 | 0.83979445 | Cg_05D_38 | subunit B of DNA polymerase alpha |
| M896_051020 | 0.005770411 | 1.06691919 | Cg_05D_39 | chromatin remodeling protein |
| M896_051030 | 0.003600823 | 0.27890121 | Cg_05D_40 | DnaJ-like protein |
| M896_051040 | 0.001117818 | 0.12805392 | Cg_05D_41 | polyadenylate-binding protein |
| M896_051060 | 0.000549511 | Inf | Cg_05D_42 | subunit of transcription initiation factor TFIID |
| M896_051070 | 0.001303449 | 0.16200789 | Cg_05D_43 | subunit 2 of origin recognition complex |
| M896_051080 | 0.002426694 | 1.17790364 | Cg_05D_44 | guanine nucleotide exchange factor |
| M896_051090 | 0.002710716 | 0.48039477 | Cg_05D_45 | hypothetical protein |
| M896_051100 | 0.002436647 | Inf | Cg_05D_46 | hypothetical protein |
| M896_051110 | 0.002258831 | 0.95240851 | Cg_05D_47 | hypothetical protein |
| M896_051120 | 0.002729045 | 0.50554899 | Cg_05D_48 | Rad14-like DNA excision repair protein |
| M896_051130 | 0.001268861 | Inf | Cg_05D_49 | ubiquitin fusion-degradation protein |
| M896_051140 | 0.002956803 | 1.77423768 | Cg_05D_50 | hypothetical protein |
| M896_051150 | 0.001595442 | Inf | Cg_05D_51 | subunit beta of casein kinase II |
| M896_051160 | 0.00193613 | 1.03147387 | Cg_05D_52 | DNA polymerase epsilon |
| M896_051170 | 0 | Inf | Cg_05D_53 | hypothetical protein |
| M896_051180 | 0.003748446 | 3.53379156 | Cg_05D_54 | importin beta |
| M896_051190 | 0.002095113 | 0.28421181 | Cg_05D_55 | hypothetical protein |
| M896_051200 | 0.001152263 | Inf | Cg_05D_56 | lysophospholipid acyltransferase |
| M896_051210 | 0.001239057 | 0.52697981 | Cg_05D_57 | PP2A-like protein phosphatase |
| M896_051220 | 0.002300437 | Inf | Cg_05D_58 | hypothetical protein |
| M896_051230 | 0 | Inf | Cg_05D_59 | ribosomal protein L44 |
| M896_051240 | 0.001338688 | Inf | Cg_05D_60 | ubiquitin-conjugating enzyme E2 |
| M896_051250 | 0.00137741 | 1.19970799 | Cg_05D_61 | SNF2-like helicase |
| M896_051330 | 0.002935863 | Inf | Cg_05D_62 | nucleotide excision repair endonuclease Nef1 |
| M896_051340 | 0.001817558 | 0.51592582 | Cg_05D_63 | Rrp4-like RNA-binding protein |
| M896_051350 | 0.002727414 | 1.65511061 | Cg_05D_64 | putative RNA-processing beta-lactamase |
| M896_051360 | 0.001362963 | Inf | Cg_05D_65 | hypothetical protein |
| M896_051370 | 0.004280328 | 1.08269922 | Cg_05D_66 | phosphatidylinositol 5-phosphate phosphatase |
| M896_051380 | 0.00194298 | 0.89140133 | Cg_05D_67 | ubiquitin-protein ligase |
| M896_051390 | 0.000560917 | Inf | Cg_05D_68 | hypothetical protein |
| M896_051400 | 0.001054131 | 0.93842802 | Cg_05D_69 | hypothetical protein |
| M896_051410 | 0.0022087 | 0.27145504 | Cg_05D_70 | hypothetical protein |
| M896_051420 | 0.000363306 | Inf | Cg_05D_71 | regulatory subunit of ATP-dependent 26S proteasome |
| M896_051430 | 0.003003003 | 1.37624386 | Cg_05D_72 | chromatin remodeling transcription factor |
| M896_051440 | 0.001224105 | Inf | Cg_05D_73 | PAP2-like phosphatidic acid phosphatase |
| M896_051450 | 0 | Inf | Cg_05D_74 | beta type-6 subunit of proteasome |
| M896_051460 | 0.002469136 | 1.0318323 | Cg_05D_75 | glutaminyl-tRNA synthetase |
| M896_051470 | 0 | Inf | Cg_05D_76 | hypothetical protein |
| M896_051480 | 0.001977796 | 0.89979039 | Cg_05D_77 | insulinase-like Zn-dependent peptidase |
| M896_051490 | 0 | Inf | Cg_05D_78 | putative homeodomain-containing transcription factor |
| M896_051500 | 0.001076658 | Inf | Cg_05D_79 | hypothetical protein |
| M896_051510 | 0.000818382 | 0.77131783 | Cg_05D_80 | hypothetical protein |
| M896_051520 | 0.001615368 | 0.19317147 | Cg_05D_81 | hypothetical protein |
| M896_051550 | 0.00405855 | 0.73268293 | Cg_05E_2 | White-like ABC transporter |
| M896_051560 | 0 | Inf | Cg_05E_3 | hypothetical protein |
| M896_051570 | 0.002564103 | 0.41585815 | Cg_05E_4 | ubiquitin-protein ligase |
| M896_051580 | 0.001833517 | 0 | Cg_05E_5 | hypothetical protein |
| M896_051590 | 0.001413627 | 0 | Cg_05E_6 | ribosomal protein S24 |
| M896_051610 | 0.002973777 | 1.48231848 | Cg_05E_7 | subunit D2 of Sm-like ribonucleoprotein |
| M896_051620 | 0.001304121 | 0 | Cg_05E_8 | choline-phosphate cytidylyltransferase |
| M896_051630 | 0.001006441 | Inf | Cg_05E_9 | Mnd1-like meiotic recombination protein |
| M896_051640 | 0.003703704 | 1.80503145 | Cg_05E_10 | hypothetical protein |
| M896_051650 | 0.005175038 | Inf | Cg_05E_11 | hypothetical protein |
| M896_051660 | 0.001771801 | 0.95284924 | Cg_05E_12 | hypothetical protein |
| M896_051670 | 0.001108958 | 0.21013728 | Cg_05E_13 | hypothetical protein |
| M896_051680 | 0.002388889 | 0.26201754 | Cg_05E_15 | spore wall protein Swp1b |
| M896_051690 | 0.001831591 | 0.07420233 | Cg_05E_16 | ATP-dependent RNA helicase |
| M896_051700 | 0.002391789 | 0.16250365 | Cg_05E_17 | enolase |
| M896_051710 | 0.005787037 | 0.5467944 | Cg_05E_18 | hypothetical protein |
| M896_051720 | 0.001298929 | 0.29759001 | Cg_05E_19 | dynamin-like vacuolar protein-sorting protein |
| M896_051730 | 0.002394029 | 0.03013875 | Cg_05E_20 | hypothetical protein |
| M896_051740 | 0.000576132 | 0 | Cg_05E_21 | hypothetical protein |
| M896_051750 | 0.00333478 | 0.21413704 | Cg_05E_22 | cellular morphogenesis/cytokinesis regulation |
| M896_051760 | 0.001968906 | 0.147443 | Cg_05E_23 | Spc97/Spc98-like spindle pole body protein |
| M896_051770 | 0.000589971 | 0.26571251 | Cg_05E_24 | cyclophilin-type peptidylprolyl cis-trans |
| M896_051780 | 0.002575827 | 0.4405346 | Cg_05E_25 | leucyl aminopeptidase |
| M896_051790 | 0.003257158 | 0.21847203 | Cg_05E_26 | hypothetical protein |
| M896_051800 | 0.000794566 | 0.17005697 | Cg_05E_27 | glycyl-tRNA synthetase |
| M896_051810 | 0.001151298 | Inf | Cg_05E_28 | hypothetical protein |
| M896_051820 | 0.003106332 | 3.38372962 | Cg_05E_29 | hypothetical protein |
| M896_051830 | 0.002270012 | Inf | Cg_05E_30 | hypothetical protein |
| M896_051840 | 0.001448954 | 1.32253913 | Cg_05E_31 | elongation factor 3 |
| M896_051850 | 0.001776163 | 0.1968006 | Cg_05E_32 | putative DNA primase |
| M896_051860 | 0.003578458 | 0.24365821 | Cg_05E_33 | putative subunit of t-SNARE complex |
| M896_051870 | 0 | Inf | Cg_05E_34 | putative G10 protein |
| M896_051880 | 0.002338494 | 0.17138373 | Cg_05E_35 | putative gamma-glutamyltransferase |
| M896_051890 | 0.001624431 | 0.19269358 | Cg_05E_36 | hypothetical protein |
| M896_051900 | 0.002819092 | Inf | Cg_05E_37 | putative ATP-dependent RNA helicase |
| M896_051910 | 0.002228369 | 0.28257284 | Cg_05E_39 | putative subunit of Mre11 |
| M896_051930 | 0.002379451 | 0.52286545 | Cg_05F_27 | hypothetical protein |
| M896_051940 | 0 | Inf | Cg_05F_26 | LSM domain-containing protein |
| M896_051950 | 0.001490248 | Inf | Cg_05F_25 | ribosomal protein S4-like protein |
| M896_051960 | 0.004923999 | Inf | Cg_05F_24 | putative DNA replication protein kinase |
| M896_051970 | 0.003179145 | 0.26173285 | Cg_05F_23 | alpha subunit of proteasome |
| M896_051980 | 0 | Inf | Cg_05F_22 | PUA domain-containing protein |
| M896_052000 | 0.003379883 | 0.34588929 | Cg_05F_21 | putative nucleotidyl transferase |
| M896_052010 | 0.000912242 | 0.26786954 | Cg_05F_20 | putative Zn-dependent protease |
| M896_052020 | 0.003609701 | 0.69529453 | Cg_05F_19 | glutaredoxin domain-containing protein |
| M896_052030 | 0.002291896 | 0.13797437 | Cg_05F_18 | peptidase M48 domain-containing protein |
| M896_052040 | 0.00146871 | 0 | Cg_05F_17 | putative alpha subunit of proteasome |
| M896_052050 | 0.003398118 | 0.26258776 | Cg_05F_16 | hypothetical protein |
| M896_052060 | 0.001061498 | 0 | Cg_05F_15 | hypothetical protein |
| M896_052070 | 0.001861547 | 0.13469207 | Cg_05F_14 | hypothetical protein |
| M896_052080 | 0.000831444 | Inf | Cg_05F_13 | hypothetical protein |
| M896_052100 | 0.000718629 | 0.31745836 | Cg_05F_12 | hypothetical protein |
| M896_052110 | 0.000519818 | 0 | Cg_05F_11 | hypothetical protein |
| M896_052120 | 0 | Inf | Cg_05F_10 | hypothetical protein |
| M896_052130 | 0 | Inf | Cg_05F_9 | ribosomal protein L30E |
| M896_052140 | 0 | Inf | Cg_05F_8 | hypothetical protein |
| M896_052150 | 0.000269974 | Inf | Cg_05F_7 | hypothetical protein |
| M896_052160 | 0.001527778 | 0.39621314 | Cg_05F_6 | protein kinase domain-containing protein |
| M896_052170 | 0.001819813 | 0.14356279 | Cg_05F_5 | acyl-CoA thioesterase domain-containing protein |
| M896_052180 | 0.001711459 | 0.42695421 | Cg_05F_4 | DNA replication factor RFC1 |
| M896_060010 | 0.000569801 | Inf | Cg_07_67 | putative zinc finger protein |
| M896_060020 | 0.001386182 | Inf | Cg_07_66 | hypothetical protein |
| M896_060050 | 0.010344288 | 0.55279732 | Cg_06A_69 | hypothetical protein |
| M896_060060 | 0.001870813 | Inf | Cg_06A_68 | hypothetical protein |
| M896_060070 | 0.002786273 | Inf | Cg_06A_67 | hypothetical protein |
| M896_060080 | 0.001196581 | Inf | Cg_06A_65 | hypothetical protein |
| M896_060090 | 0.001664731 | 1.74155275 | Cg_06A_64 | transketolase |
| M896_060100 | 0.002792957 | 0.76660959 | Cg_06A_63 | hypothetical protein |
| M896_060110 | 0.004248717 | 0.50189419 | Cg_06A_62 | hypothetical protein |
| M896_060120 | 0.001317217 | 0.13156376 | Cg_06A_61 | hypothetical protein |
| M896_060130 | 0 | Inf | Cg_06A_60 | hypothetical protein |
| M896_060140 | 0.004152637 | 0.19490587 | Cg_06A_59 | hypothetical protein |
| M896_060150 | 0.002514861 | 0.20869068 | Cg_06A_58 | hypothetical protein |
| M896_060160 | 0.002279202 | 0.44438819 | Cg_06A_57 | hypothetical protein |
| M896_060170 | 0.002536783 | Inf | Cg_06A_56 | hypothetical protein |
| M896_060180 | 0.001811374 | 0.52133917 | Cg_06A_55 | hypothetical protein |
| M896_060190 | 0.004643449 | 0.27754237 | Cg_06A_54 | hypothetical protein |
| M896_060200 | 0.000899165 | 0 | Cg_06A_53 | putative ATPase |
| M896_060210 | 0.002003888 | 0.1032121 | Cg_06A_52 | Rad3-like DNA repair helicase |
| M896_060220 | 0.000770077 | 0 | Cg_06A_51 | subunit beta of translation initiation factor 2 |
| M896_060230 | 0.001969395 | 0.31638808 | Cg_06A_50 | beta-tubulin folding cofactor D |
| M896_060240 | 0.003142536 | 0.39166299 | Cg_06A_49 | hypothetical protein |
| M896_060250 | 0.001558016 | 0.19113785 | Cg_06A_48 | polar tube protein 2 |
| M896_060260 | 0.002463353 | 1.15368148 | Cg_06A_47 | polar tube protein 1 |
| M896_060270 | 0.000925926 | Inf | Cg_06A_46 | putative E2F transcription factor |
| M896_060280 | 0 | Inf | Cg_06A_45 | subunit of transport protein particle complex |
| M896_060290 | 0.006713717 | 0.2703444 | Cg_06A_44 | leucyl-tRNA synthetase |
| M896_060300 | 0.002363453 | 1.07452316 | Cg_06A_42 | hypothetical protein |
| M896_060310 | 0.003462158 | 0.72705315 | Cg_06A_41 | hypothetical protein |
| M896_060320 | 0.001414898 | 0.43352872 | Cg_06A_40 | queuine tRNA-ribosyltransferase |
| M896_060330 | 0 | Inf | Cg_06A_39 | RNA polymerases N |
| M896_060340 | 0.001782233 | Inf | Cg_06A_38 | PPIase/rotamase |
| M896_060350 | 0.001663818 | 1.19491272 | Cg_06A_37 | minichromosome maintenance protein |
| M896_060360 | 0.002469136 | 0.14871691 | Cg_06A_36 | subunit of transcription initiation factor IIE |
| M896_060370 | 0.002401488 | Inf | Cg_06A_35 | single-stranded DNA-binding replication protein |
| M896_060380 | 0.001624631 | 1.22378038 | Cg_06A_34 | hypothetical protein |
| M896_060390 | 0.005902472 | 1.32315907 | Cg_06A_33 | hypothetical protein |
| M896_060400 | 0.000350877 | Inf | Cg_06A_32 | RNase PH-like exoribonuclease |
| M896_060410 | 0.001893939 | 0.28099508 | Cg_06A_31 | Mn2+-dependent serine/threonine protein kinase |
| M896_060430 | 0.00291439 | 0.55951847 | Cg_06A_30 | subunit of dynactin complex |
| M896_060450 | 0.001341574 | 0.40288084 | Cg_06A_29 | ribosome biogenesis GTP-binding protein |
| M896_060460 | 0.002572016 | 0.30843009 | Cg_06A_28 | dUTPase |
| M896_060470 | 0.00100852 | 1.14969295 | Cg_06A_27 | hypothetical protein |
| M896_060480 | 0.00342556 | 0.63800148 | Cg_06A_26 | hypothetical protein |
| M896_060490 | 0.001006441 | Inf | Cg_06A_25 | N-acetyltransferase |
| M896_060500 | 0.001455884 | 0.37136439 | Cg_06A_24 | subunit Nup170 of nuclear pore complex |
| M896_060510 | 0.001449848 | 0.43450142 | Cg_06A_23 | Tom40-like porin |
| M896_060520 | 0.002628903 | 0.12135686 | Cg_06A_22 | subunit zeta of T-complex protein 1 |
| M896_060530 | 0.001927541 | 0.8319986 | Cg_06A_21 | hypothetical protein |
| M896_060540 | 0.000936569 | Inf | Cg_06A_20 | Tub-like protein |
| M896_060550 | 0.000942418 | 0 | Cg_06A_19 | TPR repeat-containing protein |
| M896_060560 | 0.001618837 | 0.09556787 | Cg_06A_18 | hypothetical protein |
| M896_060570 | 0.003689404 | 0.30218757 | Cg_06A_17 | hypothetical protein |
| M896_060580 | 0.002259402 | 0 | Cg_06A_16 | hypothetical protein |
| M896_060590 | 0.005333333 | 1.35234331 | Cg_06A_15 | hypothetical protein |
| M896_060600 | 0.003843123 | 0.33364757 | Cg_06A_14 | hypothetical protein |
| M896_060610 | 0 | Inf | Cg_06A_13 | subunit RPB3 of DNA-directed RNA polymerase II |
| M896_060620 | 0.004542664 | 0.23836741 | Cg_06A_12 | putative HemK-like methylase |
| M896_060630 | 0.002440492 | 1.30994292 | Cg_06A_11 | histidyl-tRNA synthetase |
| M896_060640 | 0.002421164 | 0.44756506 | Cg_06A_10 | hypothetical protein |
| M896_060650 | 0.003060546 | Inf | Cg_06A_8 | adenylate kinase |
| M896_060660 | 0.003030303 | 1.16618965 | Cg_06A_7 | hypothetical protein |
| M896_060670 | 0.002120051 | 0.15895978 | Cg_06A_6 | hypothetical protein |
| M896_060680 | 0.002039118 | 0 | Cg_06A_5 | hypothetical protein |
| M896_060690 | 0.001913171 | 0.06732145 | Cg_06A_4 | hypothetical protein |
| M896_060700 | 0.00394714 | 0.63349224 | Cg_06A_3 | hypothetical protein |
| M896_060710 | 0.002125076 | 0.46391753 | Cg_06A_2 | phosphotyrosyl phosphatase activator |
| M896_060720 | 0.001709402 | 0 | Cg_06A_1 | ribonucleoside-diphosphate reductase |
| M896_060730 | 0.00078693 | 0 | Cg_06B_2 | hypothetical protein |
| M896_060740 | 0.001986928 | 0.34240302 | Cg_06B_3 | subunit of vesicle coat complex |
| M896_060760 | 0.001758324 | 0.08136168 | Cg_06B_4 | symplekin domain-containing protein |
| M896_060770 | 0.000711359 | Inf | Cg_06B_5 | aspartyl-tRNA synthetase |
| M896_060780 | 0.001951793 | 0.21561285 | Cg_06B_6 | hypothetical protein |
| M896_060790 | 0.00242963 | 0 | Cg_06B_7 | hypothetical protein |
| M896_060800 | 0.001462348 | 0.3252666 | Cg_06B_8 | Snf2/Rad54-like helicase |
| M896_060810 | 0.001376676 | 0.14266779 | Cg_06B_9 | mRNA turnover and stability protein |
| M896_060820 | 0 | Inf | Cg_06B_10 | subunit A of RNA polymerase II |
| M896_060830 | 0.002763958 | Inf | Cg_06B_12 | hypothetical protein |
| M896_060840 | 0.001806685 | 0.33978446 | Cg_06B_13 | hypothetical protein |
| M896_060850 | 0.001972625 | 1.52461251 | Cg_06B_14 | subunit Sec63 of preprotein translocase |
| M896_060860 | 0.002395587 | 0.19810666 | Cg_06B_15 | hypothetical protein |
| M896_060870 | 0.002923977 | Inf | Cg_06B_16 | hypothetical protein |
| M896_060880 | 0.000649773 | 0 | Cg_06B_17 | ribosomal protein L5 |
| M896_060900 | 0.001137699 | Inf | Cg_06B_18 | ubiquitin C-terminal hydrolase |
| M896_060910 | 0.002842206 | 0.44414597 | Cg_06B_19 | pre-mRNA splicing helicase |
| M896_060920 | 0.002343084 | 0.14323617 | Cg_06B_20 | P-type ATPase |
| M896_060930 | 0.003506889 | 0.96998833 | Cg_06B_22 | hypothetical protein |
| M896_060940 | 0.001564114 | Inf | Cg_06B_23 | hypothetical protein |
| M896_060950 | 0.001047254 | 0.3711771 | Cg_06B_24 | hypothetical protein |
| M896_060960 | 0.002506628 | 0.54879968 | Cg_06B_25 | hypothetical protein |
| M896_060970 | 0.000688365 | Inf | Cg_06B_26 | subunit epsilon of t-complex protein 1 |
| M896_070050 | 0.000838574 | Inf | Cg_07_65 | Myb-like transcription factor |
| M896_070060 | 0.002706956 | 0.09373498 | Cg_07_64 | RNA 3'-terminal phosphate cyclase |
| M896_070070 | 0.000569801 | Inf | Cg_07_63 | ribulose-5-phosphate 3-epimerase |
| M896_070080 | 0.001555556 | Inf | Cg_07_62 | zinc finger domain-containing protein |
| M896_070090 | 0.000387597 | 0 | Cg_07_61 | ribosomal protein L19 |
| M896_070100 | 0.003042328 | 0.2969919 | Cg_07_59 | mitochondrial sulfhydryl oxidase |
| M896_070110 | 0.000878443 | 0 | Cg_07_58 | subunit of transcription initiation factor TFIIIB |
| M896_070120 | 0.000779727 | 0 | Cg_07_57 | ribosomal protein S18 |
| M896_070130 | 0.003196881 | 0 | Cg_07_56 | ribosomal protein L36 |
| M896_070140 | 0.001851852 | 0.15687811 | Cg_07_55 | hypothetical protein |
| M896_070150 | 0.002222222 | 0 | Cg_07_54 | hypothetical protein |
| M896_070160 | 0 | Inf | Cg_07_53 | small nuclear ribonucleoprotein |
| M896_070170 | 0.002624672 | Inf | Cg_07_52 | hypothetical protein |
| M896_070180 | 0.001571268 | 0.18671152 | Cg_07_51 | hypothetical protein |
| M896_070190 | 0.001851852 | Inf | Cg_07_50 | hypothetical protein |
| M896_070200 | 0.000346545 | Inf | Cg_07_49 | GTP-dependent nucleic acid-binding protein EngD |
| M896_070210 | 0.003237095 | 0.99261988 | Cg_07_48 | hypothetical protein |
| M896_070220 | 0.002266446 | Inf | Cg_07_47 | hypothetical protein |
| M896_070230 | 0.005668934 | Inf | Cg_07_46 | hypothetical protein |
| M896_070240 | 0.006385696 | Inf | Cg_07_45 | hypothetical protein |
| M896_070250 | 0.00378174 | 3.34654261 | Cg_07_44 | hypothetical protein |
| M896_070260 | 0.002203065 | 0.48298005 | Cg_07_43 | TruD-like pseudouridine synthase |
| M896_070270 | 0.002494331 | Inf | Cg_07_42 | hypothetical protein |
| M896_070280 | 0.001084838 | 1.52219787 | Cg_07_41 | ATP-dependent RNA helicase |
| M896_070300 | 0.003683684 | 0.02460154 | Cg_07_40 | hypothetical protein |
| M896_070310 | 0.000654798 | 0 | Cg_07_39 | HMG domain-containing chromatin-associated |
| M896_070320 | 0.001177146 | Inf | Cg_07_38 | hypothetical protein |
| M896_070330 | 0.002821869 | 0 | Cg_07_37 | hypothetical protein |
| M896_070340 | 0.002705542 | 0.32628173 | Cg_07_36 | ATP-binding protein |
| M896_070360 | 0.000914495 | Inf | Cg_07_35 | ATP-binding protein |
| M896_070370 | 0.000974659 | 0 | Cg_07_34 | ATP-binding protein |
| M896_070380 | 0.001122334 | 0.12375054 | Cg_07_33 | subunit DPH2 of diphthamide synthase |
| M896_070390 | 0.002825506 | 0.11768935 | Cg_07_32 | hypothetical protein |
| M896_070400 | 0.002238502 | 0.16564954 | Cg_07_31 | hypothetical protein |
| M896_070410 | 0 | Inf | Cg_07_30 | hypothetical protein |
| M896_070420 | 0.001939112 | 0 | Cg_07_29 | hypothetical protein |
| M896_070430 | 0.002809706 | 0.14628353 | Cg_07_28 | ferritin |
| M896_070440 | 0.001015808 | 0.12543646 | Cg_07_27 | hypothetical protein |
| M896_070450 | 0.000703497 | Inf | Cg_07_26 | hypothetical protein |
| M896_070460 | 0.002153316 | 0.26470588 | Cg_07_25 | hypothetical protein |
| M896_070470 | 0.001690689 | 0.66258694 | Cg_07_23 | SPX domain-containing vacuolar polyphosphate protein |
| M896_070480 | 0.001683502 | 0.5247296 | Cg_07_21 | hypothetical protein |
| M896_070490 | 0.00272073 | 0.09908695 | Cg_07_20 | translation elongation factor EF-1 alpha |
| M896_070510 | 0.005449735 | 0.18834591 | Cg_07_19 | Cdc50-like protein |
| M896_070520 | 0.002270995 | 0.35276357 | Cg_07_18 | Sec1-like vacuolar protein sorting-associated protein |
| M896_070530 | 0.001707552 | 0.47090614 | Cg_07_17 | hypothetical protein |
| M896_070540 | 0.002530864 | 1.27007364 | Cg_07_16 | L-type amino acid transporter |
| M896_070550 | 0.002459491 | Inf | Cg_07_15 | hypothetical protein |
| M896_070560 | 0.000930579 | Inf | Cg_07_13 | putative hemolysin III-like integral membrane protein |
| M896_070570 | 0.000315956 | Inf | Cg_07_12 | CCR4-NOT transcriptional regulation complex protein |
| M896_070580 | 0.000724884 | Inf | Cg_07_11 | DNA polymerase sigma |
| M896_070590 | 0 | Inf | Cg_07_10 | nucleoside diphosphate kinase |
| M896_070600 | 0.002479402 | 2.1577381 | Cg_07_9 | hypothetical protein |
| M896_070610 | 0 | Inf | Cg_07_8 | hypothetical protein |
| M896_070630 | 0.001596424 | 0.05808544 | Cg_07_6 | ER lumen protein retaining receptor |
| M896_070650 | 0.002065527 | 1.38116498 | Cg_07_5 | Psp1-like protein |
| M896_070660 | 0.003255485 | 0.49434599 | Cg_07_4 | hypothetical protein |
| M896_070670 | 0.001147275 | Inf | Cg_07_3 | hypothetical protein |
| M896_070680 | 0.003161698 | 0.8451796 | Cg_07_2 | hypothetical protein |
| M896_080010 | 0.001312336 | 0.20528192 | Cg_02C_4 | hypothetical protein |
| M896_080020 | 0.002407407 | 0.72151899 | Cg_02C_5 | hypothetical protein |
| M896_080040 | 0.001409374 | 0.88767456 | Cg_02C_7 | NGG1p interacting factor 3-like protein |
| M896_080050 | 0 | Inf | Cg_02C_8 | acidic ribosomal protein P2 |
| M896_080060 | 0.002469136 | Inf | Cg_02C_9 | hypothetical protein |
| M896_080070 | 0 | Inf | Cg_02C_10 | gene silencing histone chaperone |
| M896_080080 | 0.001134857 | 4.55032677 | Cg_02C_11 | hypothetical protein |
| M896_080090 | 0.002957907 | 0.17019296 | Cg_02C_12 | hypothetical protein |
| M896_080100 | 0 | Inf | Cg_02C_13 | histidine acid phosphatase |
| M896_080110 | 0 | Inf | Cg_02C_14 | regulatory subunit of ATP-dependent 26S proteasome |
| M896_080120 | 0.002952603 | 0.19525832 | Cg_02C_15 | hypothetical protein |
| M896_080130 | 0.001851852 | 0.41903208 | Cg_02C_16 | subunit of transcription initiation factor IIE |
| M896_080140 | 0.0023216 | 0.53054951 | Cg_02C_17 | hypothetical protein |
| M896_080160 | 0.003233392 | 0.05992848 | Cg_02C_18 | mitochondrial import inner membrane translocase |
| M896_080170 | 0.002496306 | 0.13157814 | Cg_02C_19 | hypothetical protein |
| M896_080180 | 0.004007739 | 0.68032609 | Cg_02C_20 | hypothetical protein |
| M896_080190 | 0.003261698 | 0.26571719 | Cg_02C_21 | hypothetical protein |
| M896_080200 | 0.003607085 | 0.52863215 | Cg_02C_22 | hypothetical protein |
| M896_080210 | 0.001948718 | 1.80330075 | Cg_02C_23 | hypothetical protein |
| M896_080220 | 0.000939913 | 0.24212984 | Cg_02C_24 | tRNA/rRNA cytosine-C5-methylase |
| M896_080230 | 0.001926469 | Inf | Cg_02C_25 | RING-finger domain-containing ubiquitin ligase |
| M896_080240 | 0.002013664 | Inf | Cg_02C_26 | U2 snRNP/pre-mRNA association factor |
| M896_080250 | 0.001160705 | 1.40622916 | Cg_02C_27 | separase |
| M896_080260 | 0.00257649 | Inf | Cg_02C_28 | hypothetical protein |
| M896_080270 | 0.000716846 | 0 | Cg_02C_29 | serine/threonine protein kinase |
| M896_080280 | 0.001834545 | 0.09873138 | Cg_02C_30 | hypothetical protein |
| M896_080290 | 0.001216779 | Inf | Cg_02C_31 | RNA exonuclease |
| M896_080300 | 0.002230545 | Inf | Cg_02C_32 | chromosome segregation ATPase |
| M896_080310 | 0.002614379 | Inf | Cg_02C_33 | hypothetical protein |
| M896_080320 | 0.000939746 | Inf | Cg_02C_34 | ubiquitin carboxyl-terminal hydrolase |
| M896_080330 | 0.001305404 | 0.96903232 | Cg_02C_35 | hypothetical protein |
| M896_080340 | 0.003641352 | Inf | Cg_02C_36 | hypothetical protein |
| M896_080350 | 0.000658793 | 0.40020436 | Cg_02C_37 | hypothetical protein |
| M896_080360 | 0.001947446 | Inf | Cg_02C_38 | hypothetical protein |
| M896_080370 | 0.002035002 | Inf | Cg_02C_39 | hypothetical protein |
| M896_080380 | 0.003277614 | Inf | Cg_02C_40 | ribosomal protein L7Ae |
| M896_080390 | 0.001653439 | Inf | Cg_02C_41 | hypothetical protein |
| M896_080400 | 0.004009877 | 2.45718808 | Cg_02C_43 | hypothetical protein |
| M896_080410 | 0.00167127 | 1.16243862 | Cg_02C_44 | Cdc46/Mcm ATPase |
| M896_080420 | 0.0016122 | Inf | Cg_02C_45 | hypothetical protein |
| M896_080430 | 0.002557078 | Inf | Cg_02C_46 | hypothetical protein |
| M896_080440 | 0.001355932 | 0.41096374 | Cg_02C_47 | serine/threonine kinase |
| M896_080500 | 0 | Inf | Cg_08A_90 | putative ATP binding protein |
| M896_080510 | 0.003149257 | Inf | Cg_08A_89 | hypothetical protein |
| M896_080520 | 0.003730445 | 1.51363337 | Cg_08A_88 | EPP-like transporter |
| M896_080530 | 0.00110257 | 0.46429405 | Cg_08A_87 | hypothetical protein |
| M896_080540 | 0.003132777 | Inf | Cg_08A_86 | hypothetical protein |
| M896_080550 | 0.000711446 | Inf | Cg_08A_85 | 26S proteasome regulatory complex protein |
| M896_080560 | 0.001275046 | Inf | Cg_08A_84 | adrenodoxin-like ferredoxin |
| M896_080570 | 0.002214657 | 1.66872847 | Cg_08A_83 | Rad50-like protein |
| M896_080580 | 0.004540024 | Inf | Cg_08A_82 | hypothetical protein |
| M896_080590 | 0.00123832 | 0.40407865 | Cg_08A_81 | hypothetical protein |
| M896_080600 | 0.002323893 | Inf | Cg_08A_80 | hypothetical protein |
| M896_080610 | 0 | Inf | Cg_08A_79 | hypothetical protein |
| M896_080620 | 0.001814059 | Inf | Cg_08A_78 | small ubiquitin-related modifier protein |
| M896_080630 | 0 | Inf | Cg_08A_77 | hypothetical protein |
| M896_080650 | 0.001618234 | 0.46589201 | Cg_08A_76 | chromosome segregation ATPase |
| M896_080660 | 0.00151005 | 0.30558962 | Cg_08A_75 | hypothetical protein |
| M896_080670 | 0.001590072 | Inf | Cg_08A_74 | subunit Sec62 of preprotein translocase |
| M896_080680 | 0.000727107 | Inf | Cg_08A_73 | hypothetical protein |
| M896_080690 | 0 | Inf | Cg_08A_72 | hypothetical protein |
| M896_080700 | 0 | Inf | Cg_08A_71 | hypothetical protein |
| M896_080710 | 0.002213369 | 0.16519435 | Cg_08A_70 | aquaporin |
| M896_080720 | 0.003161002 | 0.40421218 | Cg_08A_69 | hypothetical protein |
| M896_080730 | 0.000759259 | 0 | Cg_08A_68 | DnaJ-class molecular chaperone |
| M896_080740 | 0.001967112 | 3.38499302 | Cg_08A_67 | nuclear pore protein |
| M896_080750 | 0.000841751 | Inf | Cg_08A_66 | hypothetical protein |
| M896_080760 | 0.002336029 | 1.68561691 | Cg_08A_65 | hypothetical protein |
| M896_080770 | 0.000383555 | Inf | Cg_08A_64 | glyceraldehyde-3-phosphate dehydrogenase |
| M896_080780 | 0.002146733 | Inf | Cg_08A_63 | hypothetical protein |
| M896_080790 | 0.001652649 | 0.19524102 | Cg_08A_62 | acidic ribosomal protein P0 |
| M896_080800 | 0.002696388 | 0.51054665 | Cg_08A_61 | Ski2-like helicase |
| M896_080810 | 0.001638807 | Inf | Cg_08A_60 | subunit Rpb8 of RNA polymerase |
| M896_080820 | 0.001282271 | 0.85705884 | Cg_08A_58 | subunit Cullin of E3 ubiquitin ligase |
| M896_080840 | 0.000644122 | Inf | Cg_08A_55 | hypothetical protein |
| M896_080850 | 0.000789622 | 0 | Cg_08A_54 | RNA polymerase II transcription |
| M896_080870 | 0.004042162 | 1.30220027 | Cg_08A_53 | TFIIF-interacting CTD phosphatase |
| M896_080880 | 0.005152979 | 2.28452592 | Cg_08A_52 | alpha/beta hydrolase domain-containing protein |
| M896_080890 | 0.001783983 | 0.30654544 | Cg_08A_51 | hypothetical protein |
| M896_080900 | 0.003890054 | 0.17387625 | Cg_08A_50 | hypothetical protein |
| M896_080910 | 0.000694444 | Inf | Cg_08A_49 | ribosomal protein L24 |
| M896_080920 | 0.005185185 | Inf | Cg_08A_48 | putative subunit RPA3 of replication protein A |
| M896_080930 | 0.000995619 | Inf | Cg_08A_47 | subunit Rpb5 of DNA-directed RNA polymerase |
| M896_080940 | 0.002080732 | Inf | Cg_08A_46 | hypothetical protein |
| M896_080950 | 0 | Inf | Cg_08A_45 | hypothetical protein |
| M896_080960 | 0.000911321 | Inf | Cg_08A_44 | subunit DPH2 of diphthamide synthase |
| M896_080970 | 0.002436647 | Inf | Cg_08A_43 | hypothetical protein |
| M896_080980 | 0.00119911 | 0.79093608 | Cg_08A_42 | hypothetical protein |
| M896_080990 | 0.002071563 | 0.09713989 | Cg_08A_41 | hypothetical protein |
| M896_081000 | 0.000766284 | Inf | Cg_08A_40 | ribosomal protein L37a |
| M896_081010 | 0.000813224 | 0.21683115 | Cg_08A_38 | subunit POB3 of nucleosome-binding factor SPN |
| M896_081020 | 0.00295584 | 0.23952794 | Cg_08A_37 | WD40 domain-containing protein |
| M896_081030 | 0.005954416 | 0.31016049 | Cg_08A_36 | hypothetical protein |
| M896_081040 | 0.003127572 | 0.07809415 | Cg_08A_35 | subunit alpha of proteasome |
| M896_081050 | 0.003295421 | 1.26949737 | Cg_08A_34 | NAD kinase |
| M896_081060 | 0.002049957 | 0.05422188 | Cg_08A_33 | ATP-dependent RNA helicase |
| M896_081070 | 0.002288152 | Inf | Cg_08A_32 | hypothetical protein |
| M896_081080 | 0.005499439 | 0.27342366 | Cg_08A_31 | hypothetical protein |
| M896_081090 | 0.003190883 | 0.33649658 | Cg_08A_30 | hypothetical protein |
| M896_081100 | 0.001129575 | 1.03594267 | Cg_08A_29 | ATP-dependent DNA helicase RecQ |
| M896_081110 | 0.001477261 | 0.52076719 | Cg_08A_28 | subunit 3 of splicing factor 3a |
| M896_081120 | 0.001763668 | 0 | Cg_08A_27 | hypothetical protein |
| M896_081140 | 0.003840036 | 0.57106291 | Cg_08A_25 | tRNA-dihydrouridine synthase |
| M896_081150 | 0.001967593 | Inf | Cg_08A_24 | hypothetical protein |
| M896_081160 | 0.002493583 | 1.13530046 | Cg_08A_23 | alpha tubulin |
| M896_081170 | 0.003112356 | Inf | Cg_08A_22 | hypothetical protein |
| M896_081180 | 0.001601602 | Inf | Cg_08A_21 | hypothetical protein |
| M896_081190 | 0.001773781 | Inf | Cg_08A_20 | formin homology 2 domain-containing protein |
| M896_081200 | 0.001030596 | Inf | Cg_08A_19 | hypothetical protein |
| M896_081210 | 0.000995025 | Inf | Cg_08A_18 | hypothetical protein |
| M896_081220 | 0.003404415 | 0.68472085 | Cg_08A_17 | hypothetical protein |
| M896_081230 | 0.000214362 | 0 | Cg_08A_16 | Golgi nucleoside diphosphatase |
| M896_081240 | 0.003805175 | 0.85314404 | Cg_08A_15 | serine/threonine kinase |
| M896_081250 | 0.001503928 | 0.46348312 | Cg_08A_14 | glucosamine-fructose-6-phosphate |
| M896_081260 | 0.001186468 | 0 | Cg_08A_13 | hypothetical protein |
| M896_081270 | 0.004032604 | 0.66763472 | Cg_08A_12 | Hsp90 ATPase activator |
| M896_081280 | 0 | Inf | Cg_08A_11 | hypothetical protein |
| M896_081290 | 0 | Inf | Cg_08A_10 | hypothetical protein |
| M896_081300 | 0.001753697 | 0.57668126 | Cg_08A_9 | subunit beta of translation initiation factor IF-2 |
| M896_081310 | 0.002211564 | 0.42755769 | Cg_08A_8 | rRNA methylase |
| M896_081320 | 0.001010101 | 0 | Cg_08A_7 | putative V-type ATP synthase |
| M896_081330 | 0.00464135 | Inf | Cg_08A_6 | hypothetical protein |
| M896_081340 | 0.001457683 | 0.23546566 | Cg_08A_5 | hypothetical protein |
| M896_081350 | 0.002682998 | Inf | Cg_08A_3 | ubiquitin/L40 ribosomal protein fusion |
| M896_081360 | 0.001481481 | Inf | Cg_08A_1 | ribosomal protein L22/L17e |
| M896_081370 | 0.001363819 | 0.32418637 | Cg_08B_1 | beta type-1 subunit of proteasome |
| M896_081380 | 0.001359369 | 0.92737385 | Cg_08B_2 | hypothetical protein |
| M896_081390 | 0.001657414 | 0.21968222 | Cg_08B_3 | Dopey-like leucine zipper transcription factor |
| M896_081400 | 0.003898635 | Inf | Cg_08B_4 | hypothetical protein |
| M896_081410 | 0.007070707 | 0.1014245 | Cg_08B_5 | ribosomal protein L37 |
| M896_081420 | 0.001346801 | 0 | Cg_08B_6 | hypothetical protein |
| M896_081440 | 0.000758956 | Inf | Cg_08B_7 | hypothetical protein |
| M896_081460 | 0.005677534 | 0.1843736 | Cg_08B_8 | hypothetical protein |
| M896_081480 | 0.003392378 | 2.90684076 | Cg_08B_10 | type V P-ATPase |
| M896_081490 | 0.001413862 | Inf | Cg_08B_11 | hypothetical protein |
| M896_081510 | 0.003383739 | 0.84448821 | Cg_08B_16 | hypothetical protein |
| M896_081530 | 0.003703704 | 0.27319154 | Cg_08B_18 | hypothetical protein |
| M896_081540 | 0.001881246 | Inf | Cg_08B_19 | threonyl-tRNA synthetase |
| M896_081550 | 0.002645503 | Inf | Cg_08B_20 | hypothetical protein |
| M896_081560 | 0.00313647 | 1.50436823 | Cg_08B_21 | hypothetical protein |
| M896_081570 | 0.000982415 | Inf | Cg_08B_22 | hypothetical protein |
| M896_081580 | 0.002523659 | 0.88228926 | Cg_08B_23 | tRNA-dihydrouridine synthase |
| M896_081590 | 0 | Inf | Cg_08B_24 | hypothetical protein |
| M896_081600 | 0.002333841 | 0.46860137 | Cg_08B_25 | mRNA decapping enzyme 2 |
| M896_081610 | 0.000713735 | 0.19448884 | Cg_08B_26 | regulatory subunit 4 of proteosome |
| M896_081620 | 0.001332268 | Inf | Cg_08B_27 | hypothetical protein |
| M896_081630 | 0 | Inf | Cg_08B_28 | hypothetical protein |
| M896_081640 | 0.001158504 | Inf | Cg_08B_29 | subunit alpha of phenylalanyl-tRNA synthetase |
| M896_081650 | 0.001865502 | 1.94084043 | Cg_08B_30 | hypothetical protein |
| M896_081660 | 0.004640272 | Inf | Cg_08B_31 | putative acetyltransferase |
| M896_081670 | 0.002081937 | 0.49321424 | Cg_08B_32 | transcriptional accessory-like protein |
| M896_081680 | 0.003751804 | Inf | Cg_08B_33 | ribosomal protein S2 |
| M896_081690 | 0.002364066 | 0.50369932 | Cg_08B_35 | hypothetical protein |
| M896_081700 | 0 | Inf | Cg_08B_36 | hypothetical protein |
| M896_081710 | 0.00158584 | 0.57493713 | Cg_08B_37 | hypothetical protein |
| M896_081720 | 0.003902439 | Inf | Cg_08B_38 | MADS domain-containing protein |
| M896_081730 | 0 | Inf | Cg_08B_39 | subunit 7 of RNA polymerase II |
| M896_090020 | 0 | Inf | Cg_09A_3 | hypothetical protein |
| M896_090030 | 0.001234568 | Inf | Cg_09A_4 | hypothetical protein |
| M896_090040 | 0.001336767 | Inf | Cg_09A_5 | ABCG White-like ABC transporter |
| M896_090050 | 0.002366522 | 1.05939732 | Cg_09A_6 | hypothetical protein |
| M896_090060 | 0.000801833 | Inf | Cg_09A_7 | hypothetical protein |
| M896_090070 | 0.002763958 | Inf | Cg_09A_8 | putative small nuclear ribonucleoprotein |
| M896_090080 | 0.002723312 | Inf | Cg_09A_9 | hypothetical protein |
| M896_090090 | 0.002508961 | Inf | Cg_09A_10 | hypothetical protein |
| M896_090100 | 0 | Inf | Cg_09A_11 | small subunit of clathrin adaptor complex |
| M896_090110 | 0 | Inf | Cg_09A_12 | EF-Hand Ca2+-binding protein |
| M896_090120 | 0.000985389 | Inf | Cg_09A_13 | hypothetical protein |
| M896_090130 | 0.001702509 | 1.3325867 | Cg_09A_14 | hypothetical protein |
| M896_090140 | 0.002066277 | Inf | Cg_09A_15 | hypothetical protein |
| M896_090150 | 0.00546566 | 0.24980288 | Cg_09A_16 | hypothetical protein |
| M896_090160 | 0.002326363 | 1.20022626 | Cg_09A_17 | hypothetical protein |
| M896_090180 | 0.00255144 | Inf | Cg_09A_18 | cyclin-dependent protein kinase |
| M896_090190 | 0.001675485 | Inf | Cg_09A_19 | GTP-binding protein |
| M896_090200 | 0.000513809 | Inf | Cg_09A_20 | subunit H of vacuolar H+-ATPase V1 |
| M896_090210 | 0.00222484 | 0.66278091 | Cg_09A_21 | cyclin |
| M896_090220 | 0.001038961 | 0.16044191 | Cg_09A_23 | hypothetical protein |
| M896_090230 | 0.000676819 | Inf | Cg_09A_24 | beta type-2 subunit of proteasome |
| M896_090240 | 0.00138214 | 0.15561736 | Cg_09A_25 | minichromosome maintenance protein |
| M896_090250 | 0.000713558 | 0 | Cg_09A_26 | hypothetical protein |
| M896_090260 | 0.002808849 | 0.1522553 | Cg_09A_28 | hypothetical protein |
| M896_090270 | 0 | Inf | Cg_09A_29 | DNA-directed RNA polymerase |
| M896_090280 | 0.002777778 | Inf | Cg_09A_30 | subunit H of ATP synthase |
| M896_090290 | 0.003870968 | 0.1013777 | Cg_09A_31 | putative GTPase |
| M896_090300 | 0.001665781 | 0.51546696 | Cg_09A_32 | hypothetical protein |
| M896_090310 | 0.003328645 | 0.20255488 | Cg_09A_33 | ribosomal protein S15 |
| M896_090320 | 0.002496195 | 0.17856007 | Cg_09A_34 | hypothetical protein |
| M896_090330 | 0.003627075 | 0 | Cg_09A_35 | ribosomal protein L26 |
| M896_090340 | 0.000842549 | 0 | Cg_09A_36 | RNA binding protein |
| M896_090350 | 0.005282802 | 0.33974359 | Cg_09A_37 | histidine triad nucleotide-binding protein |
| M896_090360 | 0.000959954 | 0.04161079 | Cg_09A_38 | tRNA/rRNA cytosine-C5-methylase |
| M896_090370 | 0 | Inf | Cg_09A_39 | histone H2B-like protein |
| M896_090380 | 0.002696629 | 0.29970695 | Cg_09A_40 | hypothetical protein |
| M896_090390 | 0.004227181 | 1.84644178 | Cg_09A_41 | hypothetical protein |
| M896_090400 | 0 | Inf | Cg_09A_42 | hypothetical protein |
| M896_090410 | 0.001909126 | Inf | Cg_09A_43 | hypothetical protein |
| M896_090420 | 0.00162174 | Inf | Cg_09A_44 | E1 ubiquitin activating enzyme-like protein |
| M896_090430 | 0.00462963 | 0.26001284 | Cg_09A_45 | peptidyl-prolyl cis-trans isomerase |
| M896_090440 | 0.000903342 | Inf | Cg_09A_46 | hypothetical protein |
| M896_090450 | 0.000589817 | Inf | Cg_09A_47 | cysteinyl-tRNA-synthetase |
| M896_090460 | 0 | Inf | Cg_09A_48 | hypothetical protein |
| M896_090470 | 0.0022285 | 0.12843549 | Cg_09A_49 | hypothetical protein |
| M896_090480 | 0.00177703 | 0.81416054 | Cg_09A_50 | RNA binding domain-containing protein |
| M896_090490 | 0.006475183 | 0.6557543 | Cg_09A_51 | hypothetical protein |
| M896_090500 | 0 | Inf | Cg_09A_52 | Sm-like protein |
| M896_090510 | 0.000980186 | 0.47891508 | Cg_09A_53 | arginyl-tRNA synthetase |
| M896_090520 | 0.000877533 | Inf | Cg_09A_54 | Zn-finger domain-containing protein |
| M896_090560 | 0.0019696 | 0 | Cg_09A_55 | hypothetical protein |
| M896_090570 | 0.002837153 | 0.38797907 | Cg_09A_56 | hypothetical protein |
| M896_090580 | 0.002955083 | 0.03677705 | Cg_09A_57 | ribosomal protein L18 |
| M896_090590 | 0.002185792 | 0.268133 | Cg_09A_58 | chromosome condensation complex Condensin |
| M896_090600 | 0.002896407 | 0.43263151 | Cg_09A_59 | hypothetical protein |
| M896_090610 | 0.001936919 | 0.53058292 | Cg_09A_60 | catalytic subunit of DNA primase |
| M896_090620 | 0.001976977 | 0.66432036 | Cg_09A_62 | hypothetical protein |
| M896_090630 | 0.004063492 | 0.87872966 | Cg_09A_63 | subunit E' of DNA-directed RNA polymerase |
| M896_090640 | 0.000732601 | Inf | Cg_09A_65 | hypothetical protein |
| M896_090660 | 0.002010582 | 0.7438291 | Cg_09A_66 | gamma-tubulin |
| M896_090670 | 0.001003584 | 0 | Cg_09A_67 | subunit zeta of vesicle coat complex COPI |
| M896_090680 | 0.000647501 | Inf | Cg_09A_68 | hypothetical protein |
| M896_090690 | 0 | Inf | Cg_09A_69 | Golgi-to-ER retrieval protein |
| M896_090700 | 0.002629544 | 0.44305273 | Cg_09A_70 | hypothetical protein |
| M896_090710 | 0.002942097 | 0.32092215 | Cg_09A_71 | hypothetical protein |
| M896_090720 | 0.000614086 | Inf | Cg_09A_73 | Rab5-like GTPase |
| M896_090730 | 0.002988954 | Inf | Cg_09A_74 | MRP-like protein |
| M896_090740 | 0.003144455 | 0.66873447 | Cg_09A_75 | ERCC4-type nuclease |
| M896_090750 | 0.004368471 | Inf | Cg_09A_76 | subunit Rpb4 of RNA polymerase II |
| M896_090760 | 0.001678657 | Inf | Cg_09A_77 | 8-oxoguanine DNA glycosylase |
| M896_090770 | 0.003113733 | 7.13009756 | Cg_09A_78 | hypothetical protein |
| M896_090780 | 0.005023414 | 0.93991313 | Cg_09A_79 | hypothetical protein |
| M896_090790 | 0.000925926 | Inf | Cg_09A_80 | signal recognition particle protein Srp19 |
| M896_090800 | 0.002382194 | Inf | Cg_09A_81 | hypothetical protein |
| M896_090810 | 0.000787037 | 0 | Cg_09A_82 | ribosomal protein L4 |
| M896_090820 | 0.001880262 | 0.87688507 | Cg_09A_83 | hypothetical protein |
| M896_090830 | 0.000945626 | Inf | Cg_09A_84 | subunit of mRNA deadenylase |
| M896_090840 | 0.001346801 | Inf | Cg_09A_85 | ubiquitin-conjugating enzyme E2 |
| M896_090850 | 0.000713967 | Inf | Cg_09A_86 | ribosomal protein S4 |
| M896_090860 | 0.003198653 | 0.51523968 | Cg_09A_87 | endonuclease III |
| M896_090870 | 0.001434943 | Inf | Cg_09A_88 | kinesin-like protein |
| M896_090880 | 0.002103338 | Inf | Cg_09A_90 | translation initiation factor 4E |
| M896_090890 | 0.000690989 | 0.15270822 | Cg_09A_91 | hypothetical protein |
| M896_090900 | 0.000493827 | 0 | Cg_09A_92 | hypothetical protein |
| M896_090910 | 0 | Inf | Cg_09A_93 | cyclic nucleotide-binding domain-containing protein |
| M896_090940 | 0.001559454 | Inf | Cg_09A_94 | hypothetical protein |
| M896_090950 | 0.000894479 | Inf | Cg_09A_95 | hypothetical protein |
| M896_090960 | 0.000252525 | 0 | Cg_09A_96 | heat shock transcription factor |
| M896_090970 | 0.002668071 | Inf | Cg_09A_97 | hypothetical protein |
| M896_090990 | 0.000958056 | Inf | Cg_09A_98 | hypothetical protein |
| M896_091000 | 0.002277396 | Inf | Cg_09A_99 | hypothetical protein |
| M896_091010 | 0.002032908 | 0.25277391 | Cg_09A_100 | hypothetical protein |
| M896_091020 | 0.003703704 | 0.11340051 | Cg_09A_101 | hypothetical protein |
| M896_091030 | 0.000868455 | Inf | Cg_09A_102 | hypothetical protein |
| M896_091050 | 0.000890313 | Inf | Cg_09A_107 | hypothetical protein |
| M896_091060 | 0.001217656 | Inf | Cg_09A_106 | ribosomal protein S13 |
| M896_091080 | 0.001699083 | 0.10968764 | Cg_09B_2 | ATP-dependent RNA helicase |
| M896_091090 | 0.001081081 | 0.29887445 | Cg_09B_3 | histone acetyltransferase |
| M896_091100 | 0.003265048 | 0.40378022 | Cg_09B_4 | subunit beta of coatomer complex |
| M896_091110 | 0.000199005 | 0 | Cg_09B_5 | WD40 domain-containing protein |
| M896_091120 | 0.001017812 | 0.15043773 | Cg_09B_6 | Rad3-like DNA helicase |
| M896_091130 | 0.004232804 | 0.16168627 | Cg_09B_7 | hypothetical protein |
| M896_091140 | 0.003776325 | 0.13399148 | Cg_09B_8 | hypothetical protein |
| M896_091150 | 0.000429415 | Inf | Cg_09B_9 | putative short-chain alcohol dehydrogenase |
| M896_091160 | 0.003135983 | Inf | Cg_09B_10 | ribosomal protein L23 |
| M896_091170 | 0.002962963 | Inf | Cg_09B_11 | hypothetical protein |
| M896_091180 | 0.003734313 | Inf | Cg_09B_12 | hypothetical protein |
| M896_091190 | 0.003105481 | 0.72298875 | Cg_09B_13 | hypothetical protein |
| M896_091200 | 0.002684336 | 0.49267902 | Cg_09B_14 | putative subunit p30 of RNase P/RNase MRP |
| M896_091210 | 0.007611281 | 0.47312754 | Cg_09B_15 | putative subunit p30 of RNase P/RNase MRP |
| M896_091220 | 0.002783675 | 0.07976026 | Cg_09B_16 | hypothetical protein |
| M896_091230 | 0.001693122 | 0 | Cg_09B_17 | hypothetical protein |
| M896_091250 | 0.002796847 | 0.89116303 | Cg_09B_18 | hypothetical protein |
| M896_091260 | 0.001634738 | 0.07569346 | Cg_09B_19 | hypothetical protein |
| M896_091270 | 0.002260235 | 0.43842744 | Cg_09B_20 | WD40 domain-containing protein |
| M896_091280 | 0.002759206 | 1.10830517 | Cg_09B_21 | hypothetical protein |
| M896_091290 | 0.001877359 | 0 | Cg_09B_22 | developmentally regulated GTP-binding protein |
| M896_091300 | 0.001346801 | 0.35169965 | Cg_09B_23 | hypothetical protein |
| M896_091310 | 0.003574368 | 0.03211918 | Cg_09B_24 | subunit beta of prenyltransferase |
| M896_091320 | 0.001221293 | 0.08338103 | Cg_09B_25 | ATP/ADP translocase |
| M896_091330 | 0.002855868 | 0.2253937 | Cg_09B_26 | hypothetical protein |
| M896_091340 | 0.001209373 | 0 | Cg_09B_27 | subunit M of DNA-directed RNA polymerase |
| M896_091350 | 0.002145215 | 0.16836989 | Cg_09B_28 | hypothetical protein |
| M896_091360 | 0.002494591 | 0.26797386 | Cg_09B_29 | TRAM protein transporter |
| M896_091370 | 0.009710551 | 0.90605229 | Cg_09B_30 | glutaredoxin |
| M896_091380 | 0.004983786 | 0.49764023 | Cg_09B_31 | GATA Zn-finger-containing transcription |
| M896_091390 | 0.000794786 | Inf | Cg_09B_32 | hypothetical protein |
| M896_091400 | 0.00144966 | 2.05406004 | Cg_09B_33 | phosphoinositide polyphosphatase |
| M896_091410 | 0.002798354 | 0.10285714 | Cg_09B_34 | hypothetical protein |
| M896_091420 | 0.001263962 | Inf | Cg_09B_35 | hypothetical protein |
| M896_091430 | 0.003143275 | 0.34453842 | Cg_09B_36 | hypothetical protein |
| M896_091440 | 0.002901519 | 0.16345535 | Cg_09B_37 | N2%2CN2-dimethylguanosine tRNA methyltransferase |
| M896_091450 | 0.00682121 | 0.41007874 | Cg_09B_38 | hypothetical protein |
| M896_091460 | 0.004088615 | 2.69421615 | Cg_09B_39 | serine/threonine kinase |
| M896_091470 | 0.004265448 | 0.42477208 | Cg_09B_40 | hypothetical protein |
| M896_091480 | 0.001698947 | 0.30365449 | Cg_09B_41 | hypothetical protein |
| M896_091490 | 0.003161123 | 0.11773733 | Cg_09B_42 | ribosome biogenesis regulatory protein |
| M896_091500 | 0.002890696 | 0.41843734 | Cg_09B_43 | vacuolar-type H+-ATPase |
| M896_091510 | 0.003312073 | 0.12414398 | Cg_09B_44 | putative pseudouridylate synthase |
| M896_091520 | 0.001146953 | 0 | Cg_09B_45 | hypothetical protein |
| M896_091530 | 0.005331089 | 0.69191686 | Cg_09B_47 | hypothetical protein |
| M896_091540 | 0.001427115 | 0 | Cg_09B_48 | ribosomal protein L10 |
| M896_091550 | 0.000949668 | 0 | Cg_09B_49 | subunit alpha of type-2 proteasome |
| M896_091560 | 0.002539182 | 0.61781241 | Cg_09B_50 | hypothetical protein |
| M896_091570 | 0.000223199 | Inf | Cg_09B_51 | hypothetical protein |
| M896_091580 | 0.002579365 | 1.91488438 | Cg_09B_52 | putative subunit of exosome |
| M896_091590 | 0.002907208 | 0 | Cg_09B_53 | serine/threonine kinase |
| M896_091600 | 0.002026555 | 0.32655215 | Cg_09B_54 | cleavage and polyadenylation specificity factor |
| M896_091610 | 0.000496475 | Inf | Cg_09B_55 | chromatin remodeling bromodomain-containing protein |
| M896_091620 | 0.006251338 | 0.88606341 | Cg_09B_56 | hypothetical protein |
| M896_091630 | 0 | Inf | Cg_09B_57 | transcription elongation factor SPT4 |
| M896_091640 | 0.001413627 | Inf | Cg_09B_58 | hypothetical protein |
| M896_091650 | 0.002578068 | 0.10279322 | Cg_09B_59 | hypothetical protein |
| M896_091660 | 0.00213791 | 0.46938776 | Cg_09B_60 | GTPase-activating protein |
| M896_091670 | 0.001957242 | 0.08140234 | Cg_09B_61 | hypothetical protein |
| M896_091680 | 0.001680486 | Inf | Cg_09B_62 | hypothetical protein |
| M896_091690 | 0.001977066 | 0.18810166 | Cg_09B_64 | hypothetical protein |
| M896_091700 | 0.002279202 | 0.1208802 | Cg_09B_65 | subunit of RNA polymerase III transcription factor IIIC |
| M896_091710 | 0.004008715 | 0.10681147 | Cg_09B_66 | ribosomal protein L14E/L6E/L27E |
| M896_091720 | 0.003691718 | 0.26964631 | Cg_09B_67 | cyclin-dependent protein kinase |
| M896_091730 | 0.002234994 | Inf | Cg_09B_69 | hypothetical protein |
| M896_091740 | 0.001953602 | Inf | Cg_09B_70 | hypothetical protein |
| M896_091750 | 0.001335241 | Inf | Cg_09B_71 | putative PP-loop ATPase |
| M896_091760 | 0.00085676 | Inf | Cg_09B_72 | glucose-6-phosphate 1-dehydrogenase |
| M896_091780 | 0.001298929 | Inf | Cg_09B_73 | subunit beta of type-5 proteasome |
| M896_091790 | 0.001257862 | 0.68961614 | Cg_09B_74 | hypothetical protein |
| M896_091800 | 0.001057348 | 0.75541744 | Cg_09B_77 | AAA ATPase |
| M896_091830 | 0.000246002 | Inf | Cg_09C_1 | cyclin-dependent protein kinase |
| M896_091840 | 0.002237654 | Inf | Cg_09C_3 | ADP-ribosylation factor 1 |
| M896_091850 | 0.002095832 | 1.05904625 | Cg_09C_5 | hypothetical protein |
| M896_091860 | 0.00079905 | Inf | Cg_09C_6 | cyclin-dependent protein kinase |
| M896_091880 | 0.001414359 | 1.60717045 | Cg_09C_8 | hypothetical protein |
| M896_091890 | 0.001030169 | 0 | Cg_09C_9 | hypothetical protein |
| M896_091900 | 0.000750117 | Inf | Cg_09C_10 | ribosomal protein L12 |
| M896_091910 | 0.001224612 | 0.40450508 | Cg_09C_11 | hypothetical protein |
| M896_091920 | 0.001040366 | Inf | Cg_09C_12 | hypothetical protein |
| M896_091930 | 0.003174603 | 0.23098395 | Cg_09C_13 | hypothetical protein |
| M896_091940 | 0.002527233 | Inf | Cg_09C_14 | hypothetical protein |
| M896_100020 | 0.000151172 | Inf | Cg_10_6 | hypothetical protein |
| M896_100030 | 0.003876774 | 1.46447334 | Cg_10_7 | hypothetical protein |
| M896_100040 | 0.001221293 | Inf | Cg_10_8 | hypothetical protein |
| M896_100050 | 0 | Inf | Cg_10_9 | hypothetical protein |
| M896_100060 | 0.002082948 | 2.25408652 | Cg_10_10 | hypothetical protein |
| M896_100070 | 0.006045126 | 0.55493063 | Cg_10_11 | RING-finger-containing E3 ubiquitin ligase |
| M896_100080 | 0 | Inf | Cg_10_12 | hypothetical protein |
| M896_100090 | 0.000972101 | 0.23723615 | Cg_10_13 | pyruvate kinase |
| M896_100100 | 0.001210361 | Inf | Cg_10_14 | hypothetical protein |
| M896_100110 | 0.000755124 | Inf | Cg_10_15 | hypothetical protein |
| M896_100120 | 0.001881624 | Inf | Cg_10_16 | histone deacetylase |
| M896_100130 | 0.003886777 | 0.22856188 | Cg_10_17 | hypothetical protein |
| M896_100140 | 0.000973236 | 0.2543235 | Cg_10_18 | hypothetical protein |
| M896_100150 | 0.001556178 | 0.49456522 | Cg_10_19 | Prp40-like splicing factor |
| M896_100160 | 0.001637427 | 0.24844857 | Cg_10_20 | hypothetical protein |
| M896_100170 | 0.000751476 | Inf | Cg_10_21 | beta type-7 subunit of proteasome |
| M896_100180 | 0.004557823 | 0.33592538 | Cg_10_22 | hypothetical protein |
| M896_100190 | 0.001863499 | 0.08310029 | Cg_10_23 | hypothetical protein |
| M896_100200 | 0.002867384 | 0.23287671 | Cg_10_24 | hypothetical protein |
| M896_100210 | 0.000801716 | 0 | Cg_10_25 | 5'-3' exonuclease |
| M896_100220 | 0.004489867 | 0.36954529 | Cg_10_26 | subunit B of DNA polymerase delta |
| M896_100230 | 0.005041152 | 0.25003696 | Cg_10_28 | hypothetical protein |
| M896_100240 | 0.003966689 | 0.68065309 | Cg_10_29 | hypothetical protein |
| M896_100250 | 0 | Inf | Cg_10_30 | hypothetical protein |
| M896_100260 | 0.002751654 | 0.7332372 | Cg_10_31 | hypothetical protein |
| M896_100270 | 0.00152587 | 0.16806278 | Cg_10_32 | septin |
| M896_100280 | 0.002540705 | 0.2681189 | Cg_10_33 | ribonuclease |
| M896_100290 | 0.001607268 | Inf | Cg_10_34 | hypothetical protein |
| M896_100300 | 0 | Inf | Cg_10_35 | hypothetical protein |
| M896_100310 | 0.006643757 | 0.05771608 | Cg_10_36 | hypothetical protein |
| M896_100320 | 0.001678657 | 0.10004997 | Cg_10_37 | hypothetical protein |
| M896_100330 | 0.003059244 | 0.20875902 | Cg_10_38 | hypothetical protein |
| M896_100340 | 0.001647319 | 0.16701089 | Cg_10_39 | hypothetical protein |
| M896_100350 | 0.003284072 | 1.04529593 | Cg_10_40 | hypothetical protein |
| M896_100360 | 0.001893004 | 0.12665536 | Cg_10_41 | hypothetical protein |
| M896_100370 | 0.001008109 | 0.32444896 | Cg_10_42 | deoxyhypusine synthase |
| M896_100380 | 0.001723356 | 0.35443643 | Cg_10_43 | 23S rRNA methylase |
| M896_100390 | 0.002405002 | 0 | Cg_10_44 | Yos1-like protein |
| M896_100400 | 0.002673893 | 0 | Cg_10_45 | Rad52 recombination DNA repair protein |
| M896_100410 | 0.002165909 | 0.27717842 | Cg_10_46 | ER to Golgi transport membrane protein |
| M896_100420 | 0.002043423 | Inf | Cg_10_47 | hypothetical protein |
| M896_100430 | 0.00497385 | 0.25464019 | Cg_10_48 | hypothetical protein |
| M896_100440 | 0.003849903 | 0.17264032 | Cg_10_49 | hypothetical protein |
| M896_100450 | 0.003543743 | 0.08346759 | Cg_10_50 | PP2A-like phosphoprotein phosphatase |
| M896_100460 | 0.000848176 | 0.59684721 | Cg_10_51 | hypothetical protein |
| M896_100480 | 0.000749064 | Inf | Cg_10_53 | hypothetical protein |
| M896_100490 | 0.001569366 | Inf | Cg_10_54 | hypothetical protein |
| M896_100500 | 0 | Inf | Cg_10_55 | cyclin |
| M896_100510 | 0.002222222 | 0.08793201 | Cg_10_56 | subunit of pyruvate dehydrogenase E1 |
| M896_100520 | 0.001602772 | Inf | Cg_10_57 | hypothetical protein |
| M896_100530 | 0.002251271 | 0.41674717 | Cg_10_58 | hypothetical protein |
| M896_100540 | 0.00308642 | Inf | Cg_10_59 | glycosylphosphatidylinositol transamidase |
| M896_100550 | 0.003251215 | Inf | Cg_10_60 | hypothetical protein |
| M896_100560 | 0 | Inf | Cg_10_61 | hypothetical protein |
| M896_100570 | 0 | Inf | Cg_10_62 | hypothetical protein |
| M896_100580 | 0.001224612 | 0.37284961 | Cg_10_63 | hypothetical protein |
| M896_100590 | 0.000973545 | Inf | Cg_10_64 | hypothetical protein |
| M896_100600 | 0.00063234 | Inf | Cg_10_65 | hypothetical protein |
| M896_110010 | 0 | Inf | Cg_11A_16 | hypothetical protein |
| M896_110020 | 0.004433078 | 2.41659755 | Cg_11A_15 | Ndc80p-complex mitotic spindle protein |
| M896_110030 | 0.002984279 | 0.78466503 | Cg_11A_14 | asparaginyl-tRNA synthetase |
| M896_110040 | 0.002615999 | 0.77969599 | Cg_11A_13 | hypothetical protein |
| M896_110050 | 0.001054614 | 0.81537753 | Cg_11A_11 | hypothetical protein |
| M896_110060 | 0.001827291 | 0.47972693 | Cg_11A_9 | autoantigen NGP-1 |
| M896_110070 | 0.00154321 | 0.65607164 | Cg_11A_8 | ATP-dependent RNA helicase |
| M896_110080 | 0.000831444 | 0.08908026 | Cg_11A_7 | hypothetical protein |
| M896_110090 | 0.003239875 | 2.05916447 | Cg_11A_6 | DNA repair flap endonuclease |
| M896_110100 | 0.002636535 | Inf | Cg_11A_5 | subunit Tfb5 of transcription factor TFIIH complex |
| M896_110110 | 0.002112933 | Inf | Cg_11A_4 | translation factor |
| M896_110120 | 0.001440329 | Inf | Cg_11A_3 | ribosomal biogenesis protein |
| M896_110130 | 0.002297485 | Inf | Cg_11A_18 | Yip1-interacting factor |
| M896_110140 | 0.001222987 | Inf | Cg_11B_1 | vacuole import and degradation protein |
| M896_110150 | 0.000428528 | Inf | Cg_11B_2 | NADPH cytochrome p450 reductase |
| M896_110160 | 0.002065336 | 2.5090022 | Cg_11B_3 | hypothetical protein |
| M896_110170 | 0 | Inf | Cg_11B_4 | Cdc73-like RNA polymerase II accessory factor |
| M896_110180 | 0.001429767 | 1.14916899 | Cg_11B_5 | hypothetical protein |
| M896_110190 | 0.003727806 | 0.48244477 | Cg_11B_6 | putative methyl transferase |
| M896_110200 | 0.005282332 | 0.67897912 | Cg_11B_8 | hypothetical protein |
| M896_110210 | 0.001685824 | Inf | Cg_11B_9 | hypothetical protein |
| M896_110230 | 0.001756614 | 0.06719054 | Cg_11B_10 | subunit I of vacuolar-type H+-ATPase |
| M896_110240 | 0.00143636 | 0.19131139 | Cg_11B_11 | DNA helicase |
| M896_110250 | 0.006304177 | 0.17512438 | Cg_11B_12 | small nuclear ribonucleoprotein |
| M896_110260 | 0.002046647 | 0.40130043 | Cg_11B_13 | hypothetical protein |
| M896_110270 | 0.001010101 | 0 | Cg_11B_14 | hypothetical protein |
| M896_110280 | 0.002079272 | 0.39285505 | Cg_11B_15 | hypothetical protein |
| M896_110290 | 0.00039784 | 0 | Cg_11B_16 | regulatory subunit of ATP-dependent 26S proteasome |
| M896_110310 | 0.002124834 | Inf | Cg_11B_17 | PHD zinc finger domain-containing protein |
| M896_110320 | 0.001761606 | 0.67066715 | Cg_11B_18 | subunit IKAP of IkappaB kinase complex |
| M896_110330 | 0.000991543 | 0.65651103 | Cg_11B_19 | undecaprenyl pyrophosphate synthase |
| M896_110340 | 0.002353182 | Inf | Cg_11B_20 | hypothetical protein |
| M896_110350 | 0.001833517 | Inf | Cg_11B_21 | DNA/RNA helicase |
| M896_110360 | 0.00080558 | Inf | Cg_11B_22 | hypothetical protein |
| M896_110370 | 0.000478412 | Inf | Cg_11B_23 | structural maintenance of chromosomes protein |
| M896_110380 | 0.002979066 | 0.19538984 | Cg_11B_24 | hypothetical protein |
| M896_110390 | 0.000986414 | Inf | Cg_11B_25 | hypothetical protein |
| M896_110400 | 0.000766284 | 0 | Cg_11B_26 | hypothetical protein |
| M896_110410 | 0.001045752 | Inf | Cg_11B_27 | membrane traffic protein |
| M896_110420 | 0.002763958 | Inf | Cg_11B_28 | deoxycytidylate deaminase |
| M896_110430 | 0.00082811 | 1.40684713 | Cg_11B_29 | myosin heavy chain |
| M896_110440 | 0.001040366 | Inf | Cg_11B_30 | hypothetical protein |
| M896_110450 | 0.004702628 | 1.29836863 | Cg_11B_31 | hypothetical protein |
| M896_110460 | 0.002259492 | Inf | Cg_11B_32 | hypothetical protein |
| M896_120010 | 0 | Inf | Cg_12A_5 | hypothetical protein |
| M896_120020 | 0.000291918 | 0 | Cg_12A_6 | protein kinase domain-containing protein |
| M896_120040 | 0.006418002 | 0.16363744 | Cg_12A_8 | putative transcription regulator protein |
| M896_120050 | 0.000569801 | Inf | Cg_12A_9 | hypothetical protein |
| M896_120070 | 0.000733753 | Inf | Cg_12A_11 | putative CDK-activating kinase assembly factor |
| M896_120080 | 0 | Inf | Cg_12A_12 | ribosomal protein S21e |
| M896_120090 | 0.002083333 | 0.52362771 | Cg_12A_13 | triosephosphate isomerase |
| M896_120100 | 0.002319783 | 0.06642467 | Cg_12A_14 | putative exportin 1 |
| M896_120110 | 0 | Inf | Cg_12A_15 | dephospho-CoA kinase |
| M896_120120 | 0.004047346 | 0.15987362 | Cg_12A_16 | hypothetical protein |
| M896_120130 | 0.001686835 | 0.04663905 | Cg_12A_17 | hypothetical protein |
| M896_120150 | 0.005228758 | 0.8348153 | Cg_12A_19 | hypothetical protein |
| M896_120160 | 0.002815485 | 0.7708544 | Cg_12A_20 | zinc finger domain-containing protein |
| M896_120170 | 0.001468891 | 0.45053129 | Cg_12A_21 | putative RAB escort protein |
| M896_120180 | 0.004018913 | 0.23761875 | Cg_12A_22 | putative Rad5p-binding protein |
| M896_120200 | 0.005144033 | Inf | Cg_12A_24 | hypothetical protein |
| M896_120220 | 0.000745551 | 0.49759641 | Cg_12A_25 | subunit SEC6 of exocyst complex |
| M896_120230 | 0.000984529 | Inf | Cg_12A_26 | putative transmembrane adaptor Erv26 |
| M896_120240 | 0.000444444 | 0 | Cg_12A_27 | hypothetical protein |
| M896_120260 | 0.001704051 | Inf | Cg_12A_28 | hypothetical protein |
| M896_120270 | 0.002469136 | 0.1158648 | Cg_12A_29 | Sel1 repeat domain-containing protein |
| M896_120280 | 0.001786183 | Inf | Cg_12A_30 | RNP domain-containing protein |
| M896_120290 | 0.002974303 | 0.20565626 | Cg_12A_31 | phosphoinositide 4-kinase |
| M896_120300 | 0.003631436 | 0.41680365 | Cg_12A_32 | ribosomal biogenesis protein |
| M896_120310 | 0.001624021 | 0.80446387 | Cg_12A_33 | kinesin motor domain-containing protein |
| M896_120320 | 0.001844968 | 0.44014178 | Cg_12A_34 | GATase1 CTP synthase |
| M896_120330 | 0.004994071 | 0.44175555 | Cg_12A_35 | hypothetical protein |
| M896_120340 | 0.004257131 | 0.08260292 | Cg_12A_36 | RNP domain-containing protein |
| M896_120360 | 0.001562744 | Inf | Cg_12A_37 | polysaccharide deacetylase domain-containing protein |
| M896_120370 | 0.002757017 | Inf | Cg_12A_38 | hypothetical protein |
| M896_120380 | 0.004112474 | Inf | Cg_12A_39 | hypothetical protein |
| M896_120390 | 0.001387614 | Inf | Cg_12A_41 | aminoacyl-tRNA ligase |
| M896_120400 | 0.003124116 | 0.72665374 | Cg_12A_42 | Hsp70-like protein |
| M896_120410 | 0.001968635 | 1.33885438 | Cg_12A_43 | putative Arf GTPase activating protein |
| M896_120420 | 0.005913477 | 0.39349407 | Cg_12A_44 | hypothetical protein |
| M896_120430 | 0.001971756 | Inf | Cg_12A_45 | regulatory subunit of proteasome |
| M896_120440 | 0.005506391 | Inf | Cg_12A_47 | hypothetical protein |
| M896_120450 | 0.002258356 | 0 | Cg_12A_48 | subunit NuA4 of histone acetyltransferase |
| M896_120460 | 0.000673401 | 0 | Cg_12A_49 | subunit AC19 of RNA polymerase |
| M896_120470 | 0.003416797 | 0.24607309 | Cg_12A_50 | hypothetical protein |
| M896_120480 | 0.001123196 | 4.13668242 | Cg_12A_51 | pre-mRNA cleavage and polyadenylation |
| M896_120490 | 0.003003003 | 0.25589289 | Cg_12A_52 | CDP-alcohol phosphatidyltransferase |
| M896_120500 | 0.003348943 | 0.242397 | Cg_12A_53 | hypothetical protein |
| M896_120510 | 0.002469136 | Inf | Cg_12A_54 | hypothetical protein |
| M896_120520 | 0.006046863 | 0.96079545 | Cg_12A_55 | inositol metabolism VAMP-associated protein |
| M896_120530 | 0.001395231 | Inf | Cg_12A_56 | PP1 serine/threonine phosphatase |
| M896_120540 | 0.001557632 | 0 | Cg_12A_57 | eukaryotic translation initiation factor eIF2A |
| M896_120550 | 0.002386279 | 0.86104317 | Cg_12A_58 | hypothetical protein |
| M896_120560 | 0.00244606 | Inf | Cg_12A_59 | GDP-mannose pyrophosphorylase |
| M896_120570 | 0.003071364 | Inf | Cg_12A_60 | hypothetical protein |
| M896_120580 | 0.000629882 | Inf | Cg_12A_61 | hypothetical protein |
| M896_120590 | 0.002209227 | 0.09866137 | Cg_12A_62 | ribosomal protein S10 |
| M896_120600 | 0.00182716 | Inf | Cg_12A_63 | class 2 transcription repressor NC2 beta |
| M896_120610 | 0.002962963 | 0.23691099 | Cg_12A_64 | hypothetical protein |
| M896_120620 | 0.000364299 | Inf | Cg_12A_65 | putative RNA-binding protein |
| M896_120630 | 0.002992677 | 0.52372251 | Cg_12A_66 | putative exonuclease |
| M896_120650 | 0.002262922 | 3.76283847 | Cg_12A_67 | putative mRNA deadenylase |
| M896_120660 | 0.001583974 | Inf | Cg_12A_68 | hypothetical protein |
| M896_120670 | 0.001654846 | 0.57960868 | Cg_12A_70 | Sec23-like protein transport protein |
| M896_120680 | 0.002333042 | 1.82391865 | Cg_12A_72 | minichromosome maintenance protein |
| M896_120690 | 0.001493429 | Inf | Cg_12A_73 | putative ribosomal protein S8 |
| M896_120700 | 0.001705948 | 0.11396287 | Cg_12A_74 | DNA repair protein Rad51 |
| M896_120710 | 0.004122991 | 0.04801753 | Cg_12A_75 | subunit of transcription initiation factor TFIID |
| M896_120720 | 0.002409278 | 0.09895002 | Cg_12A_76 | AAA family ATPase |
| M896_120730 | 0.001572062 | 0.18274818 | Cg_12A_77 | HrpA-like helicase |
| M896_120750 | 0.003210641 | 0.30641974 | Cg_12A_78 | putative ethanolamine-phosphate |
| M896_120770 | 0.001679747 | 0.10527888 | Cg_12A_79 | methionyl-tRNA synthetase |
| M896_120780 | 0.003033276 | 0.26292482 | Cg_12A_80 | subunit of U2 snRNP spliceosome |
| M896_120790 | 0.003183886 | 0.42799172 | Cg_12A_81 | hypothetical protein |
| M896_120800 | 0.001251251 | Inf | Cg_12A_82 | hypothetical protein |
| M896_120810 | 0.003569512 | 0.62756968 | Cg_12A_83 | subunit alpha of translation initiation factor 2 |
| M896_120820 | 0.002893519 | 0 | Cg_12A_84 | hypothetical protein |
| M896_120830 | 0.002353756 | Inf | Cg_12A_85 | HS6-type ribosomal protein |
| M896_120840 | 0.000828801 | Inf | Cg_12A_86 | hypothetical protein |
| M896_120850 | 0.000723851 | Inf | Cg_12A_87 | serine/threonine kinase |
| M896_120860 | 0.001493931 | 0 | Cg_12A_88 | hypothetical protein |
| M896_120870 | 0.007446394 | 0.84294719 | Cg_12A_89 | putative synaptobrevin/VAMP-like protein |
| M896_120880 | 0.004109589 | 0.06348253 | Cg_12A_90 | hypothetical protein |
| M896_120890 | 0.004270153 | 0.25406504 | Cg_12A_91 | hypothetical protein |
| M896_120900 | 0.003145611 | 0.47838328 | Cg_12A_92 | hypothetical protein |
| M896_120910 | 0.004890918 | 0.30091503 | Cg_12A_93 | hypothetical protein |
| M896_120920 | 0.002276508 | 0.06716124 | Cg_12A_94 | putative AAA+ class ATPase |
| M896_120930 | 0.002420721 | Inf | Cg_12A_95 | hypothetical protein |
| M896_120940 | 0.003217841 | 0.57091179 | Cg_12A_96 | putative proton-dependent oligopeptide transport protein |
| M896_120950 | 0.002816505 | 0.27669903 | Cg_12A_97 | hypothetical protein |
| M896_120960 | 0.00177884 | 0.28380827 | Cg_12A_98 | hypothetical protein |
| M896_120970 | 0.00215911 | 0 | Cg_12A_100 | superoxide dismutase |
| M896_120980 | 0.001730703 | 0.42217082 | Cg_12A_101 | hypothetical protein |
| M896_120990 | 0.002521823 | 0.12044006 | Cg_12A_102 | isoleucyl-tRNA synthetase |
| M896_121000 | 0.002298851 | 0.81953187 | Cg_12A_103 | hypothetical protein |
| M896_121010 | 0.002625272 | 0.41758265 | Cg_12A_104 | translation elongation factor 2 |
| M896_121020 | 0.002962963 | 0.422323 | Cg_12A_105 | hypothetical protein |
| M896_121030 | 0.004086845 | Inf | Cg_12A_106 | putative membrane protein |
| M896_121040 | 0.000356779 | 0 | Cg_12A_107 | hypothetical protein |
| M896_121050 | 0.005023205 | 0.50447079 | Cg_12A_108 | hypothetical protein |
| M896_121060 | 0.002102102 | 0.39762686 | Cg_12A_109 | putative nucleotide binding protein |
| M896_121070 | 0.004375951 | 0.10746248 | Cg_12A_110 | putative mitochondrial ABC transporter |
| M896_121080 | 0.001585243 | 0.31481744 | Cg_12A_112 | hypothetical protein |
| M896_121090 | 0.001310541 | 0.34477555 | Cg_12A_113 | hypothetical protein |
| M896_121100 | 0.002384359 | 0.27986106 | Cg_12A_114 | hypothetical protein |
| M896_121110 | 0.001579521 | 0.31348499 | Cg_12A_115 | hypothetical protein |
| M896_121120 | 0 | Inf | Cg_12A_116 | hypothetical protein |
| M896_121140 | 0.001624538 | 0.7388912 | Cg_12A_118 | putative DNA mismatch repair enzyme |
| M896_121150 | 0.001700381 | Inf | Cg_12A_119 | DNA helicase |
| M896_121160 | 0.001732804 | 3.57369103 | Cg_12A_120 | catalytic subunit A of V/A-type ATP synthase |
| M896_121170 | 0.002845385 | 1.25011929 | Cg_12A_121 | cyclin-dependent protein kinase |
| M896_121180 | 0.00462963 | 0.30255005 | Cg_12A_122 | hypothetical protein |
| M896_121190 | 0.002992258 | 1.97115724 | Cg_12A_123 | hypothetical protein |
| M896_121200 | 0.002046784 | 0.26315789 | Cg_12A_124 | transport protein particle |
| M896_121210 | 0.003948696 | 0.45369097 | Cg_12A_125 | putative transcription factor |
| M896_121220 | 0.010525879 | 5.2402664 | Cg_12A_131 | putative ABC-like lipid transport protein |
| M896_121240 | 0.001920439 | Inf | Cg_12A_134 | hypothetical protein |
| M896_121250 | 0.001304513 | 0.08884113 | Cg_12A_135 | putative negative regulator of transcription |
| M896_121260 | 0.004615671 | 0.06037736 | Cg_12A_136 | ribosomal protein L15 |
| M896_121270 | 0 | Inf | Cg_12A_137 | subunit of polyadenylation factor I complex |
| M896_121280 | 0.001388889 | Inf | Cg_12A_138 | putative DNA-directed RNA polymerase |
| M896_121290 | 0.00246085 | 0.70921797 | Cg_12A_139 | hypothetical protein |
| M896_121300 | 0.001997009 | 0.25251005 | Cg_12A_141 | ATP binding subunit of chaperone protease |
| M896_121310 | 0.004008715 | Inf | Cg_12A_142 | hypothetical protein |
| M896_121320 | 0.000968661 | 0.10362135 | Cg_12A_143 | diphthamide biosynthesis methyltransferase |
| M896_121330 | 0.00177492 | 0.18395178 | Cg_12A_144 | hypothetical protein |
| M896_121340 | 0.002069717 | 0.41699401 | Cg_12A_146 | hypothetical protein |
| M896_121350 | 0.001385621 | 0.20084469 | Cg_12A_147 | translation elongation factor |
| M896_121360 | 0.001991239 | Inf | Cg_12A_148 | subunit of class 2 transcription repressor NC2 |
| M896_121370 | 0.001596424 | 0.47464941 | Cg_12A_149 | hypothetical protein |
| M896_121380 | 0.001315671 | 0.93193639 | Cg_12A_150 | hypothetical protein |
| M896_121390 | 0.002042484 | 0.55665796 | Cg_12A_151 | NIMA-like serine/threonine kinase |
| M896_121400 | 0.004264871 | 0.09979425 | Cg_12A_152 | putative inorganic ion transport protein |
| M896_121410 | 0.004585538 | 0.24221886 | Cg_12A_153 | putative S-adenosylmethionine-dependent |
| M896_121420 | 0.002516619 | 0.38375463 | Cg_12A_154 | hypothetical protein |
| M896_121430 | 0.003174603 | 0.21483627 | Cg_12A_155 | putative endonuclease |
| M896_121440 | 0.006325427 | 4.3977938 | Cg_12A_156 | subunit of t-SNARE complex |
| M896_121450 | 0.004196104 | 0.30272782 | Cg_12A_157 | hypothetical protein |
| M896_121460 | 0.000410142 | Inf | Cg_12A_158 | ATPase domain 1 of RNase L inhibitor |
| M896_121470 | 0.00329554 | Inf | Cg_12A_159 | putative spindle pole body associated protein |
| M896_121480 | 0.002030178 | Inf | Cg_12A_160 | hypothetical protein |
| M896_121490 | 0.001583468 | Inf | Cg_12A_161 | tRNA pseudouridine synthase domain-containing protein |
| M896_121500 | 0.001196581 | Inf | Cg_12A_162 | ribosomal protein S19 |
| M896_121510 | 0.003740374 | 0.4699409 | Cg_12A_163 | hypothetical protein |
| M896_121520 | 0.002116402 | Inf | Cg_12A_164 | inorganic phosphate transport protein |
| M896_121530 | 0.001506591 | Inf | Cg_12A_165 | hypothetical protein |
| M896_121540 | 0.001646091 | Inf | Cg_12A_166 | transcriptional repressor |
| M896_121550 | 0.002048131 | Inf | Cg_12A_167 | putative proteasome |
| M896_121560 | 0.003423592 | Inf | Cg_12A_169 | hypothetical protein |
| M896_121570 | 0.000614086 | Inf | Cg_12A_170 | hypothetical protein |
| M896_121580 | 0.000798838 | 0.67485667 | Cg_12A_171 | putative GTPase-activating protein |
| M896_121590 | 0.003866504 | Inf | Cg_12A_172 | hypothetical protein |
| M896_121600 | 0.003047619 | 0.33211168 | Cg_12A_173 | hypothetical protein |
| M896_121610 | 0.002827255 | Inf | Cg_12A_174 | hypothetical protein |
| M896_121620 | 0.001666667 | 0.82879954 | Cg_12A_175 | RNA binding repeat domain-containing protein |
| M896_121630 | 0.000932923 | Inf | Cg_12A_176 | putative amino acid permease |
| M896_121640 | 0.000814147 | 0 | Cg_12A_177 | hypothetical protein |
| M896_121650 | 0.00179937 | Inf | Cg_12A_178 | importin beta binding domain-containing protein |
| M896_121660 | 0.002126879 | 0.90078979 | Cg_12A_179 | cysteine desulfurase/transaminase |
| M896_121670 | 0.000586029 | Inf | Cg_12A_180 | UDP-N-acetylglucosamine pyrophosphorylase |
| M896_121680 | 0.002536783 | 0.54014371 | Cg_12A_181 | Man1-Src1p-C-terminal domain-containing protein |
| M896_121690 | 0.00210114 | Inf | Cg_12A_182 | hypothetical protein |
| M896_121700 | 0.001804589 | 0.62087665 | Cg_12A_183 | hypothetical protein |
| M896_121710 | 0.001809627 | 0.49452658 | Cg_12A_184 | putative heat shock protein |
| M896_121720 | 0.000877655 | Inf | Cg_12A_185 | hypothetical protein |
| M896_121730 | 0.001041667 | 0 | Cg_12A_186 | hypothetical protein |
| M896_121740 | 0.00339307 | 0.43573366 | Cg_12A_187 | hypothetical protein |
| M896_121750 | 0.002502503 | 0.55433393 | Cg_12A_188 | putative major facilitator superfamily protein |
| M896_121760 | 0.001403687 | Inf | Cg_12A_189 | putative major facilitator superfamily permease |
| M896_121770 | 0.003003003 | Inf | Cg_12A_190 | hypothetical protein |
| M896_121780 | 0.002223876 | 0.36137889 | Cg_12A_191 | hypothetical protein |
| M896_121790 | 0.002187227 | Inf | Cg_12A_192 | brix domain-containing protein |
| M896_121800 | 0.004221636 | 0.53579016 | Cg_12A_193 | putative membrane protein |
| M896_121810 | 0.002602977 | 1.16663131 | Cg_12A_194 | SPT5-like transcription initiation protein |
| M896_121820 | 0.001023123 | Inf | Cg_12A_196 | brix domain-containing protein |
| M896_121830 | 0.001868738 | 0.81377448 | Cg_12A_197 | septin-like protein |
| M896_121840 | 0.00141792 | 0.18511605 | Cg_12A_198 | hypothetical protein |
| M896_121850 | 0.001191997 | 0.28571429 | Cg_12A_199 | hypothetical protein |
| M896_121860 | 0 | Inf | Cg_12A_200 | putative ubiquitin-protein ligase |
| M896_121870 | 0.00318287 | 1.27045765 | Cg_12A_201 | SMC N-terminal domain-containing protein |
| M896_121880 | 0.000948148 | Inf | Cg_12A_202 | hypothetical protein |
| M896_121890 | 0.001141975 | Inf | Cg_12A_203 | kinesin motor domain-containing protein |
| M896_121900 | 0.000532141 | 0 | Cg_12A_204 | STE-like transcription factor |
| M896_121910 | 0.002043423 | Inf | Cg_12A_205 | hypothetical protein |
| M896_121920 | 0.001165981 | Inf | Cg_12A_206 | hypothetical protein |
| M896_121930 | 0.001674471 | Inf | Cg_12A_208 | hypothetical protein |
| M896_121940 | 0 | Inf | Cg_12A_209 | ribosomal protein L35 |
| M896_121950 | 0.004938272 | Inf | Cg_12A_210 | LSM domain-containing protein |
| M896_140020 | 0.000967098 | 1.26529296 | Cg_14_3 | aminopeptidase N |
| M896_140030 | 0.001842818 | 0.19007581 | Cg_14_4 | hypothetical protein |
| M896_140040 | 0.00514353 | 2.12206027 | Cg_14_6 | hypothetical protein |
